# ALS/FTD-linked TDP-43 alterations prevent HSV-1 infection by disrupting cell-adhesion pathways

**DOI:** 10.64898/2026.09.28.755069

**Authors:** Stephanie Jackvony Infurna, Laraib Ijaz, Evelyn J. Hardin, Abulimiti Aikedan, Constance Zhou, Tejabhiram Yadavalli, Minwoo Wendy Jang, Matthew Mahoney, Li Gan, Deepak Shukla, Adam L. Orr, Anna G. Orr

## Abstract

TAR DNA-binding protein-43 (TDP-43) alterations are a key hallmark of amyotrophic lateral sclerosis (ALS), frontotemporal dementia (FTD), and other neurodegenerative disorders. Increasing evidence links these conditions to viral infections, but it is not known if and how viral pathogens may be affected by disease-associated TDP-43. We found that ALS/FTD-linked forms of human TDP-43 prevent infection by herpes simplex virus type-1 (HSV-1), a common neurotropic virus. This cell-autonomous protective effect was conserved across different TDP-43 mutations, sporadic ALS, and *C9ORF72*-ALS, in neural and non-neural cell types, and in patient-derived cells and animal models. In addition to restricting infection, TDP-43 mutation prevented inflammatory and cell stress-related responses to HSV-1 exposure. TDP-43 knockdown but not overexpression was sufficient to protect cells from HSV-1 infection, implicating a loss-of-function mechanism. Neurons with TDP-43 mutation had disruptions in adhesion-related pathways and, accordingly, had reduced early-stage viral binding and entry. Further analyses revealed alternative RNA splicing and impairment of focal adhesion kinase (FAK/PTK2), a key mediator of cell-adhesion pathways. Inhibition of FAK was sufficient to decrease HSV-1 infection, and genetic enhancement of functional but not inactive FAK promoted infectivity in cells with TDP-43 mutation. Together, these data indicate that TDP-43 affects virus-host interactions by influencing adhesion pathways required for viral infection.

## Introduction

Transactive response region (TAR) DNA binding protein-43 (TDP-43) is a ubiquitous nucleic acid-binding protein that is expressed by most cell types, including neuronal, glial, and non-neural cells in the brain and periphery. In addition to its N- and C-terminal domains, TDP-43 protein contains two RNA-recognition motifs and a nuclear localization signal. TDP-43 is predominantly localized in the nucleus and has crucial roles in RNA splicing and transcriptional regulation. However, TDP-43 can also be found in extranuclear sites in the cell and affect stress granule formation, axonal transport, and other intracellular processes.

TDP-43 is strongly implicated in neurodegenerative disorders and serves as the main component of pathological aggregates in amyotrophic lateral sclerosis (ALS), with TDP-43 alterations present in over 95% of ALS cases.^1^ TDP-43 pathology is also prevalent in frontotemporal dementia (FTD) and other neurodegenerative disorders, including Alzheimer’s disease (AD) and limbic-predominant age-related TDP-43 encephalopathy (LATE).^1–9^ Over 70 mutations in *TARDBP* (the gene encoding TDP-43) are causatively linked to ALS/FTD.^10–12^ Most disease-associated mutations in TDP-43, including Q331K, M337V, and N352S, occur in the glycine-rich C-terminal domain. Mutations in this region can promote TDP-43 aggregation and toxicity,^12^ and affect RNA splicing and neuroimmune signaling, potentially independent of aggregation.^13,14^ Mutations in other genes, including hexanucleotide repeat expansions in *C9ORF72*, also cause TDP-43 pathology.^15^ Although *C9ORF72* expansions are the most common genetic cause of ALS/FTD, most cases with TDP-43 pathology are sporadic with no known family history or associated genetic mutations.^16,17^ The specific effects of disease-associated TDP-43 alterations on various cell functions in health and disease remain unclear. Notably, TDP-43 alterations can influence immune signaling mechanisms,^14,18–22^ and knockdown or overexpression of TDP-43 can modulate viral infections.^22–24^ However, it is not known if and how ALS/FTD- associated TDP-43 mutants and sporadic changes might influence viral infections.

Herpes simplex virus type 1 (HSV-1) is a prevalent neurotropic virus present in over 65% of the population worldwide^25^ and accounts for up to 75% of viral encephalitis cases. HSV-1 first infects peripheral epithelial cells and innervating sensory neurons, and can retrogradely transport to the brainstem and connected neocortical regions.^26–28^ Although neurons and other neural cell types have intrinsic immune responses that can be engaged to defend against pathogens,^29–33^ neurons are particularly vulnerable to neurotropic viruses, including HSV-1, due in part to their limited renewal capacity and relative separation from peripheral immune cascades. Moreover, HSV-1 has been linked to pathological hallmarks of neurodegenerative disease, including amyloid- β accumulation and tau hyperphosphorylation.^28,34–36^ The specific factors that determine individual susceptibility to HSV-1 infection may be crucial to infectious disease and neurological conditions.^37,38^ Despite the known connections between TDP-43 dysregulation and antiviral signaling, and mounting evidence linking viral pathogens to neurodegenerative disease,^36,39^ the effects of human disease-associated TDP-43 on neurotropic viral pathogens such as HSV-1 have not been explored.

Here, we investigated the effects of human TDP-43 alterations on cell-intrinsic susceptibility to HSV-1 infection using primary mouse neural cells, human iPSC-derived neural cells, patient- derived fibroblasts, and transgenic mice, and assessed the mechanisms by which TDP-43 influences virus-host interactions. We show that TDP-43 dysfunction confers resistance to HSV-1 infection. This effect is independent of neuroinflammatory and stress-linked mechanisms of antiviral defense and is instead associated with disruptions in host factors required for viral binding and entry, which protected cells from HSV-1 invasion and resulting neuropathological cascades.Complimentary genetic manipulations involving TDP-43 knockdown and overexpression systems revealed that loss-of-function in TDP-43 is sufficient to suppress HSV-1 infectivity. TDP-43 dysfunction impaired the expression of multiple cell-surface and adhesion factors. Focal adhesion kinase (FAK), a key mediator of cell-adhesion signaling pathways, was alternatively spliced and had reduced phosphorylation in neurons with mutant TDP-43. Pharmacological modulation of FAK activity was sufficient to alter cell resistance to HSV-1, and overexpression of a functionally intact FAK but not an inactive mutant variant of FAK promoted HSV-1 infectivity in cells with TDP-43 dysfunction, suggesting that FAK causally links TDP-43 impairments to changes in viral infection.

Together, our results indicate that TDP-43 is a key determinant of virus-host interactions and its dysfunction influences HSV-1 binding and entry. In particular, TDP-43 affected cell-intrinsic susceptibility to viral infection through disruptions in cellular adhesion pathways. These findings suggest that TDP-43 alterations, which are prevalent in neurodegenerative disorders, regulate cell resistance to infectious pathogens and may impact cellular functions through its effects on adhesion mechanisms.

## Results

### Disease-associated TDP-43 alterations restrict HSV-1 infection in isolated cells

HSV-1 is highly neurotropic and has direct access to neurons from peripheral epithelial cells. Thus, we first assessed whether human variants of TDP-43 can alter the susceptibility of neurons to acute infection by HSV-1. For these experiments, we first isolated postnatal mouse neurons derived from transgenic mice overexpressing human TDP-43 with the ALS-associated Q331K mutation regulated by the prion promoter,^13^ which is active in neurons and other cell types, and from littermate nontransgenic (NTG) mice expressing endogenous mouse TDP-43. The primary cultures were confirmed to be highly enriched for neurons (∼95%; **Fig. S1A**). Pilot experiments enabled selection of the appropriate viral titer (multiplicity of infection (MOI) of 0.01), similar to previously used doses.^22^ Neurons were exposed to HSV-1 strain H129-772,^40^ which was originally isolated from the brain of an individual with viral encephalitis and engineered to express an eGFP reporter in infected cells. After 72 h, virus-specific DNA encoding the envelope protein glycoprotein B (gB) was measured by quantitative PCR (qPCR). gB is essential for viral entry and conserved across all herpesviruses,^41^ and is therefore a robust measure of HSV-1 infection. Specificity of the qPCR assay was confirmed by the lack of detectable gB in noninfected cells (not shown). Strikingly, we found that neurons expressing TDP-43 with the Q331K mutation had minimal levels of viral DNA (gB) as compared to control neurons expressing endogenous mouse TDP-43 (**Fig. 1A**), suggesting that TDP-43 mutation causes a cell-autonomous decrease in HSV-1 infection in mouse neurons.

**Fig. 1.**
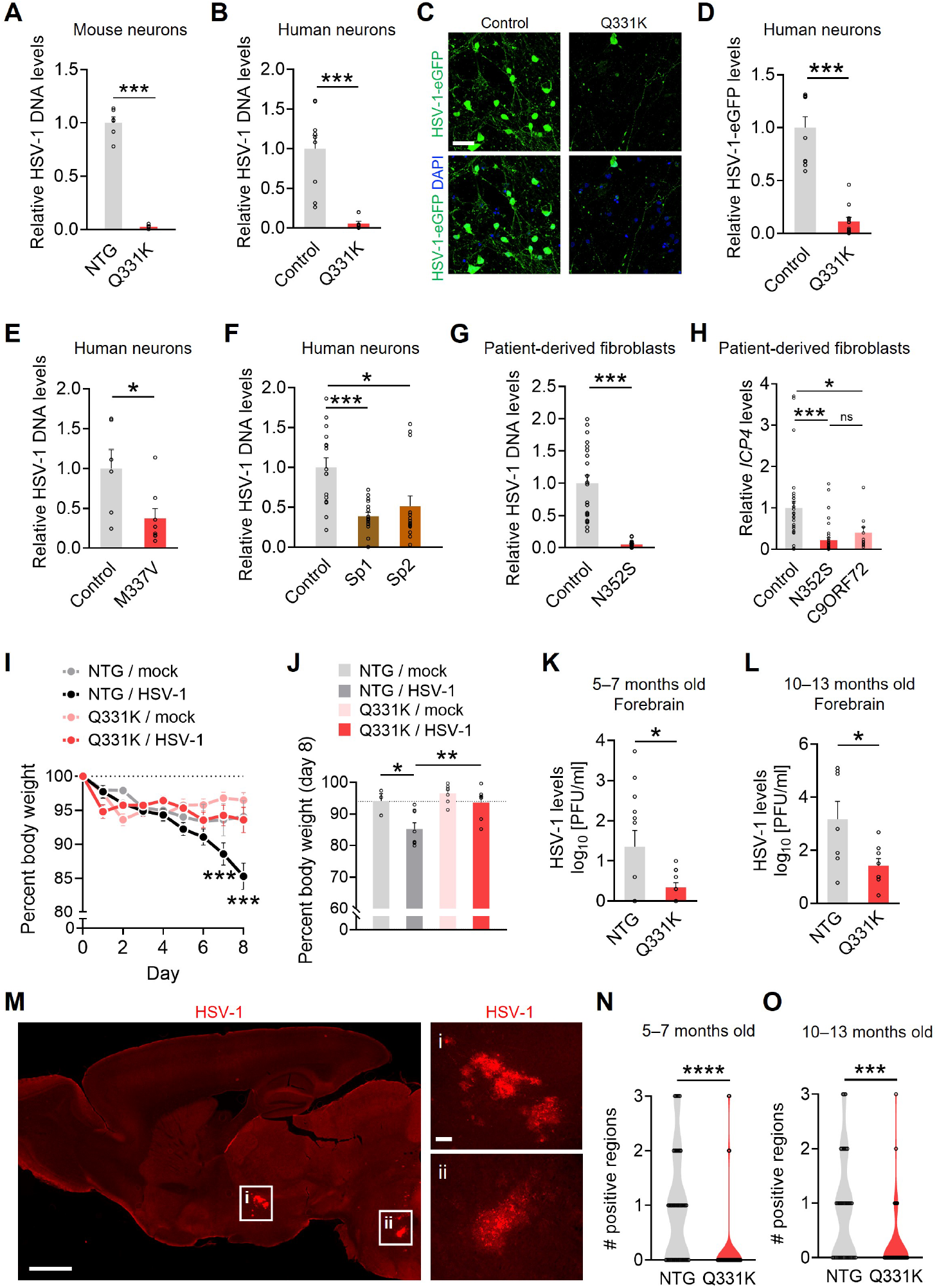
TDP-43 alterations restrict HSV-1 infection in isolated cells and *in vivo*. (**A**) Primary mouse neurons were exposed to HSV-1-eGFP (H129HSV772, 0.01 MOI) for 72 h and DNA levels of HSV gB gene were measured by qPCR. Normalized to 18S and NTGs. n = 4–6 wells. Welch’s *t* test: ***p < 0.0005. (**B**–**D**) Human iPSC-derived neurons were exposed to HSV-1-eGFP (0.01 MOI) for 72 h and DNA levels of HSV gB gene were measured by qPCR (B) or HSV-1-eGFP levels were assessed by imaging (C, D). B: Normalized to 18S and Controls. n = 9 wells from two independent differentiations. Welch’s *t* test: ***p = 0.0005. Scale bar: 40 µm. D: Normalized to MAP2 area and Control + HSV-1 group, and background corrected to noninfected controls. n = 10–12 wells from two independent differentiations, three ROIs per well. Welch’s *t* test: ***p < 0.0001. (**E**, **F**) Human iPSC-derived neurons were exposed to HSV-1-eGFP (0.01 MOI) for 72 h and DNA levels of HSV gB gene were measured by PCR. Normalized to 18S and Controls. Sp: sporadic disease cases. n = 6–16 wells from one or two independent differentiations. Student’s *t* test: *p = 0.0264 (E); Brown-Forsythe ANOVA, Dunnett’s post-hoc test: *p = 0.0290, ***p = 0.0004 (F). (**G**) Patient-derived fibroblasts were exposed to HSV-1-eGFP (0.01 MOI) for 48 h and DNA levels of HSV gB gene were measured by qPCR. Normalized to 18S and Control. n = 30–40 wells from two independent patient lines per group. Student’s *t* test, ***p < 0.0001. (**H**) Patient-derived fibroblasts were exposed to HSV-1-eGFP (0.01 MOI) for 72 h and HSV-1 immediate-early (ICP4) transcripts were measured by RT-qPCR. Normalized to *ACTB* and *TBP1* per sample and to Control. n = 10–35 wells from 3–4 separate independent patient lines per group. Welch’s ANOVA, Dunnett’s post-hoc test: *p = 0.0250, ***p = 0.0003. (**I**) Cornea of anesthetized mice were scratched using a 30-gauge needle and inoculated with 5×10^5^ pfu of McKrae strain HSV-1 or vehicle. Mice were weighed daily, and brain tissue was harvested at 8 DPI. n = 3–8, 10–13 months old. Two-way ANOVA, time factor: *F*(8, 189) = 14.88, ****p < 0.0001; *F*(24, 189) = 4.095, time x group factor: ****p < 0.0001. Dunnett’s post-hoc test: ***p < 0.001 vs. Q331K / HSV-1 at indicated timepoints. (**J**) Comparison of body weights across groups on day 8 DPI. Two-way ANOVA: *F*(1, 21) = 8.736, p = 0.0075 for HSV-1 effect; *F*(1, 21) = 7.364, p = 0.0130 for genotype effect; Bonferroni’s post-hoc test: *p = 0.0199, **p = 0.0042 vs. indicated groups. (**K**, **L**) Plaque-forming units of HSV-1 per ml detected in mouse forebrain at 5–7 months of age (K) and 10–13 months of age (L). n = 7–8 mice per genotype. Welch’s *t* test: *p < 0.05. (**M**) Immunolabeling for HSV-1 in NTG mouse. Scale bars: 600 µm, 25 µm (inset). (**N**, **O**) Quantification of HSV-1 immunolabeling as sum of focal infections across sagittal sections. n = 7–8 mice per genotype, 4–7 sections per mouse. 5–7 months old (N) and 10–13 months old (O). Mann-Whitney test: ***p = 0.0005, ****p < 0.0001.

To determine whether mutation-associated decrease in HSV-1 infection was conserved across species and conditions and not caused by TDP-43 overexpression, we examined mature human cortical neurons differentiated from patient-derived induced pluripotent stem cells (iPSCs) heterozygous for the ALS-linked Q331K mutation in TDP-43 (**Fig. S1B**).^42^ iPSC-derived neurons were used on day 35–45 of differentiation. Isogenic control neurons and Q331K neurons had similarly high expression of neuronal markers MAP2 and NeuN (**Fig. S1C**–**S1E**), with minimal detection of astrocytic marker SOX9 and microglial/macrophage marker Iba1 (**Fig. S1F** and **S1G**). Neuronal differentiation was further verified by the presence of characteristic electrophysiological properties, including spontaneous excitatory postsynaptic currents (sEPSCs), current-voltage relationships, and action potentials (**Fig. S1H**–**S1J**). There were no detectable differences in these parameters between isogenic controls and Q331K neurons. Passive membrane properties were also comparable between the groups (**Fig. S1K** and **S1L**). Additionally, transcriptional profiling of iPSC-derived neurons using RNAseq showed robust expression of various neuronal genes, including multiple subtypes of ion channels, presynaptic release machinery, neurotransmitter receptors, and postsynaptic markers (**Fig. S2**). These transcripts were also similar between isogenic controls and Q331K neurons, further indicating that neuronal differentiation and maturation were comparable across groups and that the Q331K mutation did not affect general neuronal viability and physiological properties.

We exposed human iPSC-derived neurons to HSV-1 strain H129-772 at MOI of 0.01 for 72 h, similar to mouse-derived neurons. Consistent with the results in mouse-derived primary neurons, human Q331K neurons were strongly protected from HSV-1 infection as compared to isogenic control neurons (**Fig. 1B**). In addition to neurons, human Q331K astrocytes were partially protected as compared to isogenic control astrocytes (**Fig. S3**). Fluorescence imaging further confirmed reduced levels of HSV-1 in human Q331K neurons as compared to isogenic controls (**Fig. 1C** and **1D**). To test whether protection from HSV-1 infection was specific to the Q331K mutation or extended to other disease-linked mutant forms of TDP-43, we also examined human iPSC-derived neurons heterozygous for the ALS-linked TDP-43 mutation M337V.^12,43,44^ Similar to the effects of Q331K, human neurons with the M337V mutation had reduced HSV-1 infection as compared to isogenic controls (**Fig. 1E**).

Although TDP-43 dysregulation and inclusions are the predominant pathological feature of ALS, approximately 90% of ALS cases are sporadic and without a known genetic cause.^45^ Thus, we next tested if human iPSC-derived neurons from individuals with sporadic ALS also have altered susceptibility to HSV-1. We analyzed two lines from sporadic ALS cases and an age- and sex-matched control line (**Table S1**).^46^ Neurons derived from these ALS cases have TDP-43 impairments as detected by specific changes in RNA splicing patterns.^46,47^ Similarly to cells with mutations in TDP-43, HSV-1-infected neurons from sporadic ALS cases had lower levels of HSV-1 infection as compared to neurons from control cases (**Fig. 1F**), suggesting that TDP-43 dysfunction linked to sporadic disease can also confer protection against HSV-1 infection.

In addition to neurons, HSV-1 is capable of infecting peripheral cells.^48^ We next examined a non-neural cell type by obtaining patient-derived fibroblasts from multiple independent ALS cases associated with the disease-linked N352S mutation in TDP-43 and independent control patient lines (**Table S2**). The fibroblasts were grown to similar densities and confirmed to have similar molecular features between groups (**Fig. S4A** and **S4B**). Consistent with the effects of Q331K and M337V in neurons, human N352S fibroblasts had decreases in HSV-1 levels (**Fig. 1G** and **S4C**). Further testing across time-points showed that N352S cells had consistently lower HSV-1 infection relative to control cells over multiple days after viral exposure (**Fig. S4D**), with the largest effect sizes detected at earlier timepoints (**Fig. S4E**).

We also obtained three additional independent ALS cases with disease-linked repeat expansions in the *C9ORF72* gene, which is the most common genetic cause of ALS/FTD and is associated with TDP-43 pathology^1,15,45^ (**Table S2**). *C9ORF72* expansions were associated with marked suppression of HSV-1 infection in patient-derived fibroblasts, similar to N352S fibroblasts, as detected by reduced levels of immediate-early (*ICP4*) and early (*ICP8*, *TKB)* HSV-1 RNA transcripts (**Fig. 1H**, **S4F**, and **S4G**). Together, these data reveal conserved and cell-autonomous effects of disease-associated TDP-43 dysregulation on neural and non-neural cell permissivity to HSV-1 infection.

We also examined whether TDP-43 mutation affects infection by other viruses by testing human cytomegalovirus (HCMV, HHV-5), another highly prevalent neurotropic virus in the *Herpesviridae* family that has been linked to neurodegenerative disease.^49^ Although HCMV shares some structural features with HSV-1, it has distinct tropism and clinicopathologic effects. In contrast to HSV-1 infection, we did not detect protection against HCMV in either human iPSC- derived neurons or patient-derived fibroblasts expressing TDP-43 with either Q331K or N352S mutations, respectively (**Fig. S5A** and **S5B**). Instead, TDP-43 mutations were associated with marked increases in HCMV levels in both models, revealing increased susceptibility to HCMV infection. These results suggest that the effects of TDP-43 mutation are virus-specific but consistent across cell types and different mutations.

### Disease-associated TDP-43 alterations restrict HSV-1 infection *in vivo*

We next investigated whether the cell-autonomous protection against HSV-1 in cultures was sufficient to reduce vulnerability to viral infection *in vivo*. For these experiments, we used corneal scarification to unilaterally infect the mouse cornea with 5 x 10^5^ plaque-forming units (PFU) of McKrae strain HSV-1 (**Fig. S6A**). The McKrae strain is a highly neuroinvasive strain purified from an individual with herpes simplex keratitis.^50,51^ Lytic virus peaks in the mouse brain at approximately 8 days post-infection (8 DPI) by corneal scarification.^26,52^ To assess the effects of TDP-43 mutation *in vivo*, we examined 5–7 and 10–13-month-old transgenic mice overexpressing human TDP-43 with the Q331K mutation under the prion promoter,^13^ which is active in neurons and other cell types. Controls included age- and sex-matched nontransgenic (NTG) littermates and mock-infected controls that received corneal scarification without virus.

NTG mice infected with HSV-1 lost approximately 15% of their body weight by eight days post-infection (**Fig. 1I** and **1J**). In contrast, HSV-1-infected Q331K mice had significantly less weight change than infected NTG mice and lost no more weight than mock-infected control mice (**Fig. 1I** and **1J**), suggesting that the Q331K mutation alleviated a primary systemic indicator of disease. Of note, baseline body weight was affected by both sex and genotype (**Fig. S6B**). In addition, female NTG mice had greater weight loss post-infection than male NTG mice (**Fig. S6C**), and Q331K females were more robustly protected from infection-related weight loss (**Fig. S6D**).

Upon infection of the corneal epithelium, HSV-1 has a direct path to the sensory neurons of the trigeminal ganglia. From the trigeminal ganglia, the virus can be retrogradely transported to the brainstem and connected forebrain areas.^48^ Accordingly, we next measured the levels of HSV-1 in the forebrain and trigeminal ganglion using the viral plaque assay to assess virus levels. We found that both age groups of Q331K mice had significantly lower levels of infectious virus in the forebrain as compared to NTG controls, with up to 465-fold lower lytic HSV-1 in Q331K mice as compared to NTG littermate controls (**Fig. 1K** and **1L**). There were no detectable sex differences in these outcomes. Similar Q331K effects were detected in the trigeminal ganglia in both age groups (**Fig. S6E** and **S6F**). Western blotting confirmed that human mutant TDP-43 protein was expressed in the trigeminal ganglia and forebrain of Q331K mice, with higher levels in the forebrain (**Fig. S6G**–**S6J**). Further analyses of HSV-1 immunoreactivity showed that Q331K mice exposed to HSV-1 had reduced occurrence of focal infection cores throughout the brain at 5–7 and 10–13 months of age as compared to NTG controls (**Fig. 1M**–**1O** and **S6K**). Together, these results indicate that ALS-linked Q331K mutation reduces brain vulnerability to peripheral HSV-1 infection.

### Peripheral HSV-1 infection induces neuroimmune responses in the brain

We next examined brain inflammatory responses to acute peripheral HSV-1 infection by assessing gene expression patterns in mice using bulk RNAseq and pathway analyses. The hippocampus is particularly vulnerable to inflammation^53,54^ and neurodegenerative processes,^55–57^ and showed HSV-1 infection in NTG but not Q331K mice (**Fig. S7A**), making it a relevant brain region for comparing the pathological effects of HSV-1. In the hippocampus, we identified 1,729 differentially expressed genes (DEGs) in HSV-1-infected NTG mice as compared to mock-infected NTG mice, with HSV-1 infection increasing most DEGs (**Fig. 2A** and **2B, Supplemental Dataset 1**). Interferome analyses^58,59^ showed that the majority of altered transcripts (71%; 1,230 of 1,729) are regulated by type I or type II interferons (IFNs), which are essential for antiviral responses (**Fig. 2B**). Ingenuity Pathway Analysis (IPA) further revealed that HSV-1 infection increased a broad array of hippocampal immune, antiviral, cell stress, and cytokine signaling-related pathways (**Fig. 2D** and **Supplemental Dataset 2**). Predictive analyses of upstream transcription factors identified CREB1, STAT1/3, NF-κB, and interferon regulatory factors (IRFs), among others (**Fig. S7B**). These findings show that acute peripheral HSV-1 exposure induces brain inflammatory responses.

**Fig. 2.**
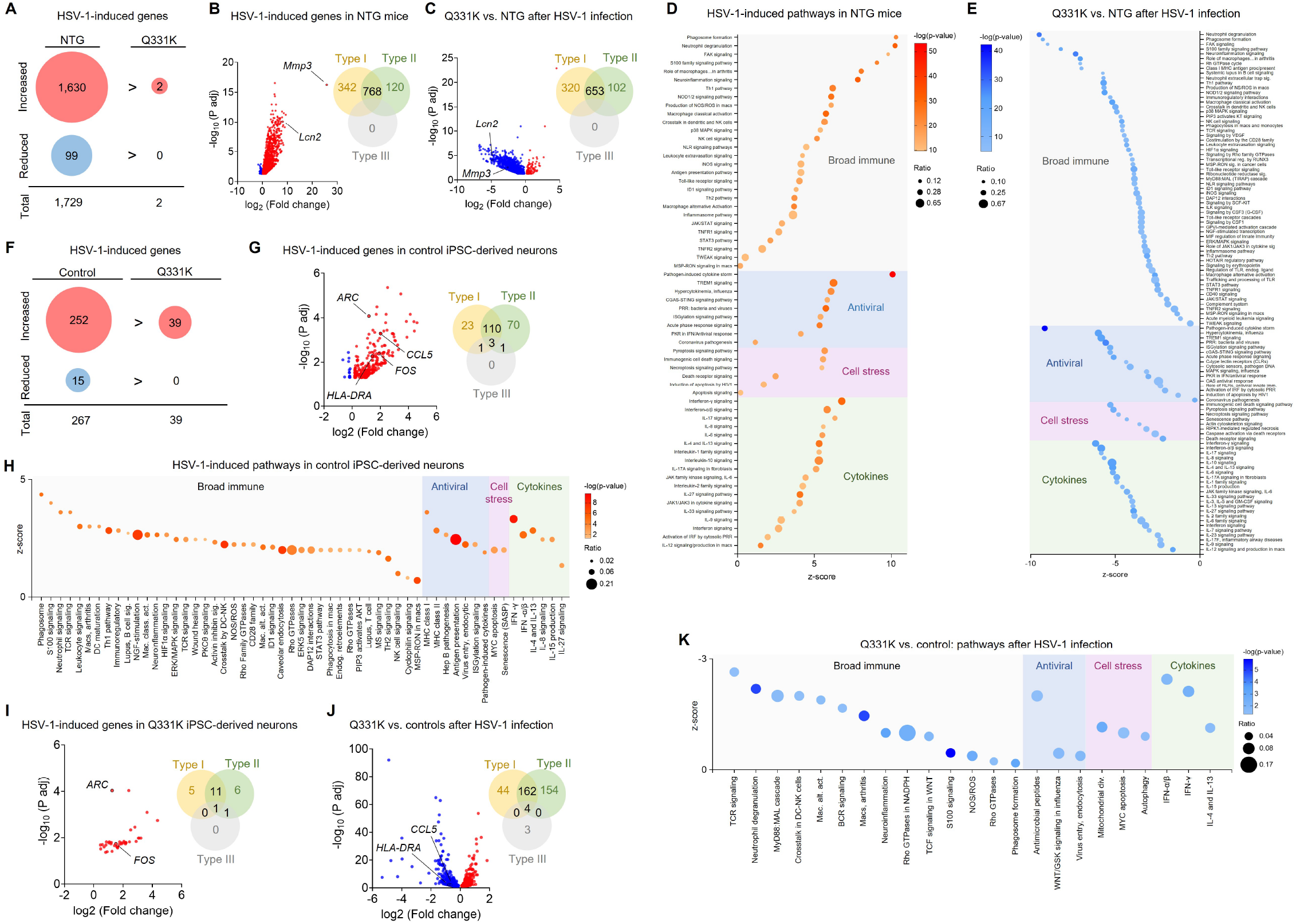
TDP-43 alterations prevent neuroimmune and cell-stress responses to HSV-1. **(A)** DEG counts in NTG and Q331K mouse hippocampi following inoculation with HSV-1 through the corneal scarification route. n = 4 mice per condition, 10–13 months old, 8 DPI. **(B)** Volcano plot of genes differentially expressed in NTG hippocampi from mice inoculated with HSV-1 via corneal scarification versus mock-infected controls. Counts of interferon-regulated genes based on Interferome database analysis. n = 4 mice per condition, 10–13 months old, 8 DPI. **(C)** Volcano plot of genes differentially expressed in hippocampi from Q331K mice inoculated with HSV-1 versus infected NTG controls. Counts of interferon-regulated genes based on Interferome database analysis. n = 4 mice per condition, 10–13 months old, 8 DPI. (**D**, **E**) Pathway analysis of DEGs in (B, C) with pathways relevant to antiviral and interferon responses plotted. **(F)** DEG counts in control and Q331K iPSC-derived human neurons following exposure to HSV-1. n = 3–6 wells per condition from one differentiation, MOI 0.01, 72 h. **(G)** Volcano plot of genes differentially expressed in control iPSC-derived human neurons exposed to HSV-1 versus vehicle controls. Counts of interferon-regulated genes based on Interferome database analysis. n = 3–6 wells per condition from one differentiation, MOI 0.01, 72 h. **(H)** Pathway analysis of DEGs in (B) with pathways relevant to antiviral and interferon responses plotted. **(I)** Volcano plot of genes differentially expressed in Q331K iPSC-derived human neurons exposed to HSV-1 versus vehicle controls. Counts of interferon-regulated genes based on Interferome database analysis. n = 3–6 wells per condition from one differentiation, MOI 0.01, 72 h. **(J)** Volcano plot of genes differentially expressed in Q331K iPSC-derived neurons exposed to HSV-1 versus isogenic controls. Counts of interferon-regulated genes based on Interferome database analysis, showing HSV-1-induced gene expression differences between genotypes. n = 3–6 wells per condition from one differentiation, MOI 0.01, 72 h. **(K)** Pathway analysis of DEGs in (J) with pathways relevant to antiviral and interferon responses plotted.

### TDP-43-linked resistance to infection does not involve enhanced neuroimmune responses

TDP-43 mutations and other disruptions are known to promote innate immune pathways and activate antiviral factors, including NF-κB, JAK/STAT, cGAS/STING, and IFN-related cascades,^14,22,60,61^ which we hypothesized could be responsible for the reduced viral susceptibility phenotype. Thus, we next examined whether restricted HSV-1 infection in the hippocampus of mice with mutant TDP-43 (**Fig. S7A**) was associated with enhanced neuroimmune responses, which might restrict infection by suppressing HSV-1 replication and/or facilitating viral clearance. In particular, we examined Q331K mice at baseline and after exposure to HSV-1 for evidence of enhanced or otherwise altered hippocampal neuroinflammatory, antiviral, and cell stress-related transcriptional signatures as compared to NTG mice.

Comparison of mock-infected Q331K to mock-infected NTG control mice revealed only 19 DEGs in the hippocampus (**Fig. S7C** and **Supplemental Dataset 1**), supporting the established regional specificity of degeneration and motor phenotypes in this model.^13,62^ Top altered genes were linked to intracellular trafficking and other functions but not antiviral defense, suggesting that the restricted HSV-1 infection in Q331K mice was not due to a preexisting antiviral gene signature or an augmented neuroinflammatory state. Moreover, comparison of HSV-1-infected Q331K mice to mock-infected Q331K controls identified only two DEGs (**Fig. 2A** and **S7D, Supplemental Dataset 1**), further ruling out the possibility of an amplified neuroimmune response upon HSV-1 infection as the primary cause of suppressed infection in Q331K mice. Indeed, additional comparisons between HSV-1-infected Q331K mice to HSV-1-infected NTG mice revealed 1,607 DEGs (**Fig. 2C** and **Supplemental Dataset 1**), most of which were reduced in Q331K mice. Pathway analyses confirmed that the majority of these changes (67%; 1,075 out of 1,607) occurred to type I or type II IFN factors (**Fig. 2C**). Infected Q331K mice had markedly lower engagement of immune, antiviral, cell stress, and cytokine pathways in comparison to infected NTG mice (**Fig. 2E** and **Supplemental Dataset 3**) and, accordingly, had lower predicted activities of various transcription factors as compared to infected NTG mice (**Fig. S7E**).

In complement to the unbiased RNA profiling, targeted RT-qPCR analyses of a curated set of well-known neuroimmune and neuropathology-related transcripts confirmed that many neuroimmune factors were increased in infected NTG mice as compared to mock-infected NTG controls but only one transcript (*Cxcl9*) was changed in infected Q331K mice (**Fig. S8A**). Likewise, immunolabeling for microglial markers in thalamic regions, which showed consistent focal infections in NTG mice (**Fig. S5K**), indicated that infected NTG mice had robust microgliosis as compared to mock-infected NTG controls but that microglia in Q331K mice exposed to HSV-1 were not different from mock-infected mice of either genotype (**Fig. S8B** and **S8C**), further ruling out enhanced neuroinflammatory responses as a mechanism mediating the reduced infection in mice with mutant TDP-43.

We next assessed whether TDP-43 mutation-associated lack of immune response to HSV-1 occurred cell-autonomously and was conserved in human cells. We performed RNAseq in human iPSC-derived Q331K and isogenic control neurons exposed to HSV-1 strain H129-772 and identified 252 upregulated and 15 downregulated genes in response to the virus (**Fig. 2F** and **Supplemental Dataset 4**). Similar to our findings in mice, human Q331K neurons had substantially fewer DEGs induced by HSV-1, with only 39 upregulated DEGs (85% reduction relative to control neurons) (**Fig. 2F** and **Supplemental Dataset 4**). Interferome analyses of DEGs in infected versus non-infected human control neurons showed that the majority of transcripts (208 of 267, 78%) were regulated by interferons (**Fig. 2G**). IPA confirmed strong upregulation of various immune, antiviral, cell stress, and cytokine signaling pathways in response to HSV-1 exposure in control neurons (**Fig. 2H** and **Supplemental Dataset 5**), and transcription factor analysis identified proinflammatory (RELA, STAT1/3) and antiviral (IRF8) transcription factors (**Fig. S9A**). In human Q331K neurons, Interferome analysis also suggested that many of the 39 DEGs (62%) in infected versus non-infected Q331K neurons are regulated by interferons (**Fig. 2I**). However, IPA did not reveal strong engagement of proinflammatory and antiviral pathways (out of the four predicted activated pathways, only one was relevant to these responses: NGF-stimulated transcription (−log (P adj) = 6.8, ratio = 0.103, z-score = 2)). In addition, the 39 DEGs predicted the activation of a partly distinct set of transcription factors in infected Q331K neurons (**Fig. S9B**). We detected 747 DEGs in infected Q331K neurons as compared to infected isogenic controls (376 upregulated and 371 downregulated) (**Fig. 2J** and **Supplemental Dataset 4**). After correcting for baseline genotype differences, 507 genes were uniquely altered with HSV-1 infection, with 72% of these genes regulated by interferons (**Fig. 2J**). IPA showed reduced induction of proinflammatory and antiviral pathways and reduced activities of predicted proinflammatory transcription factors in HSV-1- exposed Q331K neurons as compared to infected isogenic controls (**Fig. 2K** and **S9C**, and **Supplemental Dataset 6**).

These results further refute the possibility that TDP-43 mutation reduces HSV-1 infection by promoting innate immune and antiviral responses, which can limit replication or promote clearance of virus, and instead suggest that TDP-43 mutation restricts infection through cell-autonomous mechanisms positioned upstream of immune responses.

### TDP-43 alterations are bidirectionally linked to HSV-1 infection

Viral pathogens, cell stress, and neuroinflammation can alter TDP-43 protein homeostasis.^63–65^ To assess if HSV-1 infection influences TDP-43 homeostasis, we performed immunohistochemical analyses of subcellular localization of TDP-43 *in vivo*. We found that the thalamic region of NTG mice infected with HSV-1 had consistent increases in extranuclear TDP-43 immunoreactivity as compared to mock-infected controls, with no significant changes in total nuclear TDP-43 immunoreactivity levels (**Fig. S10A** and **S10B**). Infected NTG mice also had prominent TDP-43- immunopositive puncta reminiscent of protein aggregates (**Fig. S10A**). In contrast, TDP-43 had normal subcellular localization in Q331K mice exposed to HSV-1, with overall patterns similar to mock-infected NTG and Q331K controls and no evidence of TDP-43 aggregation (**Fig. S10A** and **S10B**). Consistent with previous findings,^13^ there were no baseline increases in extranuclear levels of TDP-43 in Q331K mice as compared to NTG controls (**Fig. S10A** and **S10B**).

To test whether HSV-1 also promotes TDP-43 alterations in human neurons, we assessed subcellular localization of TDP-43 in WT and Q331K human iPSC-derived neurons (D35) after exposure to 0.01 MOI HSV-1 or vehicle for 72 h. In this acute infection paradigm, HSV-1 exposure induced nuclear TDP-43 aggregates in control human neurons but had minimal effects on TDP-43 in isogenic Q331K neurons (**Fig. S10C** and **S10D**). HSV-1 did not affect overall neuronal densities in Q331K or isogenic control neurons at 72 h (**Fig. S10E**), suggesting that HSV-1 and the inclusions were not acutely toxic. These results are consistent with reports that iPSC-derived neurons survive at least eight days after HSV-1 infection at high MOI,^66^ and that nuclear TDP-43 inclusions might have protective effects in the context of acute cell stress.^67^ Together with the *in vivo* evidence, these results suggest that acute HSV-1 infection promotes TDP-43 inclusions, but these effects are minimized by TDP-43 mutation, likely due to the resistance of mutant cells to initial HSV-1 infection.

In addition to changes in TDP-43, acute HSV-1 infection may nonspecifically disrupt nucleocytoplasmic protein transport and thereby affect general protein homeostasis. To test this possibility, we transfected HEK293T cells with a construct encoding tdTomato-NES-NLS (tdT-NES-NLS), a fluorescent protein containing a nuclear localization signal (NLS) and a nuclear export signal (NES) that together enable bidirectional transport between the nucleus and cytoplasm and allow for assessment of nucleocytoplasmic flux across conditions.^68,69^ Exposure of the cells to HSV-1 for 48 h was sufficient to increase cytoplasmic-to-nuclear ratios of tdT-NES-NLS relative to noninfected controls (**Fig. S10F** and **S10G**), suggesting that HSV-1 nonspecifically disrupts nucleocytoplasmic protein translocation, with broad implications for proteinopathies.

These findings suggest that HSV-1 and TDP-43 homeostasis may have bidirectional functional connections. In particular, HSV-1 infection can induce changes in TDP-43 and related proteostasis mechanisms, and alterations in TDP-43 can in turn influence HSV-1 infectivity and associated pathogenic processes.

### TDP-43 alterations disrupt cell expression of viral binding and entry factors

Our results in isolated cells and mice suggest that TDP-43 mutation does not protect from HSV-1 infection by enhancing neuroimmune or cell stress-related responses. Instead, they point to effects on the initial stages of viral infection, namely viral binding and entry into cells, which precede virus-induced immune responses. To explore the upstream molecular mechanisms that affect HSV-1 infection, we next performed RNAseq analyses in non-infected human iPSC-derived cortical neurons heterozygous for the Q331K mutation and isogenic controls. In total, we detected 2,369 DEGs between Q331K and isogenic control cells at baseline, with most of these transcripts reduced by Q331K neurons (1,857, 78%) (**Fig. 3A** and **Supplemental Dataset 7**). Of the 2,369 DEGs, 568 (24%) overlapped with DEGs reported in human iPSC-derived motor neurons from ALS cases with other TDP-43 mutations,^70^ revealing consistencies between iPSC lines, neuronal differentiation procedures, and TDP-43 variants.

**Fig. 3.**
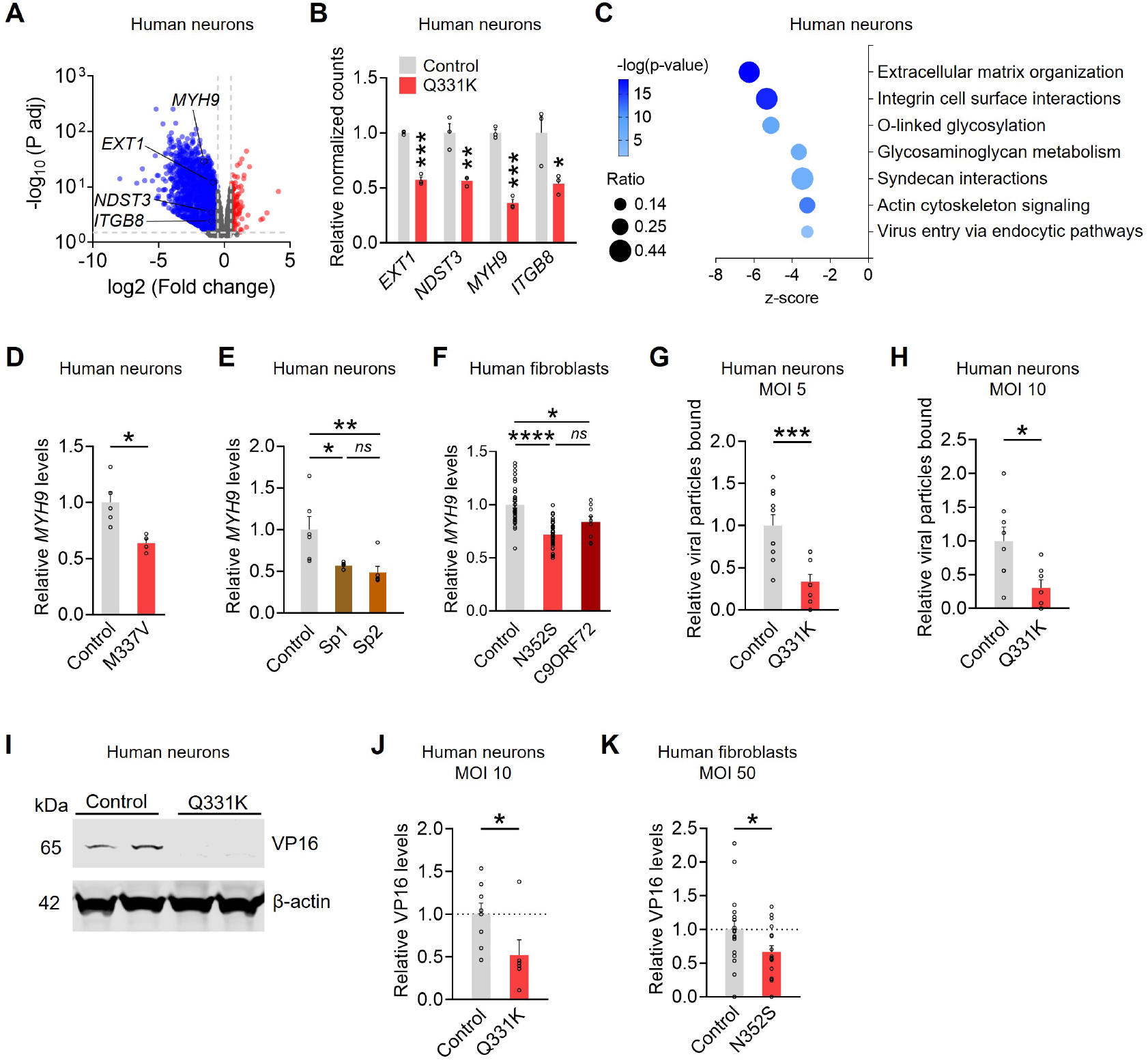
TDP-43 alterations disrupt cell-surface binding and entry by HSV-1. **(A)** Volcano plot of genes differentially expressed in iPSC-derived Q331K human neurons as compared to isogenic controls. n = 3 wells from one differentiation. **(B)** Relative normalized counts for genes involved in host HSV-1 receptor expression in iPSC-derived neurons. Normalized to controls. n = 3 wells from one differentiation. Student’s *t* test, *p = 0.0443, **p = 0.0078, ***p < 0.0005. **(C)** Pathway analysis of DEGs in (A) with pathways relevant to cell-surface and viral entry plotted. (**D**, **E**) RT-qPCR analysis of *MYH9* transcripts in iPSC-derived neurons. Normalized to *ACTB* and *TBP1* per sample and to controls. n = 4–6 wells from one differentiation. Student’s *t* test: *p = 0.0143 (D); n = 6 wells. one-way ANOVA, Tukey’s test: *p = 0.0206, **p = 0.0066 (E). (**F**) RT-qPCR analysis of *MYH9* transcripts in patient-derived fibroblasts. Normalized to *ACTB* and *TBP1* per sample and to controls. n = 9–35 wells from 3–4 individual patient lines. One-way ANOVA, Tukey’s test: *p = 0.0186, ****p < 0.0001. (**G**, **H**) Human iPSC-derived Q331K and isogenic control neurons were exposed to cold HSV-1- eGFP at an MOI of 5 (G) or 10 (H) for 30 min on ice. Cells were fixed and immunolabeled for HSV-1. Punctate viral particles were quantified. Normalized to Control and background-corrected to noninfected (vehicle) controls. n = 7–10 wells from two independent differentiations. Welch’s *t* test: *p=0.0144, ***p=0.0006. (**I**, **J**) Human iPSC-derived Q331K and isogenic control neurons were exposed to cold HSV-1- eGFP at an MOI 10 for 30 min on ice, followed by a 15-min incubation at 37°C. Representative Western blot (I) and quantification (J) of relative levels of HSV-1 tegument protein, VP16. Normalized to β-actin per sample and to the control + HSV group, and background-corrected to noninfected (vehicle) controls. n = 6–10 wells from two independent differentiations. Student’s *t* test: *p = 0.046. (**K**) Patient-derived N352S and control fibroblasts were exposed to cold McKrae strain HSV-1 at an MOI 50 for 1 h on ice, followed by a 30-min incubation at 37°C. Immunolabeling for HSV-1 VP16 particles was quantified and normalized to vimentin area and to the control + HSV-1 group. n = 17 wells from two independent patient lines per group. Student’s *t* test: *p = 0.049.

Notably, we found that multiple cell-surface factors and host receptors for HSV-1 were reduced in Q331K neurons as compared to isogenic controls, including factors required for the production of heparan sulfate (*EXT1* and *NDST3*), which serves as a binding and entry receptor for HSV-1-gB/gC,^71–73^ non-muscle myosin heavy chain IIA (NMHC-IIA, *MYH9*), which serves as an HSV-1-gB entry receptor,^74^ and the HSV-1-gH/gL receptor integrin-β8 (*ITGB8*)^75^ (**Fig. 3B**). Indeed, among the various transcriptional changes in Q331K neurons, IPA revealed strong reductions in viral entry pathways, glycosaminoglycan homeostasis (which includes heparan sulfates), extracellular matrix (ECM) organization (including collagens and syndecans), and actin cytoskeleton signaling (**Fig. 3C**). *MYH9* was confirmed through targeted RT-qPCR to have reduced expression levels across different ALS lines as compared to their respective controls (**Fig. 3D**–**3F**). These transcriptional changes further suggest that HSV-1 infection may be restricted at the initial stages of viral binding and entry, consistent with the low levels of virus-induced inflammatory and stress-related signatures in mutant mice and human cells (**Fig. 2**).

### TDP-43 alterations disrupt cell-surface binding and entry by HSV-1

Given the observed reductions in cell-surface and adhesion factors that represent host machinery for HSV-1 binding and entry, we next tested whether the early stages of HSV-1 infection were disrupted by quantifying the number of viral particles bound to neuronal surfaces after 30 min of HSV-1 exposure. Neurons were exposed to cold HSV-1 inoculum on ice to allow for viral binding but not entry into cells.^76^ Neurons were assessed for cell-surface HSV-1 levels by immunolabeling. Across MOIs, Q331K neurons had less HSV-1 bound relative to isogenic control neurons (**Fig. 3G**, **3H**, and **S11A**) with comparable cell densities (**Fig. S11B** and **S11C**). Minimal background fluorescence was detected in non-infected cells at similar densities (**Fig. S11D** and **S11E**).

After initial binding by HSV-1, various host factors, such as 3-OS-heparan sulfate and NMHC-IIA, facilitate the coordinated fusion and entry of HSV-1 into the host cell.^72,74^ We next tested if viral entry was also disrupted in Q331K neurons by measuring the levels of viral tegument protein, VP16, following a brief infection. Neurons were exposed to HSV-1 on ice with cold inoculum (MOI 10) for 30 min, followed by an additional 15-min incubation at 37°C to allow for synchronized viral entry.^77^ VP16 protein is part of the HSV-1 tegument, and levels of VP16 detected after 15 min is proportional to HSV-1 entry. Immunoblotting for VP16 in cell lysates revealed decreased VP16 levels in Q331K neurons as compared to isogenic controls (**Fig. 3I** and **3J**), further indicating that initial viral interactions with neurons are reduced by the Q331K mutation.

To determine whether the effect on viral entry extends to other cell types and TDP-43 mutations, we next tested patient-derived fibroblasts with the N352S mutation. Human fibroblasts were exposed to HSV-1 on ice with cold inoculum (MOI 50) for 1 h, followed by an additional 30-min incubation at 37°C. Immunolabeling revealed decreased VP16 levels in N352S fibroblasts as compared to controls (**Fig. 3K** and **S11F**). Minimal background fluorescence was detected in non-infected cells at similar densities (**Fig. S11G**). Together, these data indicate that reduced HSV-1 entry is a conserved effect across cell types and TDP-43 mutations.

### TDP-43 alterations modulate HSV-1 infection by impairing FAK function

Previous studies indicate that TDP-43 pathology causes cell impairments at least in part through TDP-43 loss-of-function effects,^78^ and that mutations in TDP-43 can cause a combination of loss-of-function and gain-of-function phenotypes.^13^ We next tested whether induction of loss-of-function or gain-of-function in TDP-43 is sufficient to recapitulate cell-intrinsic resistance to HSV-1 infection. For these experiments, we induced knockdown or overexpression of wild-type human TDP-43 in HEK293T cells. Transfection with a plasmid encoding an shRNA sequence targeting the TDP-43 gene (*TARDBP*) resulted in approximately 65% reduction in *TARDBP* RNA levels and 50% reduction in TDP-43 protein levels by 72 h, whereas transfection with a nontargeting control sequence had no effect relative to untransfected controls (**Fig. S11H** and **S11I**). We found that partial knockdown of TDP-43 was sufficient to confer protection against HSV-1 infection (**Fig. 4A**), implicating a loss-of-function mechanism. In contrast, acute overexpression of TDP-43 by approximately six-fold above endogenous levels had minimal effect on HSV-1 infection as compared to cells that were either untransfected or received a control plasmid (**Fig. 4B** and **Fig. S11J**). Transient knockdown of TDP-43 was also sufficient to initiate reductions in *MYH9 and ITGB8* RNA expression (**Fig. S11K**), two host factors reduced by TDP-43 mutation and known to promote HSV-1 binding and entry, further implicating TDP-43 loss-of-function in dysregulating host genes mediating HSV-1 infectivity.

**Fig. 4.**
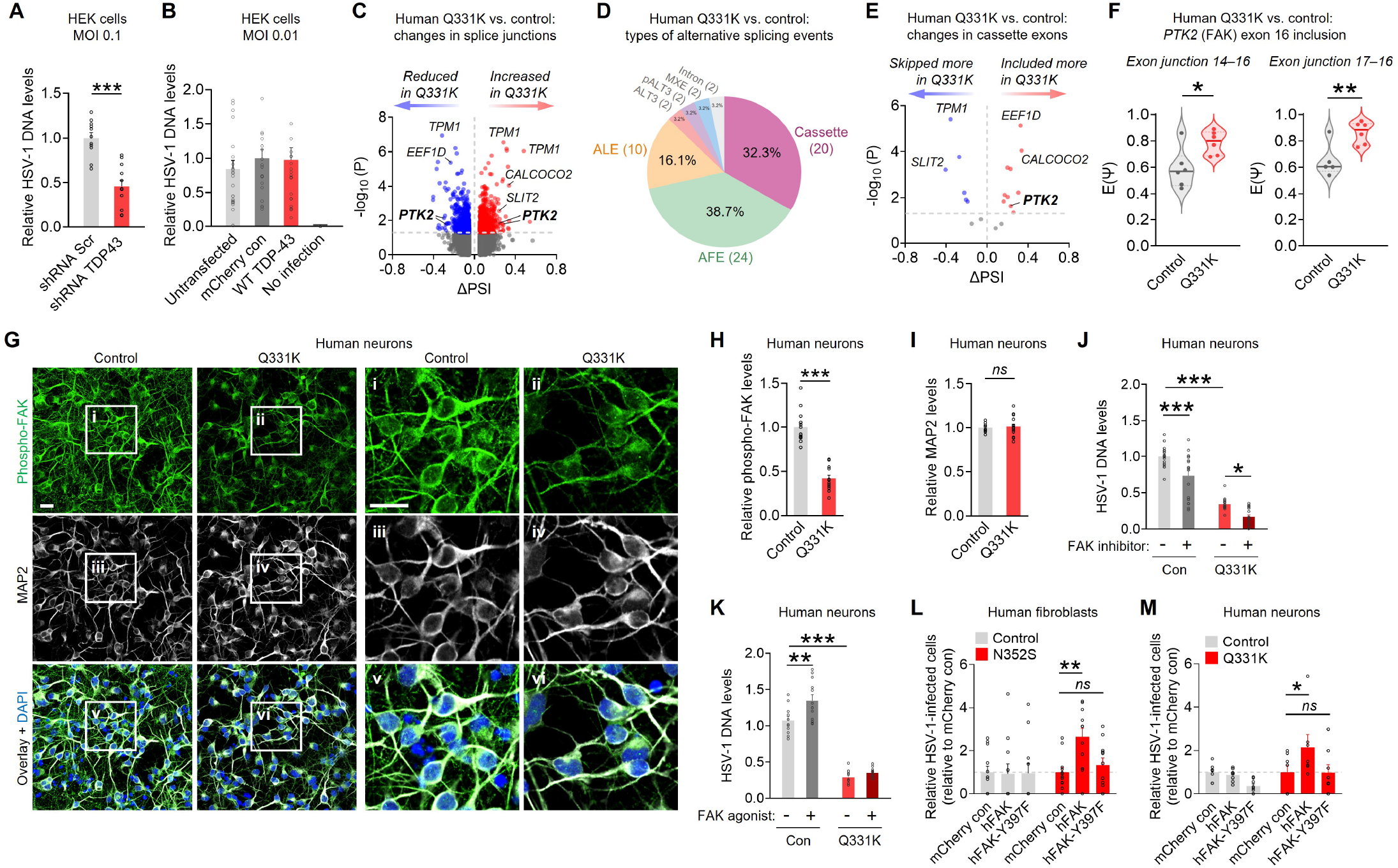
TDP-43 effects on HSV-1 infection involve impaired FAK function. **(A)** HEK293T cells were acutely transfected with indicated plasmids and exposed to HSV-1-eGFP (0.1 MOI) for 48 h. DNA levels of HSV gB gene were measured by qPCR and normalized to 18S and to nontargeting shRNA (Scr) control group. A shorter duration of viral exposure was used to avoid gene knockdown-related cell toxicity. n = 12 wells per condition from 2 independent experiments. Student’s *t* test: ***p < 0.0001. **(B)** HEK293T cells were transfected with indicated plasmids and exposed to HSV-1-eGFP (0.01 MOI) for 72 h. DNA levels of HSV gB gene were measured by qPCR and normalized to 18S RNA and to levels in the mCherry control group. n = 12–24 wells per condition from 4 independent experiments. **(C)** Volcano plot of differential splice junction usage in human Q331K iPSC-derived neurons as compared to isogenic controls. Changes in splice junction inclusion are shown as ΔPSI on the x- axis. The y-axis represents statistical significance calculated using a *t* test (−log₁₀ *P* value). Each dot corresponds to an individual splice change. Junctions with small effect sizes (|ΔPSI| < 0.05) were excluded. Splice junctions from select genes of interest, including PTK2/FAK, are labeled. n = 6 wells per genotype from one differentiation. **(D)** Pie chart summarizing the types of alternative splicing events detected in the significantly affected junctions, which were defined as passing all three statistical criteria in MAJIQ (*t* test, Tnom, and WILCOXON). Event types were classified according to MAJIQ annotations, including alternative first exon (AFE), alternative last exon (ALE), cassette exon, alternative 3′ splice (ALT3), putative alternative 3′ splice site (pALT3), mutually exclusive exons (MXE), and intron retention. The number of events in each category is indicated in parentheses. Exact splicing alterations and genomic coordinates are detailed in Fig. S12. **(E)** Volcano plot of cassette exon splicing changes in human Q331K iPSC-derived neurons as compared to isogenic controls. Cassette exon events were extracted from MAJIQ-identified significant splicing modules that passed all three statistical criteria in MAJIQ (*t* test, Tnom, and WILCOXON). Each dot corresponds to a single cassette exon. Select genes of interest, including PTK2/FAK, are labeled. Genomic coordinates of corresponding exons are detailed in Fig. S12. **(F)** Cassette exon alterations in PTK2/FAK. PSI values represent junction usage connecting the indicated exons, revealing increased inclusion of exon 16 in Q331K neurons as compared to isogenic controls (chr8, negative strand:140769575–140769592; exon 16 of transcript ENST00000696786.1, Ensembl canonical transcript). Student’s *t* test: *p = 0.02, **p = 0.009. (**G–I**) Immunolabeling (G) and quantification (H, I) of phosphorylated FAK at Y397 and neuronal marker microtubule-associated protein 2 (MAP2) in human Q331K iPSC-derived neurons and isogenic controls. Nuclei were labeled with DAPI (blue). Scale bars: 20 µm. Normalized to isogenic controls. n = 12–15 wells from two independent differentiations. Student’s *t* test: **p < 0.0001 (F), p = 0.71 (G). **(J)** Human Q331K iPSC-derived neurons and isogenic controls were treated with FAK inhibitor (PF573228, 1 µM) and exposed to HSV-1 (72 h, MOI 0.01). DNA levels of HSV gB gene were measured by qPCR and normalized to 18S levels. n = 18 wells from three independent differentiations. Two-way ANOVA: *F*(1, 68) = 25.19, p < 0.0001 for inhibitor effect; *F*(1, 68) = 199.8, p < 0.0001 for genotype effect; Bonferroni’s test: *p = 0.014, ***p < 0.0001 for indicated comparisons. **(K)** Human Q331K iPSC-derived neurons and isogenic controls were treated with FAK agonist (ZINC40099027, 1 µM) and exposed to HSV-1 (24 h, MOI 1.0). DNA levels of HSV gB gene were measured by qPCR and normalized to 18S levels. n = 11–12 wells from three independent differentiations. Two-way ANOVA: *F*(1, 43) = 4.07, p = 0.049 for interaction effect; *F*(1, 43) = 10.49, p = 0.002 for agonist effect; *F*(1, 43) = 286.0, p < 0.0001 for genotype effect; Bonferroni’s test: **p = 0.0013, ***p < 0.0001 for indicated comparisons. **(L)** Patient-derived N352S and control fibroblasts were transfected as indicated, exposed to HSV-1 (MOI 0.1, 72 h), and immunolabeled for vimentin, mCherry, and eGFP. mCherry-positive cells with HSV-1-eGFP were quantified and normalized to mCherry controls. Two-way ANOVA: *F*(2, 63) = 3.07, p = 0.053 for interaction effect; Dunnett’s test: **p = 0.0035 for indicated comparison. n = 11– 12 wells per group from two independent experiments. **(M)** Human iPSC-derived Q331K and isogenic control neurons were transfected as indicated, exposed to HSV-1 (MOI 0.01, 72 h), and immunolabeled for MAP2, mCherry, and eGFP. mCherry- positive neurons with HSV-1-eGFP were quantified and normalized to mCherry controls. Two-way ANOVA: *F*(2, 41) = 2.23, p = 0.12 for interaction effect; Dunnett’s test: **p = 0.0219 for indicated comparison. n = 7–8 wells per group from one independent experiment.

TDP-43 has important roles in RNA splicing, and TDP-43 loss-of-function is associated with alterations in splicing.^13,79,80^ Thus, we performed untargeted alternative splicing analyses using transcriptomic data collected in human iPSC-derived control and Q331K neurons analyzed with Modeling Alternative Junction Inclusion Quantification (MAJIQ, v2.5.8).^81^ Analyses of affected RNA splice junctions revealed that the Q331K mutation induced multiple changes in splicing (**Fig. 4C**). These splicing changes included diverse types of events, with the most frequent changes detected in cassette exons and alternative first or last exon usage (**Fig. 4D** and **Fig. S12**). Using stringent cutoffs to identify the most affected transcript sites (see *Methods*), we found altered cassette exon usage in a narrow set of target genes, including focal adhesion kinase (FAK/*PTK2*), a key mediator of adhesion signaling and other pathways^82^ (**Fig. 4E**). In particular, human Q331K neurons had increased retention of exon 16 in FAK/PTK2 (**Fig. 4F**), whereas total RNA levels of FAK/PTK2 were not changed relative to isogenic controls (**Supplemental Dataset 7**). Of note, the alternatively spliced *Ptk2* cassettes are conserved between mice and humans, and TDP43^Q331K^ knock-in mice have similar alternative splicing of the *Ptk2* exon in the frontal cortex,^83^ suggesting that this splicing effect occurs in adult animals.

Interestingly, HSV-2 has been previously shown to induce FAK phosphorylation at Y397 and that disruptions in FAK expression or phosphorylation at Y397 can confer protection against HSV-1 or HSV-2 infection by impairing viral entry and transport,^84^ thus linking herpesvirus infections to FAK activation. Moreover, exon 16 inclusion is located in the linker region of FAK, causing insertion of six amino acids, DEISGD, immediately after S392 and adjacent to the canonical Y397 phosphorylation site (**Fig. S13**).^85,86^ Although exact effects of this insertion are not clear, FAK Y397 can be phosphorylated through cis and trans mechanisms, with the latter involving SH2 domain-containing Src kinase and other factors that facilitate FAK activation and downstream signaling.^82^ The hexapeptide insertion near Y397 may affect autophosphorylation and/or disrupt accessibility of this site for interactions with Src and other mediators that influence FAK signaling.^85–88^ In agreement, we detected reduced levels of FAK phosphorylation at Y397 in human iPSC-derived Q331K neurons as compared to isogenic controls (**Fig. 4G** and **4H**), without changes in MAP2 levels (**Fig. 4I**), suggesting that mutant TDP-43 influences neuronal FAK activity.

It is not known if changes in FAK activity can modulate HSV-1 infection in human neurons. Thus, we next tested the effects of inhibiting FAK activity on HSV-1 infection in iPSC-derived neurons. Treatment of Q331K and isogenic control neurons with a selective FAK inhibitor that prevents Y397 phosphorylation^89^ (PF573228, 1 µM, 72 h) was sufficient to suppress HSV-1 infection, and this effect occurred irrespective of TDP-43 mutation (**Fig. 4J**), suggesting that FAK activation facilitates HSV-1 infection in nonmutant human neurons and that FAK activity mediates some of the residual infectivity in neurons with TDP-43 mutation. In agreement, a potent and selective allosteric activator of FAK (ZINC40099027, 1 µM, 24 h) that acts on the kinase domain^90^ enhanced HSV-1 infection in nonmutant human neurons (**Fig. 4K**). However, in Q331K neurons, the FAK activator had no detectable effects on the low infection levels (**Fig. 4K**), suggesting that pharmacologically enhancing intrinsic FAK kinase activity was not sufficient to reinstate HSV-1 infection in mutant neurons. Together with aberrant splicing and deficient phosphorylation of FAK detected in mutant cells, these data suggest that FAK promotes neuronal infection by HSV-1 and that TDP-43 mutation impairs the ability of FAK to engage in this function.

To further investigate whether TDP-43 mutation prevents infection due to impairments in FAK, we next tested the effects of genetic manipulation of FAK in human cells. Patient-derived control and N352S mutant fibroblasts were transfected with normal human FAK tagged to mCherry, an inactive form of FAK (Y397F) tagged to mCherry, or mCherry alone (control) for 96 h, followed by exposure to HSV-1-eGFP (MOI 0.1, 72 h). Infectivity rates were analyzed by co-immunolabeling for vimentin, mCherry, and eGFP, and comparing the proportion of transfected (mCherry-positive) cells that were infected by HSV-1-eGFP. We found that overexpression of human FAK markedly increased HSV-1 infection in mutant but not control cells, whereas the inactive form of FAK (Y397F) did not influence infectivity (**Fig. 4L**). Similar to fibroblasts, overexpression of normal but not inactive FAK increased HSV-1 infection in human iPSC-derived Q331K neurons (**Fig. 4M**). In isogenic controls, expression of inactive FAK suppressed infection, further suggesting that active FAK is required for HSV-1 infectivity in human neurons. We conclude that TDP-43-linked disruptions in FAK contribute to the cell-intrinsic resistance against HSV-1.

Altogether, our findings reveal an important role of TDP-43 in modulating herpesvirus infection and establish that TDP-43 dysfunction influences host-virus interactions through impairments in FAK-linked adhesion pathways required for HSV-1 infection (**Fig. S14**).

## Discussion

We found that ALS-linked TDP-43 alterations confer robust protection from HSV-1 infection and thus prevent infection-related neuroinflammatory and pathological responses. TDP-43 was originally discovered for its role in limiting human immunodeficiency virus (HIV) transcription^23^ and has been further linked to viral infectivity in studies using TDP-43 knockdown or overexpression.^22,24,91^ Despite this evidence and the known dysregulation of TDP-43 in familial and sporadic neurodegenerative disorders, the effects of human disease-associated TDP-43 on viral infections have not been previously investigated. Using multiple models of disease-associated TDP-43 alterations, our study reveals a key role of TDP-43 in influencing cell-intrinsic vulnerability to HSV-1 infection through regulation of cell-adhesion pathways. We conclude that TDP-43 is a functional convergence point for neurodegenerative and infectious disease mechanisms.

Using patient-derived cells, we demonstrate that ALS-linked TDP-43 dysregulation associated with Q331K and other mutations, or sporadic cases with known TDP-43 pathology, confer protection from HSV-1 infection in neural and non-neural cell types. We also found decreased expression of host HSV-1 receptors and corresponding effects on early stages of HSV-1 infection, including decreased viral binding and entry, further revealing a novel role of TDP-43 in regulating host cell surface interaction with viral pathogens. Using mouse-derived cells and a mouse model of infection, our findings show that TDP-43-associated protection against HSV-1 is conserved across species, mutations, cell types, and TDP-43 alterations caused by various mechanisms (i.e., C9ORF72 gene expansion or sporadic disease). The complementary TDP-43 knockdown and overexpression experiments further suggest that the effects of TDP-43 alterations on susceptibility to HSV-1 are due to loss-of-function mechanisms.

Using a mouse model of viral infection, we found prominent neuroimmune responses in the hippocampal formation and thalamic region upon acute HSV-1 challenge in nonmutant mice. We also detected a bidirectional link between HSV-1 and TDP-43, given that HSV-1 infection triggered TDP-43 dysregulation reminiscent of neurodegenerative disease-associated TDP-43 alterations in control mice and human neurons. Previous studies have shown that latent HSV-1 infection and reactivation in mice promotes AD-related pathology and cognitive deficits,^92^ implicating viral infection in altered proteostasis and related mechanisms. Further underlining this relationship, our study reveals that HSV-1 can rapidly trigger neuropathogenic states, including those associated with neurodegenerative conditions, potentially altering CNS vulnerability to aging and disease. Notably, exposure to HIV, Theiler’s encephalomyelitis virus, and other viruses can induce TDP-43 dysregulation.^24,63,93–95^ Although some viruses may interact with TDP-43,^96,97^ we did not detect marked colocalization of neuronal TDP-43 with HSV-1, implicating an indirect mechanism. Whether acute virus-induced changes in TDP-43 reflect a protective or maladaptive response, and whether these changes can facilitate chronic neurodegenerative pathogenesis, requires further exploration.

In line with protection from HSV-1 infection, mice with an ALS-related TDP-43 mutation and human ALS-derived neurons had strongly reduced neuroinflammatory and pathogenic responses to HSV-1, including acute changes in TDP-43, as compared to their respective controls. Together with the restricted acute infection detected across different model systems, these results indicate that disease-related forms of TDP-43 prevent HSV-1 infection in a cell-autonomous manner and through effects on early stages of host-virus interactions that precede host induction of immune and cell-stress responses. In support, we found reduced binding and entry of HSV-1 to human neurons and fibroblasts with ALS-linked mutant forms of TDP-43 within minutes of viral exposure. In line with these results, transcriptional profiling of ALS-derived human neurons revealed decreased expression of multiple host HSV-1 binding and entry receptor systems, including *MYH9*, *EXT1, and ITGB8*, which are also altered in humans with ALS.^98–100^ Studies suggest that TDP-43 may bind to *MYH9*, *EXT1* and *ITGB8* transcripts,^101^ and that TDP-43 may interact with NMHC-IIA protein,^102^ potentially through the C-terminal domain of TDP-43.^103^ We found that *MYH9* levels were decreased across different models and mutants, indicating a conserved effect. In agreement, *MYH9* is reduced in human motor cortices in ALS and FTD with repeat expansions in the *C9ORF72* gene.^99^ However, pathway analysis also underlined broader effects of disease-associated TDP-43 on cell-surface composition and adhesion pathways.

We did not detect alternative splicing of *MYH9*, *EXT1* or *ITGB8* in human Q331K iPSC- derived neurons. Instead, our analyses revealed that splicing of other transcripts was strongly altered by the Q331K mutation, including alternative splicing of FAK/PTK2. As a key mediator, FAK transduces intracellular signaling by integrins, extracellular matrix factors, receptors, and other inputs, and promotes various structural and functional effects.^82,104^ Thus, FAK is well-positioned to broadly sense and affect host cell adhesion and related pathways that influence viral binding and entry. Notably, the restrictive effect of TDP-43 dysfunction on HSV-1 infection could be prevented by expressing functional FAK, suggesting that FAK impairment in cells with TDP-43 pathology mediates resistance to HSV-1. Indeed, previous work has shown that HSV-2 induces FAK phosphorylation at Y397 and that disruptions in FAK can confer protection against infection by impacting viral entry and transport.^84^ The effects of FAK on infectivity also extend to other viruses, including Kaposi’s sarcoma-associated herpesvirus/human herpesvirus 8 (KSHV/HHV-8)^105^ and influenza A virus.^106^ Altogether, our data using selective pharmacological and genetic FAK modulation show that FAK regulates viral infectivity in neurons and that TDP-43 dysregulation restricts HSV-1 infection at least in part through disruptions in FAK and adhesion pathways.

Although FAK has been examined mostly in cancer, an ALS-linked gene variant has been reported to induce aberrant FAK activity and impair adhesion pathways.^107^ We detected increases in FAK exon 16 inclusion and changes to FAK Y397 phosphorylation in Q331K neurons. Although the exact biophysical effects of exon inclusion on FAK phosphorylation and protein-protein interactions are not clear, the predicted hexapeptide insertion is located immediately upstream of the key phosphorylation site Y397 and might therefore affect FAK activation and signaling properties. In support, alternative splicing of FAK can affect its phosphorylation dynamics.^87^ It is possible that exon 16 inclusion alters access or efficacy of Src and other kinases, thus impacting FAK-mediated signaling. Mutant TDP-43 might also further contribute to disruptions in FAK in a feedforward manner through reductions in adhesion factors that promote FAK signaling, including integrins, which were depleted in mutant neurons. Thus, although our results point to FAK as an important factor, multiple adhesion-related mechanisms are likely contributing to the downstream functional effects of TDP-43 pathology.

It is not known if there are differences in the rates of HSV-1 positivity or the risk for herpes simplex encephalitis in familial or sporadic ALS/FTD-affected individuals. Although we found that pathological TDP-43 confers protection against acute HSV-1 infection, low levels of virus were still detectable. Based on these findings and the prevalence of HSV-1 in the general population, analyses of seropositivity for antibodies against HSV-1 might not reveal marked differences. Instead, postmortem analyses of viral components in CNS might provide insights into differential resistance to infections. Indeed, HSV-1 components have been detected in human brain in *postmortem* analyses, suggesting that the virus can reach the brain and influence neural cell function without triggering encephalitis,^27,28^ but this possibility requires further studies.^48,108^ Although we did not detect a similar protective effect against HCMV, we found that TDP-43 mutation nonetheless affected HCMV infectivity, suggesting that TDP-43 regulates cell susceptibility to multiple viral pathogens and might have broad and varied effects on infectious pathogens, a possibility that requires further investigation and may provide insights into neurological disease mechanisms. The exact effects are likely dependent on the specific host cell machinery that the pathogen requires to infect cells and how this machinery is impacted by TDP-43 alterations. Given that TDP-43 pathology is also common in AD and is a defining feature of LATE cases, TDP-43 might modulate resistance to viral infections in various conditions and contribute to inconsistent detection of viral components across individuals and study cohorts, which have not accounted for the occurrence of TDP-43 alterations and their roles in infections. Indeed, multiple other factors linked to neurodegenerative disease have been implicated in antimicrobial responses. The ALS-linked gene OPTN^109^ is associated with restriction of HSV-1 virulence,^26^ and antimicrobial processes have been described in AD pathogenesis,^28,36,110–112^ pointing to amyloid-β seeding and fibrillization as well as tau phosphorylation as innate defense mechanisms against viral infections.

Here, we reveal that TDP-43 has conserved roles in host-virus interactions, and its dysregulation confers cell-intrinsic resistance to HSV-1 infection, highlighting the complex interplay between TDP-43 function and microbial pathogens. The effects of TDP-43 alterations on cell-surface factors and adhesion pathways are likely to also promote chronic changes in cellular function and influence neuropathology independently of viral infection. These TDP-43-regulated molecular pathways may therefore contribute to pathophysiological mechanisms and might provide new therapeutic strategies to prevent infectious and neurodegenerative diseases.

## Acknowledgements

We thank A.M. Fatafta and A. Bodmer for technical assistance; G. Coronas-Samano, A. Carney, and M. Garvey for administrative assistance; C. Konrad and G. Manfredi for providing passaged human fibroblasts; M. Connors, K. Karthigeyan, and S. Permar for providing human cytomegalovirus stocks. We thank R. Schwartz and J. Anrather for advice on project execution and C. Bowen for technical advice. We thank the Weill Cornell Medicine core facilities, including the Genomics Resources Core, the Research Animal Resource Center, and the Microscopy and Imaging Core for assistance and resources. We acknowledge the Weill Cornell Medicine High-Performance Computing Core for providing computational resources and support (RRID: SCR_028969). Schematics were generated with BioRender.com.

This work was funded by the National Institutes of Health: RF1NS118569 (A.G.O.), R01EY029426 (D.S.), F31AG079616 (S.J.I.), RF1AG079557 and R01AG077899 (L.G.), NIGMS Medical Scientist Training Program grant T32GM007739 to the Weill Cornell–Rockefeller–Sloan Kettering Tri-Institutional MD-PhD Program (C.Z.), and P40OD010996 for the University of Pittsburgh Center for Neuroanatomy with Neurotropic Viruses; and the Swid Family (A.G.O).

## Author Contributions

Conceptualization: S.J.I., L.I., E.J.H., A.L.O., A.G.O.; Methodology: S.J.I., L.I., E.J.H., A.A., C.Z., T.Y., M.W.J., D.S., A.L.O., A.G.O.; Investigation: S.J.I., L.I., E.J.H., A.A., M.W.J., M.M., A.G.O., A.L.O.; Formal Analysis: S.J.I., L.I., E.J.H., A.A., M.W.J., M.M., A.G.O., A.L.O.; Visualization: S.J.I., A.L.O., A.G.O.; Funding acquisition: S.J.I., D.S., L.G., A.L.O., A.G.O.; Project administration: A.L.O., A.G.O.; Supervision: L.G., D.S., A.L.O., A.G.O.; Writing – original draft: S.J.I., A.G.O.; Writing – review and editing: all authors.

## Declaration of Interests

The authors declare no competing interests.

## Correspondence

Further information and requests for resources should be directed to lead contacts: Stephanie Jackvony Infurna and Anna G. Orr.

## Resource Availability

### Materials availability

This study generated plasmids that will be made available to academic labs upon request.

### Data and code availability

All raw data reported in this paper are available upon request from the lead contact. The raw RNAseq datasets collected in this study are included in **Supplemental Datasets 1**–**7**. Our study also analyzed previously published RNAseq datasets.^70^ Any additional data used in this paper will be shared by the lead contacts upon request. This paper does not report original code. Additional information required to reanalyze the data reported in this paper is available from the lead contacts upon request.

## Methods

### Mice

All animal experiments were conducted in accordance with guidelines set by the Institutional Animal Care and Use Committee of Weill Cornell Medicine. Mice were housed in groups of 2–5 mice per cage and maintained on a 12-h light/dark cycle with *ad libitum* access to food and water. Mice were maintained on standard laboratory chow (Purina 5053). Experiments were conducted during the light cycle and included littermate controls. *Prnp-*TARDBP*Q331K mice overexpress human TAR DNA binding protein with the ALS-linked Q331K mutation (B6.Cg-Tg(Prnp-TARDBP*Q331K)103Dwc/J, Jackson Laboratory, strain #017933).

### HSV-1 and HCMV stocks

McKrae strain of HSV-1 was produced and titered in the Shukla Lab. HSV-1 H129-772-eGFP strain was generously provided by Dr. Lynn Enquist (Princeton University, Princeton, NJ). Viral stocks were grown in Vero cells (ATCC, CCL-81) maintained in DMEM (Corning #15-017-CV) with 10% FBS (VWR #89510-188), 1% Sodium Pyruvate (Gibco #11360070), 1% GlutaMAX Supplement (Gibco #35050061), and 1% PenStrep (Gibco #15140122). Prior to experiments, standard plaque assays were performed to titer viral stocks as described below in the <u>Plaque assay</u> section. HSV-1 H129-373-mCherry strain was produced and titered by the University of Pittsburgh Center for Neuroanatomy with Neurotropic Viruses and affiliated Princeton University Herpesvirus Core. Human cytomegalovirus (HCMV) AD169r-BAC-eGFP strain was provided by Dr. Sallie Permar (Weill Cornell Medicine, NY). HCMV stocks were produced and titered as described.^113,114^

### Plasmid generation

Plasmids encoding shRNA targeting human TDP-43 or a nontargeting control sequence were cloned using standard cloning methods. First, pAAV-CAG-tdTomato (Addgene 59462) was digested with KpnI and HindIII to remove the tdTomato sequence and the backbone was gel-purified. mCherry-miR30-shLuc sequence from pAAV-Gfa-mCherry-miR30-shLuciferase (Addgene 178588) was PCR-amplified with Platinum SuperFi DNA polymerase master mix (Thermo) using primers designed to overlap the purified pAAV-CAG backbone (forward primer: 5’-caaagaattggatccggtaccgccaccatggtgagcaag; reverse primer: 5’-cagaggttgattatcgataagcttgatccctcgacggatcc). pAAV-CAG-mCherry-shLuciferase, a plasmid enabling expression of an mCherry reporter and shRNA expression within a mIR-30 scaffold regulated by general CAG promoter, was then generated by NEB HiFi DNA Assembly of pAAV-CAG backbone with mCherry-shLuciferase insert following manufacturer recommendations. pUC shuttle vectors containing XhoI- and EcoRI-flanked shRNA sequence targeting human TDP-43 (5’- gcaggtcaagaaagatcttaa)^115^ or a nontargeting control sequence (5’-cctcccgaattgtcgaagtcg) were synthesized (Azenta/Genewiz) and miR-30-shRNA sequences were digested and ligated into XhoI and EcoRI-digested pAAV-CAG-mCherry-Luciferase to produce pAAV-CAG-mCherry-shTDP43 and pAAV-CAG-mCherry-Scr control.

pcDNA3.2-TDP-43-YFP plasmid (Addgene 84911) containing the CMV promoter was used to overexpress wild-type human TDP-43. The control pCMV-mCherry plasmid was generated by deleting the MHC-IIA coding sequence from Addgene plasmid 35687 using NEB Q5 mutagenesis. Primers were designed according to manufacturer recommendations to retain the stop codon, and the final plasmid was verified by restriction digestion and sequencing.

A plasmid encoding a pUC shuttle vector and a sequence encoding the nuclear-cytoplasmic shuttle protein (pUC-GW-Amp-tdTomato-NES-NLS) was synthesized by Azenta/Genewiz. The tdTomato-NES-NLS sequence was subsequently digested out of the pUC shuttle and ligated into pAAV-GFAP-EGFP (Addgene plasmid 50473) in place of the EGFP sequence to produce a plasmid encoding the tdTomato-NES-NLS shuttle downstream of the GFAP promoter.

Plasmids encoding human FAK (pCMV-mCherry-FAK-HA) and FAK-Y397F (pCMV-mCherry-FAK-Y397F-HA) were generated using NEB HiFi Assembly based on manufacturer recommendations. The plasmid backbone containing the CMV promoter and downstream HA-stop sequences was gel-purified following restriction digest of Addgene plasmid 35039 with HindIII, NheI, and NotI (Thermo). For HiFi assembly, a full-length mCherry insert, a full-length human FAK insert, and two human FAK-Y397F inserts (N- and C-terminal portions) were amplified using Platinum SuperFi DNA polymerase master mix and gel purified. mCherry sequence was amplified from Addgene plasmid 35039 using forward primer 5’-gaaccgtcagatccgctagcgctaccggtcgccac and a reverse primer 5’- gtaagcagctgccatagatctcttgtacagctcgtc containing a BglII restriction site between the mCherry and human FAK sequences. Full-length human FAK and FAK-Y397F were amplified without stop codons from Addgene plasmid 216540. For human FAK, FAK-N-term forward primer 5’-gacgagctgtacaagagatctatggcagctgcttac and FAK-C-term reverse primer 5’- gtatgggtaaaagatgcggccgtgtggtctcgtctg were used. For the FAK-Y397F mutant, two inserts were generated that overlapped and incorporated the point mutation. For the N-terminal portion, FAK-N-term forward primer and FAK-YF-N-term reverse primer 5’-ctataatctcagcaAaatcatctgtttctg were used. For the C-terminal portion, FAK-YF-C-term forward primer 5’- cagaaacagatgattTtgctgagattatag was used with FAK-C-term reverse primer. pCMV-mCherry-FAK- HA was assembled using purified CMV backbone and mCherry-BglII and FAK amplicons. pCMV-mCherry-FAK-397F-HA was assembled using purified CMV backbone and mCherry-BglII, FAK-YF- N-term, and FAK-YF-C-term amplicons. Assembly products were transformed into NEB-5α cells (NEB) and purified plasmids were fully verified by sequencing (Azenta Genewiz).

### Corneal scarification

Corneal scarification was used to infect mice as described previously, with minor modifications.^26^ Mice were anesthetized with sterile Avertin (2,2,2-tribromoethanol, 400 mg/kg body weight, Acros Organics). Meloxicam (2 mg/kg) was injected subcutaneously, and proparacaine was applied topically to relieve pain. Under a light microscope, a 3x3 superficial scratch pattern was made on the left cornea using a 30-gauge needle. Vehicle or 5 x 10^5^ plaque-forming units (PFU) of McKrae strain HSV-1 diluted to 5 µL in vehicle were applied to scratched cornea using a P10 pipette tip. Vehicle control was prepared by sonicating Vero cells five times for 5 s each at 10% power using a probe sonifier (Branson) in 5% autoclaved milk (autoclaved on liquid cycle three times for 15 mins each). Meloxicam (2 mg/kg) was injected subcutaneously every 24 h for 3 days post infection. Mice were weighed daily for 8 days after which they were anesthetized with Avertin (2,2,2- tribromoethanol, 500–600 mg/kg body weight, Acros Organics) and transcardially perfused for 2.5 min with 7.5 mL 0.9% saline. For the plaque assay, the ipsilateral trigeminal ganglion and forebrain were collected into inoculation media (DMEM with 4.5 g/L glucose (Corning #15-017-CV) with 2% fetal bovine serum (FBS, VWR #89510-188), 1% Sodium Pyruvate (Gibco #11360070), 1% GlutaMAX Supplement (Gibco #35050061), and 1% PenStrep (Gibco #15140122)) on ice or flash frozen on dry ice (no media). Frozen regions were stored at −80°C until used in plaque assay. For RNA analysis, ipsilateral hippocampus was flash frozen on dry ice and stored at −80°C. For immunohistochemistry, hemibrains were removed and stored in fixative (4% paraformaldehyde in PBS) overnight at 4°C on a rocking platform. Contralateral fixed hemibrains were subsequently incubated in cryoprotectant (30% sucrose in PBS) for at least 48 h before sectioning. Mouse brain tissue was sectioned (30 μm thickness) using a SM2010 R sliding microtome (Leica) equipped with a BFS-3MP freezing stage and cooling unit (Physitemp, Clifton, NJ). Free-floating sections were collected into cryopreservative (30% ethylene glycol, 30% glycerol, PBS) for storage at −20°C.

### Cell cultures

All cultures were maintained at 37°C in a humidified 5% CO_2_-containing atmosphere.

#### Primary neuronal cultures

Cortical and hippocampal neurons from postnatal day 0–1 TDP-43*Q331K mouse pups and littermate controls were cultured as described previously,^22^ with minor modifications. Briefly, papain-dissociated cells were filtered through a 0.4-µm cell strainer (Corning #431750) to enrich for neurons, centrifuged at 500 g for 5 min to eliminate small debris, and suspended in complete primary neuronal medium consisting of B-27 Plus Neuronal Culture system (ThermoFisher #A3653401) and 1X GlutaMAX (ThermoFisher #35050061) without antibiotics. Cells were seeded at 50,000–150,000 live cells per cm^2^ into plates coated with poly-D-lysine (0.01% w/v; Sigma P6407). Media was fully exchanged one day after plating (DIV 1) with subsequent half-media exchanges every 3–4 days.

#### Human iPSC maintenance

Mutant human iPSCs and isogenic controls were obtained from Jackson Laboratories (#JIPSC1066, heterozygous for Q331K; #JIPSC1108, heterozygous for M337V; #JIPSC1000 and #JIPSC1104, isogenic controls) and from AnswerALS, created with Cedars-Sinai (**Table S1**). All human iPSC cultures and any subsequent differentiations were maintained at 37°C in a humidified 5% CO2 environment. iPSCs were grown in Essential 8 media (ThermoFisher #A1517001) on cell culture dishes coated with vitronectin (5 µg/ml in PBS, ThermoFisher #A31804). iPSCs were passaged using 0.5M EDTA, pH 8.0 (Corning #46-034-CI). Y-27632 dihydrochloride (10 µM, MedChemExpress #HY-10583) was added to media for 24 h during thawing and passaging for proper adherence and to prevent cell death. Cells were cryopreserved in Stem-Cell Banker (AMSBIO #11890) using Mr. Frosty Freezing Containers (ThermoFisher #5100-0001) before long-term storage in liquid nitrogen.

#### Cortical neuron differentiation

Human iPSCs were differentiated into mature cortical neurons using an optimized protocol based on dual SMAD inhibition and WNT inhibition, as described previously.^42^ Briefly, iPSCs were dissociated into a single-cell suspension with Accutase solution (Innovative Cell Technologies #AT104-500) and plated at a density of 300,000 cells/cm^2^ in Essential 8 media (ThermoFisher # A1517001) onto Matrigel (VWR # BD354277) coated culture dishes. Cells were supplemented with 10 μM of Y-27632 dihydrochloride (MedChemExpress #HY-10583). On days 0–2 of differentiation, cells were maintained by fully exchanging the Essential 6 media (ThermoFisher #A1516401) containing SB-431542 (10 µM, R&D #1614/10), LDN-193189 (100 nM, StemGent #04-0074), and XAV-939 (2 µM, StemGent #3748/10) for efficient induction of anterior neuroectodermal patterning.

On days 3–9 of cortical neuron differentiation, cells were fed daily with Essential 6 medium containing SB-431542 (10 µM) and LDN-193189 (100 nM). On days 9–19 of differentiation, cells were maintained by daily media changes in a 1:1 ratio of Neurobasal media (Fisher Scientific #21103049) and Dulbecco’s modified Eagle medium (DMEM)/F-12 media (ThermoFisher #11320033) supplemented with 1X B27 supplement without vitamin A (ThermoFisher #12587010), 1X N2 supplement (ThermoFisher #17502048), and Penicillin-Streptomycin (PenStrep, ThermoFisher #15140122) for efficient generation of a neurogenic population of cortical neural progenitor cells (NPCs). On day 20 of differentiation, NPCs were dissociated into single-cell suspension using Accutase and plated at a density of 150,000–200,000/cm^2^ in Neurobasal media supplemented with 1X GlutaMAX (ThermoFisher), PenStrep, B27 supplement without vitamin A, recombinant human BDNF protein (R&D #248-BDB), recombinant human GDNF protein (PeproTech #450-10), dbcAMP (Sigma #D0627), and L-Ascorbic Acid (Sigma #A4403) on cell culture plates coated with poly-L-ornithine hydrobromide (Sigma #P3655) overnight followed by mouse laminin protein (R&D #3400-010-02) and human fibronectin protein (R&D #1918-FN-02M) overnight. Cells were supplemented with 10 μM of Y-27632 dihydrochloride (MedChemExpress #HY-10583) for 24 h for proper cell adherence and 10 μM Notch pathway inhibitor DAPT (R&D biosystems #2634/10). On day 21, media was completely exchanged to remove Y-27632 dihydrochloride. DAPT was supplemented in media until day 30. For long-term culture, neurons were maintained in Neurobasal media with supplements as described above without DAPT from day 30 onwards and fed every 4–5 days by replacing 50% of the medium volume.

#### Cortical astrocyte differentiation

Human iPSC maintenance and first 20 days of differentiation are described above and were differentiated into mature cortical astrocytes as previously described.^116^ Briefly, on day 20 of differentiation, cortical progenitors cells were plated at a density of 100,000–150,000/cm^2^ in Neurobasal media supplemented with 1X GlutaMAX (ThermoFisher), PenStrep, B27 supplement without vitamin A, recombinant human BDNF protein (R&D #248-BDB), recombinant human GDNF protein (PeproTech #450-10), dbcAMP (Sigma #D0627), and L-Ascorbic Acid (Sigma #A4403) on cell culture plates coated with poly-L-ornithine hydrobromide (Sigma #P3655) overnight followed by mouse laminin protein (R&D #3400-010-02) and human fibronectin protein (R&D #1918-FN- 02M) overnight. Cells were supplemented with 10 μM of Y-27632 dihydrochloride (MedChemExpress #HY-10583) for 24 h for proper cell adherence.

On day 21 of differentiation, cells were transduced with lentiviral vectors encoding NFIA and m2rtTA to induce astrocytic differentiation.^116^ On days 22–28 of differentiation, the media was supplemented with doxycycline (1 mg/ml, Millipore Sigma #D9891) every other day. On day 29, cells were passaged at a density of 75,000/cm^2^ onto poly-ornithine, laminin, and fibronectin-coated cell culture dishes in ScienCell Astrocyte Medium (ScienCell #1801). Cells were fed twice per week in ScienCell Astrocyte Medium until day 34 of differentiation.

From day 34 onwards, cells were fed every 2–3 days with DMEM/F-12 media (ThermoFisher) supplemented with 1X N2 supplement (ThermoFisher), sodium pyruvate (1 mM, ThermoFisher), GlutaMAX (ThermoFisher), N-acetyl-L-cysteine (5 µg/ml, Millipore Sigma #A9165), and HB-EGF (10 ng/ml, R&D #259-HE) and passaged as needed. Cells were cultured until day 60, when astrocytes were used for experiments.

#### Patient-derived fibroblasts

Dermal fibroblasts were generously provided by Giovanni Manfredi (Weill Cornell Medicine, New York, NY) as de-identified, coded samples to protect patient identity. Samples were obtained from the sources specified in **Table S2**. Fibroblasts were cultured in a growth medium consisting of DMEM with 4.5 g/L glucose (Corning #15-017-CV), 10% heat-inactivated FBS (VWR #89510-188), 1 mM Sodium Pyruvate (Gibco #11360070), 4 mM GlutaMAX Supplement (Gibco #35050061), 1% PenStrep (Gibco #15140122), and 1X Plasmocin treatment (InvivoGen #ant-mpp). Fibroblasts were cultured in 100-mm dishes for line expansion and seeded into 12-well plates for experiments. All fibroblast lines were used for experiments at passages ranging between 7 and 15.

To passage and seed for experiments, cells were rinsed once in DPBS without calcium and magnesium (Corning #45000-434), incubated in 0.05% Trypsin-EDTA (Gibco #25300054) at 37°C for 1–5 min, and mechanically dissociated to lift cells. FBS-containing growth medium was then added to each dish at equal volume of the trypsin reagent to inactivate trypsin. The cell suspension was centrifuged at 1000 g for 5 min at room temperature. After aspirating the supernatant, the cell pellet was resuspended in 1 mL growth medium, and the cell suspension was dispersed into growth medium-containing dishes or well plates.

#### Vero cells

Vero cells (ATCC, CCL-81) were maintained in DMEM (Corning #15-017-CV) with 10% FBS (VWR #89510-188), 1% Sodium Pyruvate (Gibco #11360070), 1% GlutaMAX Supplement (Gibco #35050061), 1% PenStrep (Gibco #15140122). Passage and seeding were conducted similarly to fibroblasts.

### Electrophysiology

Whole-cell patch-clamp recordings were performed at room temperature of 20–22°C in human iPSC-derived neurons (5–7 weeks in culture). Cells were identified in DIC mode using ZEISS Axio Examiner upright microscope and ORCA Fusion Digital CMOS High Resolution Camera (Hamamatsu). Whole-cell membrane currents were amplified by the Multiclamp 700B, and data acquisition was controlled by pCLAMP 11.3 software (Molecular Devices). The Digidata 1550B interface was used to convert digital-analog signals between the amplifier and the computer. Data were sampled at 10 kHz and filtered at 2 kHz. Recording pipettes (6–8 MΩ) were fabricated from borosilicate glass capillaries (1B150F-4, World Precision Instruments) using a PC-100 vertical pipette puller (Narishige). Recording pipettes were filled with (in mM): 126 K-gluconate, 5 HEPES, 0.5 MgCl_2_, and 10 BAPTA (pH adjusted to 7.3 with KOH). The standard bath solution contained (in mM): 150 NaCl, 3 KCl, 2 CaCl_2_, 2 MgCl_2_, 10 HEPES, and 5.5 D-glucose (pH adjusted to 7.3 with NaOH).

sEPSCs were recorded under voltage-clamp mode at a holding potential of −60 mV. Current- voltage (I–V) relationships were recorded by applying 10 mV voltage steps (−90 mV to +50 mV) of 1 s duration per step. Input resistance was calculated from steady-state current at −90 mV. Cell membrane capacitance was measured using the membrane test protocol built into pCLAMP. Rheobase was measured by injecting 500-ms duration current steps in 2 pA increments under current-clamp mode. Off-line analysis was conducted using Clampfit version 11.3.

### Viral infection in cultures

Cells were infected with HSV-1 H129-772-eGFP or HSV-1 H129-373-mCherry at multiplicity of infection (MOI) 0.01 for 72 h in normal cell media, unless otherwise indicated. Some cells were exposed to HCMV AD169r-BAC-eGFP at indicated MOI (0.001–5 MOI) for 72 h in normal cell media. Similar MOIs for HSV-1 and HCMV have been used previously in neural cells, including human cells.^22,117–121^ eGFP fluorescence was analyzed after cultures were fixed with 4% paraformaldehyde (PFA) in PBS for 10–15 min and immunolabeled for vimentin (fibroblasts), MAP2 (neurons), and/or stained with DAPI to visualize nuclei (all cell lines).

Fluorescence intensity (integrated density), DAPI area, and percent of total area were extracted using FIJI. Data were background-corrected using non-infected controls and represented as eGFP integrated density normalized to the total DAPI-positive area and to infected controls.

Alternatively, cells were collected for DNA or RNA analysis. RNA extraction and analysis is described in *RT*-*qPCR* section. DNA was extracted using the Quick-DNA/RNA MagBead Kit (Zymo Research #R2130). Cultured cells were rinsed once with PBS and stored at 4°C or 20°C in provided DNA/RNA Shield until DNA extraction. DNA analysis is described in RT-qPCR section.

### Plaque assay

Standard plaque assays to quantify viral load in brain lysates from infected mice and to titer viral stocks were performed as described previously, with minor modifications.^26^ Briefly, Vero cells were seeded to achieve 80% confluency on day of assay inoculation (typically 24 h after plating). On day of inoculation, samples were sonicated in inoculation media (DMEM (Corning #15-017-CV) with 2% FBS (VWR #89510-188), 1% Sodium Pyruvate (Gibco #11360070), 1% GlutaMAX Supplement (Gibco #35050061), 1% PenStrep (Gibco #15140122)) three times for 5 s each at 15% power using a probe sonifier (Branson). Sonicated samples were serially diluted in inoculation media and used to infect Vero cells at 25% of well volume. Plates were incubated normally (37°C, 5% CO_2_) and manually rocked every 15 min. After 2 h, inoculum was removed, and cells were further incubated in inoculation media containing 1% methyl cellulose (Millipore Sigma #M0512) (37°C). After 72 h, methanol was added directly to the medium at a 1:1 ratio and incubated at ambient temperature for 10 min. Methanol containing media was aspirated, and cells were counterstained with crystal violet (Sigma-Aldrich #C6158) (70% water, 20% ethanol, 10% crystal violet stock [1% crystal violet in 20% ethanol]) for 30 min followed by two quick washes with water. Viral titers were determined by visually counting plaques.

### Viral binding assay

The viral binding assay was performed as described previously, with minor adjustments.^76^ Human iPSC-derived neurons were infected on ice at MOI 5 or 10 using chilled HSV-1 H129-772-eGFP inoculum in standard growth media. After a 30 min incubation at 4°C, cells were washed with cold PBS, fixed with 4% PFA in PBS for 15 min at ambient temperature, and washed three times 5 min with PBS. Cells were blocked with PBS with 6% normal donkey serum (Jackson ImmunoResearch) at ambient temperature. Immunocytochemistry for HSV-1 (1:500; Abcam #ab9533) was conducted in PBS with 3% normal donkey serum at 4°C overnight. Samples were protected from light in all subsequent steps. Cells were rinsed three times with PBS for 5 min and incubated with the AlexaFluor555-conjugated secondary antibody (1:500; ThermoFisher #A31572) and 1:3000 14 mM DAPI (Sigma #D9542) diluted in PBS with 3% normal donkey serum for 1 h at ambient temperature. Cells were rinsed three times 5 min with PBS and whole-well images at 20 X magnification were acquired and stitched using a BX-X710 microscope and software (Keyence).

Additional high-resolution 40X magnification images were taken on a LSM 880 Laser Scanning Confocal Microscope (Zeiss). Counts of HSV-1 puncta, total DAPI area, and percent of total area were extracted using FIJI. Data were background corrected using non-infected controls and are represented as number of puncta normalized to controls.

### Viral entry assay

Human iPSC-derived neurons were infected with HSV-1 H129-772-eGFP at MOI of 10 using cold inoculum in standard growth media on ice. After a 30-min incubation at 4°C, cells were further incubated for 15 min at 37°C (standard cell maintenance incubator). Cells were rinsed with 50 mM sodium citrate buffer (pH 3) and collected for Western blot of HSV-1 tegument protein VP16 (1:1000; Abcam #ab110226) (described in *Western blotting* section). Data were background corrected, normalized to non-infected controls, to β-actin levels per sample, and to infected controls.

Patient-derived fibroblasts were infected with McKrae strain HSV-1 at MOI of 50 using cold inoculum in CO_2_-independent medium (ThermoFisher #18045088) supplemented with 10% heat-inactivated FBS (VWR #89510-188), 1 mM Sodium Pyruvate (Gibco #11360070), 4 mM GlutaMAX Supplement (Gibco #35050061), and 1% PenStrep (Gibco #15140122) on ice. After 1-h incubation at 4°C, cells were incubated for an additional 30 min at 37°C in a standard cell culture incubator.

Cells were washed with cold PBS, fixed with 4% PFA in PBS for 15 min at ambient temperature, and washed three times 5 min with PBS. Immunocytochemistry for VP16 (1:200; Abcam #ab110226) and vimentin (Abcam #ab24525) was conducted in PBS containing 0.1% Triton X-100 (PBS-T) with 3% normal donkey serum for 1 h at ambient temperature. Samples were protected from light in all subsequent steps. Cells were rinsed three times with PBS-T for 5 min and incubated with the AlexaFluor488- and AlexaFluor647-conjugated secondary antibodies (1:500; ThermoFisher #21202 and #A31572) and 1:3000 14 mM DAPI (Sigma #D9542) diluted in PBS-T with 3% normal donkey serum for 1 h at ambient temperature. Cells were rinsed two times for 5 min with PBS-T and for 5 min with PBS. Images were acquired using the ImageXpress Micro Confocal High-Content Imaging System (Molecular Devices) with a 10X objective (Nikon) and stitched using MetaXpress software. Counts of VP16 puncta and vimentin areas were extracted using FIJI. Data are shown as number of puncta normalized to vimentin and genotype controls.

### TDP-43 knockdown and overexpression

Given their high transfection efficiency, we used HEK 293T cells to test the effects of acute TDP-43 knockdown and overexpression. Plasmids were generated as described in <u>Plasmid Generation</u>. For knockdown experiments, HEK 293T cells were transfected with a plasmid encoding an shRNA against human TDP-43 (pAAV-CAG-mCherry-shRNA-hTDP43) or a nontargeting control sequence (pAAV-CAG-mCherry-shCtl). For overexpression experiments, HEK 293T cells were transfected with a plasmid encoding wild-type human TDP-43 (pcDNA3.2-TDP-43-YFP) or a fluorescent reporter as a control (pCMV-mCherry).

HEK 293T cells were seeded onto plates coated with poly-D-lysine (0.01% w/v; Sigma P6407) and cultured in growth media consisting of DMEM without glutamine or pyruvate (Corning) supplemented with 10% heat-inactivated FBS (VWR), 1X GlutaMAX (Thermo Fisher), and 100 U/mL Penicillin/Streptomycin (Thermo Fisher). At 50–60% confluency, cells were transfected with 2 µg of plasmid using Lipofectamine 3000 (Thermo Fisher), according to manufacturer protocols. After 18–24 h, the media was changed to fresh media containing HSV-1 H129-722-eGFP at multiplicity of infection 0.1 or 0.01. HEK 293T cells that were exposed to HSV-1 at MOI 0.1 were collected after 48 h, and HEK 293T cells exposed to HSV-1 at MOI 0.01 were collected after 72 h.

### Immunocytochemistry

All immunolabeling steps were performed at ambient temperature unless specified otherwise. Briefly, cells were fixed with 4% PFA in PBS for 10–15 min and permeabilized with PBS (Corning) containing 0.3% Triton X-100 (or 0.1% Triton X-100 for iPSC-derived neurons). Cells were blocked for 1 h in 6% normal donkey or goat serum containing 0.3% Triton X-100 in PBS (or 0.1% Triton X-100 in PBS without serum for iPSC-derived neurons) for 1 h. For some experiments, the blocking step was omitted to improve immunolabeling. Cells were incubated overnight at 4°C with the following primary antibodies in 3% normal donkey or goat serum in 0.3% Triton X-100 in PBS (or 0.1% Triton X-100 for iPSC-derived neurons): chicken anti-MAP2 (1:1000, Abcam #ab5392), chicken anti-MAP2 (1:1000, Novus Bio #NB300-213), rabbit anti-phospho-FAK (Tyr397, 1:250, Thermo Fisher #700255), rabbit anti-NeuN (1:1000, Abcam #ab177487), rabbit anti-Iba1 (1:1000, Wako #01919741), goat anti-Iba1 (1:1000, Abcam #ab5076), rabbit anti-SOX9 (1:500, Millipore #AB5535), chicken anti-Vimentin (1:1000, Abcam #ab24525), mouse anti-TDP-43/TARDBP (1:500, R&D Systems #MAB7778), goat anti-GFP (1:1000, Abcam #ab6658), and rabbit anti-RFP (1:1000, Abcam #ab34771). Samples were protected from light in all subsequent steps. Cells were rinsed three times with PBS with 0.3% Triton X-100 and incubated for 1 h with the following AlexaFluor-conjugated secondary antibodies and 4.7 µM DAPI diluted in 3% normal donkey or goat serum, and 0.3% Triton X-100 in PBS (or 0.1% Triton X-100 in PBS without serum for iPSC-derived neurons) (1:500; ThermoFisher #A21206, A21447, A31570, A31572, A31573, A78948, A78959). Cells were rinsed with PBS containing 0.3% Triton X-100 (or 0.1% Triton X-100 in PBS without serum for iPSC-derived neurons), and then PBS alone before imaging.

### Immunohistochemistry

Double or triple immunolabeling of free-floating mouse sections was performed with minor modifications depending on the antibodies used. All steps were performed at ambient temperature unless specified. Cryopreserved sections were rinsed in PBS, permeabilized for 30 min or longer in PBS containing 0.5% Triton X-100 (PBS-T), blocked with 6% normal donkey or goat serum in PBS-T for 1–2 h, incubated in primary antibodies (below) in 3% normal donkey or goat serum in PBS-T for up to 48 h at 4°C, and rinsed with PBS-T for 8 min four times. Sections were protected from light in all subsequent steps. Tissue was incubated with 1:500 AlexaFluor secondary antibodies (ThermoFisher) (below) and 4.7 µM DAPI in 3% normal donkey or goat serum in PBS-T for 2 h, rinsed with PBS-T for 8 min three times and with PBS for 5 min two times, mounted, and dried on Superfrost glass slides (VWR #75799-266) before sealing #1.5 coverglass (VWR #89239-734) with Vectashield Vibrance Antifade Mounting Medium (#H-1700) or Prolong Diamond Antifade Mounting Media (ThermoFisher #P36970). Slides were allowed to set overnight before acquiring images.

Unless co-labeled with an antibody requiring modified protocols described below, the following primary antibodies were used according to the general protocol: rabbit ant-HSV1 (1:1000, Abcam #ab9533), goat anti-Iba1 (1:400; Abcam #ab5076), and mouse anti-TDP-43 (1:500; R&D Systems #MAB7778; raised against amino acid sequence 1–103 in human TDP-43, which is identical to the sequence in mice). For TDP-43 detection, antigen retrieval was performed for 15 min in 0.1 M citrate buffer at pH 6.0 at 115°C. Sections were subsequently rinsed for 10 min with PBS-T before blocking according to the general protocol above. AlexaFluor-conjugated secondary antibodies were obtained from ThermoFisher (1:500; #A11055, #A78948). A biotinylated donkey anti-rabbit secondary antibody was used to improve HSV-1 primary detection (1:500; Jackson ImmunoResearch #711065152). This incubation was followed by three additional 5-min rinses with PBS-T, followed by a 2 h incubation with an Alexa 594-conjugated Streptavidin (1 µg/ml; Jackson ImmunoResearch #016580084). Brain sections were rinsed and mounted as above.

### Microscopy and image analyses

Human iPSC-derived neurons were evaluated for neural markers on BX-X710 microscope with a 20X objective (Nikon). Briefly, images of individual channels were thresholded and the proportion of DAPI-positive cells positive for each marker was determined. Additional high-resolution images at 20X magnification were taken on a Dragonfly High Speed Confocal Microscope (Andor).

Cells infected with eGFP-tagged HSV-1 were imaged to evaluate fluorescence intensity using BX-X710 microscope with a 10X objective (Nikon). Data were background-corrected using noninfected controls and normalized to infected control cells. Additional high-resolution images at 40X magnification were acquired on a LSM 880 Laser Scanning Confocal Microscope.

HSV-1 expression in brain tissue was evaluated using BX-X710 microscope with a 4X objective (Nikon). Briefly, whole section stitched images were acquired, and focal regions of infection were identified. Infection cores were identified in the following brain regions: olfactory bulb, anterior olfactory nucleus (cerebral cortex), zona incerta (thalamus), superior colliculus (midbrain), locus coeruleus (pons), motor nuclei of the brainstem (upper medulla), and spinal nucleus of the trigeminal (lower medulla). Additional high-resolution images at 20X magnification were taken on a Dragonfly High Speed Confocal Microscope (Andor). Iba-1 intensity in tissue was evaluated on the BX-X710 microscope (Keyence) with a 40X objective (Nikon). Stitched images centered on the thalamic region in each tissue section. Analysis of Iba1 intensity was conducted in CellProfiler using the Identify Primary Objects module. Data were normalized to mock-infected tissue from NTG mice.

TDP-43 alterations in the thalamic region were imaged using Dragonfly High-Speed Confocal Microscope (Andor, 25X magnification). Z-stacks were analyzed using the max projection function in Fiji. The intensities of nuclear TDP-43 immunoreactivity were measured for pixels within a DAPI-thresholded mask. Intensities of extranuclear TDP-43 immunoreactivity were measured for pixels in the entire region of interest (ROI) after omitting nuclear areas as defined by the DAPI mask. Intensities were corrected for background fluorescence per image by subtracting mean intensity of ROIs drawn in areas that did not have TDP-43 or DAPI labeling above background levels.

To assess TDP-43 localization in WT and Q331K human iPSC-derived neurons (days 35– 38), cells were imaged using a Dragonfly 200 spinning disk confocal microscope (Andor Technology) equipped with a 63x objective (Leica Microsystems). Images were analyzed using FIJI. Briefly, each image was subdivided into four non-overlapping, equally sized fields, and aggregates were quantified within MAP2-positive neurons using the Analyze Particles function with a size threshold of 0.7–1.2. Each data point represents one well and corresponds to the average of four fields per well. These analyses were derived from two independent differentiations.

To quantify relative phospho-FAK levels in WT and Q331K human iPSC-derived neurons (D35–38), cells were imaged on a Dragonfly 200 spinning-disk confocal microscope (Andor Technology) equipped with a 40X objective (Leica Microsystems). Images were analyzed in FIJI (ImageJ). Briefly, MAP2 images were thresholded to generate binary masks, and individual MAP2- positive neurons were selected from the masks using the *Create Selection* function. Each selection was then applied to the corresponding raw MAP2 and phospho-FAK images, and relative integrated densities were quantified for each image. Data are presented as one value per well, calculated as the average intensity across all neurons analyzed per well.

To quantify the percentage of human fibroblasts and iPSC-derived neurons infected with HSV-1 after FAK overexpression, cells were imaged using a Dragonfly 200 spinning-disk confocal microscope (Andor Technology) equipped with a 20x objective (Leica Microsystems). Images were analyzed using FIJI (ImageJ). Briefly, vimentin/MAP2 and RFP channels were thresholded to generate binary masks. The *Image Calculator* function was used to identify RFP-positive, vimentin- positive fibroblasts or MAP2-positive neurons, and the resulting mask was converted to a selection using the *Create Selection* function. The resulting selection was applied to the corresponding raw HSV-1 and RFP images to quantify HSV-1-positive, RFP-positive fibroblasts or neurons. For each well, the percentage of infected cells was calculated as the number of HSV-1-positive, RFP- positive cells divided by the total number of RFP-positive cells and multiplied by 100. Data are presented as a single value per well.

### Nucleocytoplasmic shuttle analyses

HEK 293T cells were seeded at low density onto glass coverslips (Warner Instruments) coated with poly-D-lysine (0.01% w/v; Sigma P6407) and cultured in HEK growth media, as described above. At 25–30% confluency, HEK 293T cells were exposed to HSV-1 H129-772-eGFP at MOI 0.1. The following day, cells were transfected with the plasmid pAAV-GFAP-tdTomato-NES-NLS (1 µg) using Lipofectamine 3000 (Thermo Fisher), according to manufacturer’s protocols. After 48 h of viral exposure, cells were fixed and immunolabeled with rabbit anti-RFP (Abcam #ab34771) and DAPI, as described in <u>Immunocytochemistry</u> section. Higher MOI for shorter infection time (48 h) was used to ensure widespread infection while minimizing excessive cell growth and division, which enabled imaging of distinct cells.

To measure extranuclear-to-nuclear ratios of tdTomato-NES-NLS signal, cells were imaged on a Dragonfly 200 spinning disk confocal microscope (Andor) with a 40x objective (Leica) and analyzed using FIJI. For each ROI, each channel was brightness-adjusted and contrast-adjusted, and thresholded to capture signal above background. Brightness/contrast and channel threshold values were consistent among images. Each data point represents one coverslip. Extranuclear tdTomato-NES-NLS levels were calculated by subtracting the DAPI mask from the total tdTomato- NES-NLS mask, and nuclear tdTomato-NES-NLS was calculated by subtracting this extranuclear tdTomato-NES-NLS mask from the total tdTomato-NES-NLS mask. To exclude debris, nuclear and extranuclear tdTomato-NES-NLS masks were size-excluded to remove objects with fewer than 100-pixel units. Total tdTomato-NES-NLS levels were calculated by adding raw integrated densities of nuclear and extranuclear tdTomato-NES-NLS. Ratios of extranuclear-to-nuclear tdTomato-NES- NLS were derived by dividing extranuclear/total by nuclear/total.

### Pharmacological modulation of FAK

To test how changes in FAK activity affect HSV-1 infection levels, human iPSC-derived neurons and isogenic controls were pretreated for 2 h with a FAK inhibitor (1 µM, PF573228, Tocris Bioscience #3239), FAK agonist (1 µM, ZINC40099027, MedChemExpress #HY-134570), or DMSO alone (0.1%) as vehicle control diluted in neuronal media. Following pre-treatment, neurons were also exposed to HSV-1 H129-772-eGFP for either 72 h at MOI 0.01 (for the inhibitor experiments) or 24 h at MOI 1.0 (for the agonist experiments). Every 24 h, 1 µM FAK inhibitor was supplemented into the existing cell media to help maintain inhibition throughout the 72-h infection period. Given the fast onset of action of the FAK agonist (within one hour),^122^ we reasoned that FAK activation would promote HSV-1 infection in earlier stages of infection. We therefore focused on the 24-h timepoint and used a higher MOI of 1.0 to model fast pharmacokinetics of the agonist and achieve sufficient levels of infection within the shorter timeframe. Neurons were collected for HSV-1 DNA analyses, as described below in <u>qPCR</u> section, or were fixed and immunolabeled, as described in the <u>Immunocytochemistry</u> section.

### Genetic modulation of FAK

We used patient-derived fibroblasts to test the effects of acute FAK overexpression on HSV-1 infectivity. Plasmids encoding human FAK or FAK-Y397F with an mCherry reporter downstream of the CMV promoter (pCMV-mCherry-FAK) were assembled as described above (see <u>Plasmid</u> <u>generation</u>). Fibroblasts were cultured on 96-well glass-bottom plates and transfected with pCMV-mCherry-FAK, pCMV-mCherry-FAK-Y397F, or mCherry as a control (pCMV-mCherry) with 0.2 µg of plasmid using Lipofectamine 3000 (Thermo Fisher), according to manufacturer’s guidelines. Cells were subsequently exposed to HSV-1 (H129-772-eGFP) at MOI 0.1 for 72 h, fixed in PFA, and co-immunolabeled for vimentin, eGFP, and RFP, as described in the <u>Immunocytochemistry</u> section. Of note, the transfection procedure altered baseline HSV-1 infection rates, thus infection levels were normalized to mCherry controls per genotype.

We also used human iPSC-derived neurons (D35) to test effects of FAK overexpression on HSV-1 infectivity. These experiments involved similar methods as detailed above, except the neurons were exposed to HSV-1 at MOI 0.01 for 72 h, and were co-immunolabeled for MAP2, eGFP, and RFP, as described in the <u>Immunocytochemistry</u> section.

### Western blotting

For cell cultures, wells were rinsed twice with ice-cold PBS before aspirating buffer and lysing directly with ice-cold 10 mM Tris pH 7.5, 150 mM NaCl, 5 mM EDTA, 0.5% deoxycholate, 0.5% Triton X-100, containing 1X complete Protease Inhibitor Cocktail (Millipore Sigma #11836153001) and 1% each of Phosphatase Inhibitor Cocktails 2 and 3 (Millipore Sigma #P5726 and #P0044). Cells were scraped, collected in 1.5 ml Eppendorf tubes, sonicated on ice for 5 s at 10% power with a probe sonifier (Branson), centrifuged for 5 min at 10,000 g at 4°C, and assayed for protein content using a detergent-compatible Bradford assay (ThermoFisher #23246).

For brain tissue, indicated regions were sub-dissected and flash-frozen on dry ice in 1.5-ml Eppendorf tubes. Frozen samples were thawed on ice for 1–2 min before adding 100–600 µl ice- cold lysis buffer (10 mM Tris pH 7.5, 150 mM NaCl, 5 mM EDTA, 0.5% deoxycholate, 0.5% Triton X-100, containing 1X complete Protease Inhibitor Cocktail (Millipore Sigma #11836153001) and 1% each of Phosphatase Inhibitor Cocktails 2 and 3 (Millipore Sigma #P5726 and #P0044). Samples were immediately homogenized using a hand-held motorized homogenizer (Bel Art), sonicated on ice for 5 s at 10% power with a probe sonifier (Branson), centrifuged for 5 min at 10,000 g at 4°C, and assayed for protein content as above.

For tissue and cell culture extracts, 15–25 µg RIPA-soluble lysates were resolved on Bis- Tris SDS-PAGE gels (ThermoFisher) and transferred onto nitrocellulose membranes using an iBlot2 Western blotting system or a Mini Blot Module (ThermoFisher). Membranes were blocked with 5% non-fat milk in TBS before probing overnight at 4°C with primary antibodies diluted in TBS containing 0.2% Tween-20 (TBS-Tw) and 3% bovine serum albumin (BSA). Primary antibodies were raised against TDP-43 (1:2000 pan-specific rabbit anti-TDP-43, ProteinTech #10782-2-AP), HSV-1 protein VP16 (1:1000; mouse; Abcam #ab110226), β-actin (1:2000; rabbit; Sigma #A2066), and γ-tubulin (1:1250; mouse; Sigma #T5326).

After overnight incubation in primary antibodies, all blots were rinsed with TBS-Tw and probed for 1 h with IR Dye 680RD donkey anti-mouse (1:15,000; LI-COR #926-68072) and IR Dye 800CW donkey anti-rabbit (1:15,000; LI-COR #926-32213) in TBS-Tw with 3% BSA. Blots were rinsed twice with TBS-Tw, once with TBS, and dried for at least 20 min before scanning on the Odyssey CLx imaging system (LI-COR). Expression levels were quantified using LI-COR Image Studio software. Background was subtracted by manual selection of background regions or by LI- COR Image Studio software median correction above and below specific bands.

### RT-qPCR

RNA was extracted using the RNeasy Mini Kit with on-column DNase treatment following manufacturer instructions (Qiagen #74106, #79256). Cultured cells were rinsed once with ice-cold PBS, scraped in freshly prepared extraction buffer and frozen at −80°C until extraction. Saline- perfused, microdissected mouse brain tissue was frozen on dry ice and stored at −80°C until RNA extraction. Tissue was homogenized in RLT buffer containing freshly added β-mercaptoethanol using a bead mill for 20 s at a speed setting of 5 in a pre-chilled adaptor tube rack. All primer sequences are detailed in **Table S3**. Microfluidic RT-qPCR was performed similarly to previously described protocols.^123^ Briefly, cDNA was synthesized with Protoscript First Strand Synthesis Kit (New England Biolabs #E6300L) and pre-amplified for 14 cycles against a pool of primers using PreAmp Grandmaster mix (TATAA Biocenter, Sweden #TA05) before exonuclease I treatment (New England Biolabs #M0293L). Pre-amplified cDNA was diluted at least 5-fold with nuclease- free water and mixed with SsoFast EvaGreen with Low ROX (BioRad #1725211) and chip-specific DNA Sample Reagents before loading into primed Flex Six or 96.96 Dynamic Array chips (Standard BioTools #100-6308 or BMK-M-96.96). Individual primers were mixed with DNA assay reagent (Standard BioTools) and loaded into chip inlets. Chips were primed and loaded using an IFC Controller HX (Standard BioTools) before measuring and analyzing amplification and melting curves on a BioMark HD System (Standard BioTools). Cycle of quantification (Cq) values were thresholded equally for all inlets across each chip run and normalized to the average of reference genes (ACTB and/or TBP1) before determining ddCq and fold-change relative to experimental control groups.

### qPCR

For targeted analysis of HSV-1 DNA (gB), qPCR was performed on isolated DNA using PrimeTime Gene Expression Master Mix (IDT) with target-specific primers and probes, and a CFX96 Touch Real-Time PCR Detection System (BioRad). Primer sequences and probes are detailed in **Table S4**. Cycle of quantification (Cq) values were normalized to the reference gene (*18S*) before determining ddCq and fold-change relative to experimental control groups.

### RNA sequencing

RNA was extracted and cDNA synthesized as described for qPCR. cDNA was submitted to the Weill Cornell Genomics Resources Core Facility for quality control analysis and bulk RNA sequencing. The libraries were sequenced with paired-end 50 bps on a NovaSeqXplus sequencer. The raw sequencing reads in BCL format were processed through bcl2fastq (v2.20) (Illumina) for FASTQ conversion and demultiplexing. The adaptors were trimmed with cutadapt (v1.18) (https://cutadapt.readthedocs.io/en/v1.18/), RNA reads were aligned and mapped to the GRCh38 human or GRCm39 mouse reference genome by STAR (v2.5.2) (https://github.com/alexdobin/STAR),^124^ and transcriptome reconstruction was performed by Cufflinks (v2.1.1) (http://cole-trapnell-lab.github.io/cufflinks/).

The abundance of transcripts was measured with Cufflinks in Fragments Per Kilobase of exon model per Million mapped reads (FPKM).^125,126^ Raw read counts per gene were extracted using HTSeq-count (v0.11.2).^127^ Gene expression profiles were constructed for differential expression, cluster, and principle component analyses with the DESeq2 package (https://bioconductor.org/packages/release/bioc/html/DESeq2.html).^128^ For differential expression analysis, pairwise comparisons between two or more groups were made using parametric tests where read-counts follow a negative binomial distribution with a gene-specific dispersion parameter. Corrected p-values were calculated based on the Benjamini-Hochberg method to adjust for multiple comparisons.

#### Gene network analyses

Qiagen Ingenuity Pathway Analysis (IPA) was used to identify gene networks. Differentially expressed genes (DEGs) were imported into IPA with log2 fold change values and pvaladj to identify involved pathways. Log2 fold change cutoffs were specified at < −0.1 and > 0.1. Pvaladj cutoff was specified at < 0.05. Fisher’s exact test was used to calculate the p-value of the overlap of our dataset with those genes of a reference set known to be regulated by particular upstream factors, with a statistical significance level p < 0.05. Z-scores were calculated, with positive and negative scores indicating activation and inhibition of upstream regulators, respectively.

#### Interferome analyses

Interferome v2.01 (https://interferome.org/interferome/home.jspx) was used to identify DEGs regulated by interferons. DEGs with a pvaladj < 0.05 were input to the database for each comparison. Species parameters were selected (Mus musculus or Homo sapiens) and fold change cutoffs were specified as <-2.0 and >2.0.

#### Alternative splicing analyses

Computational analysis of splicing was performed using the Weill Cornell Medicine High-Performance Computing Core (RRID: SCR_028969). Short-read bulk RNA sequencing data were analyzed using Modeling Alternative Junction Inclusion Quantification (MAJIQ, v2.5.8).^81^ Sequencing reads generated in paired-end mode were aligned to the GRCh38 primary human genome with STAR, employing a reference index derived from GENCODE v44 annotations. STAR-generated BAM files (6 control and 6 Q331K replicates) were provided as input to the MAJIQ *build* module. Using the GENCODE v44 GFF3 annotation, splicing graphs and individual splice junction files were generated for each sample on the GRCh38 reference genome. Differential splicing analysis was then performed using the MAJIQ Heterogen module, which does not assume replicate homogeneity and is therefore suitable for analyzing heterogeneous sample sets.

Heterogen outputs were exported as tab-separated value (TSV) files using the MAJIQ *tsv* module, with local splicing events annotated. These outputs were further processed using custom Python scripts to extract junction-level Ψ values and associated statistical metrics. Junctions were considered differentially spliced if they passed all three MAJIQ statistical tests (*t* test, TNOM, and Wilcoxon; *p* < 0.05) and exhibited an absolute change in Ψ greater than 20% between genotypes.

MAJIQ *Modulize* was used to classify significant local splicing variations into splicing event types. To facilitate biological interpretation and avoid ambiguity arising from differences in exon numbering across transcript isoforms, as well as from the junction-based nature of MAJIQ outputs, we reconstructed junction-level splicing changes into exon-level usage profiles. Exon genomic coordinates were annotated using custom Python scripts based on the MAJIQ *Modulize* output.

### Statistics

Statistical specifications are reported in the figures and corresponding figure legends. Data are presented as mean ± S.E.M, unless indicated otherwise. All statistical tests were performed using GraphPad Prism 9 or 10, except Fisher’s exact test and other statistical tests included in the Qiagen Ingenuity Pathway Analysis. The criterion for data point exclusion was established during the design of the study and was set to values above or below two standard deviations from the group mean. Two-sided Student’s *t* test was used to determine statistical significance between two groups, unless indicated otherwise. Welch’s correction was used to account for unequal variances. Differences among multiple groups were assessed by one-way or two-way ANOVA or mixed- effects model followed by Tukey’s, Dunnett’s, or Bonferroni’s multiple comparisons post-hoc tests, as specified in the figure legends. Null hypotheses were rejected at p < 0.05.

## Supplemental Information

**Fig. S1.**
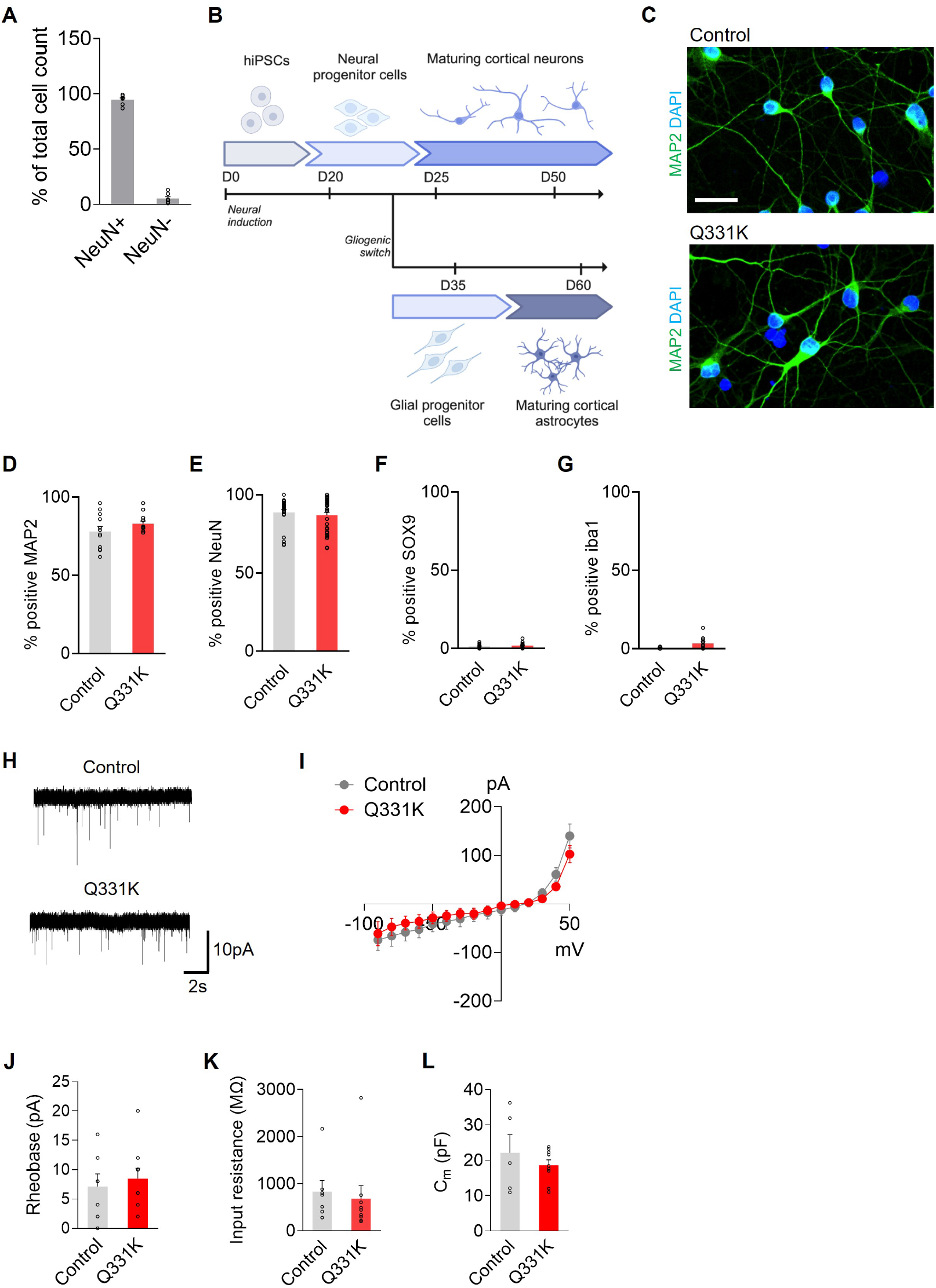
Differentiation and validation of human iPSC-derived neurons. **(A)** Quantification of NeuN-positive cells in mouse-derived primary WT neuronal cultures. n = 7 independent wells. **(B)** Timeline of human iPSC differentiation. **(C)** Immunolabeling of human iPSC-derived neurons for MAP2 (green). Nuclei were labeled with DAPI (blue). Scale bar: 25 µm. (**D**–**G**) In iPSC-derived neurons, percent of DAPI-positive cells that were also positive for neuronal markers MAP2 (D) and NeuN (E), astrocyte marker SOX9 (F), and microglial/macrophage marker Iba1 (G). n = 8–33 ROIs from 5–6 wells in two separate differentiations. **(H)** Representative sEPSC traces recorded at a holding potential of −60 mV. **(I)** Current-voltage relationships in iPSC-derived neurons. n = 7–9 cells per group. **(J)** Averaged rheobase of iPSC-derived neurons. n = 7–9 cells per group. **(K)** Averaged input resistance of iPSC-derived neurons, calculated from steady-state current at −90 mV. n = 7–9 cells per group. **(L)** Averaged membrane capacitance of iPSC-derived neurons. n = 5–9 cells per group.

**Fig. S2.**
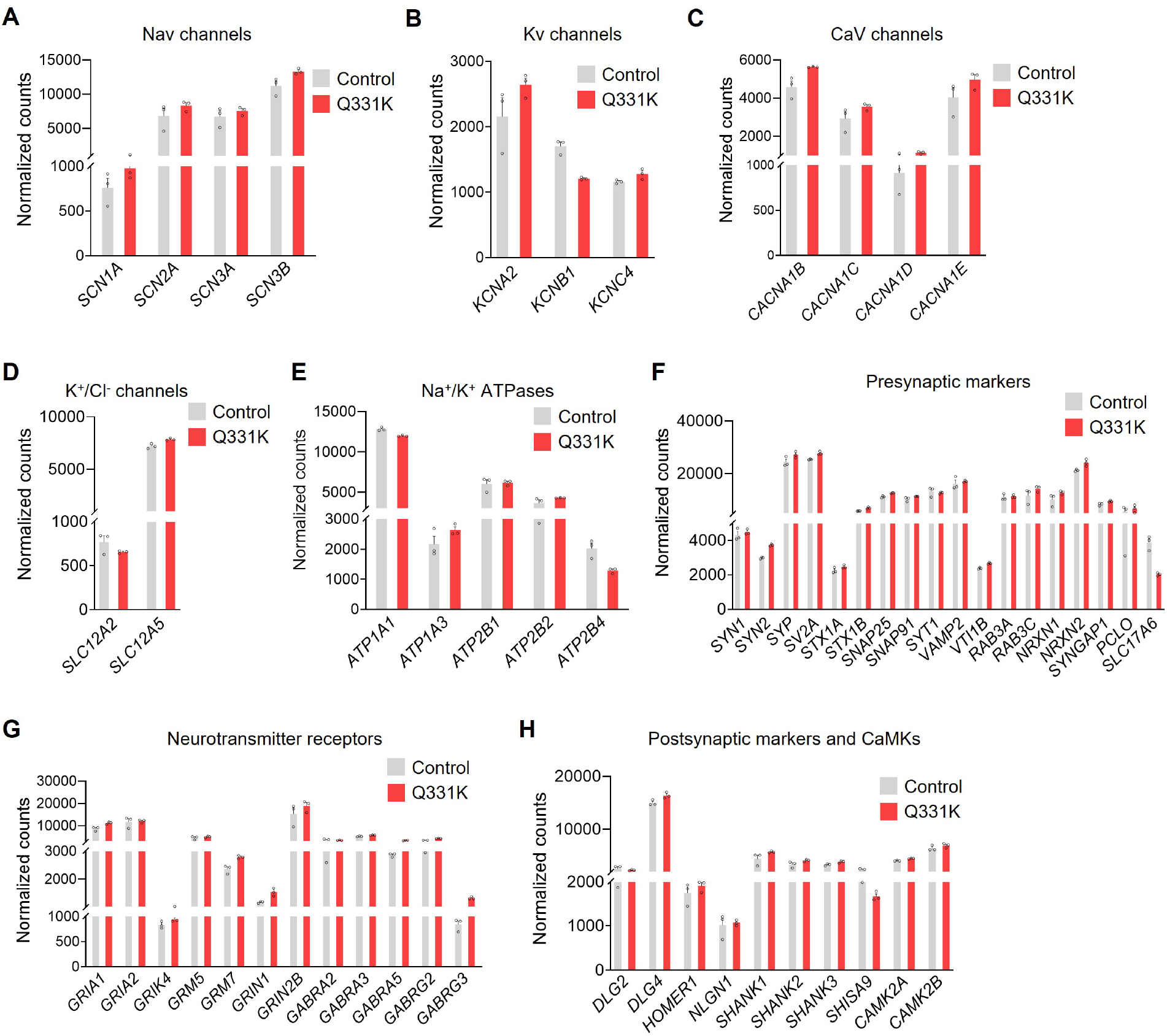
Transcriptional characterization of human iPSC-derived neurons. (**A**–**H**) Normalized counts from RNAseq analyses in human isogenic control and Q331K iPSC-derived neurons. Neuronal marker genes are grouped by functional gene families. n = 3 wells per genotype and gene.

**Fig. S3.**
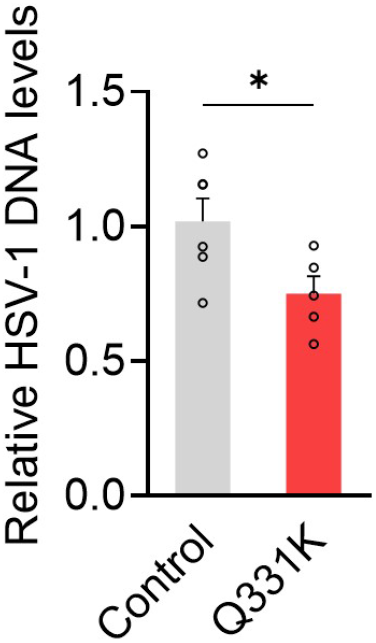
HSV-1 DNA levels in human iPSC-derived astrocytes. Human iPSC-derived astrocytes were acutely infected with HSV-1-eGFP (0.01 MOI) for 72 h and DNA levels of HSV gB were measured by qPCR. Normalized to 18S and control iPSC astrocytes. n = 5–6 wells per genotype. Student’s *t* test: *p = 0.0215.

**Fig. S4.**
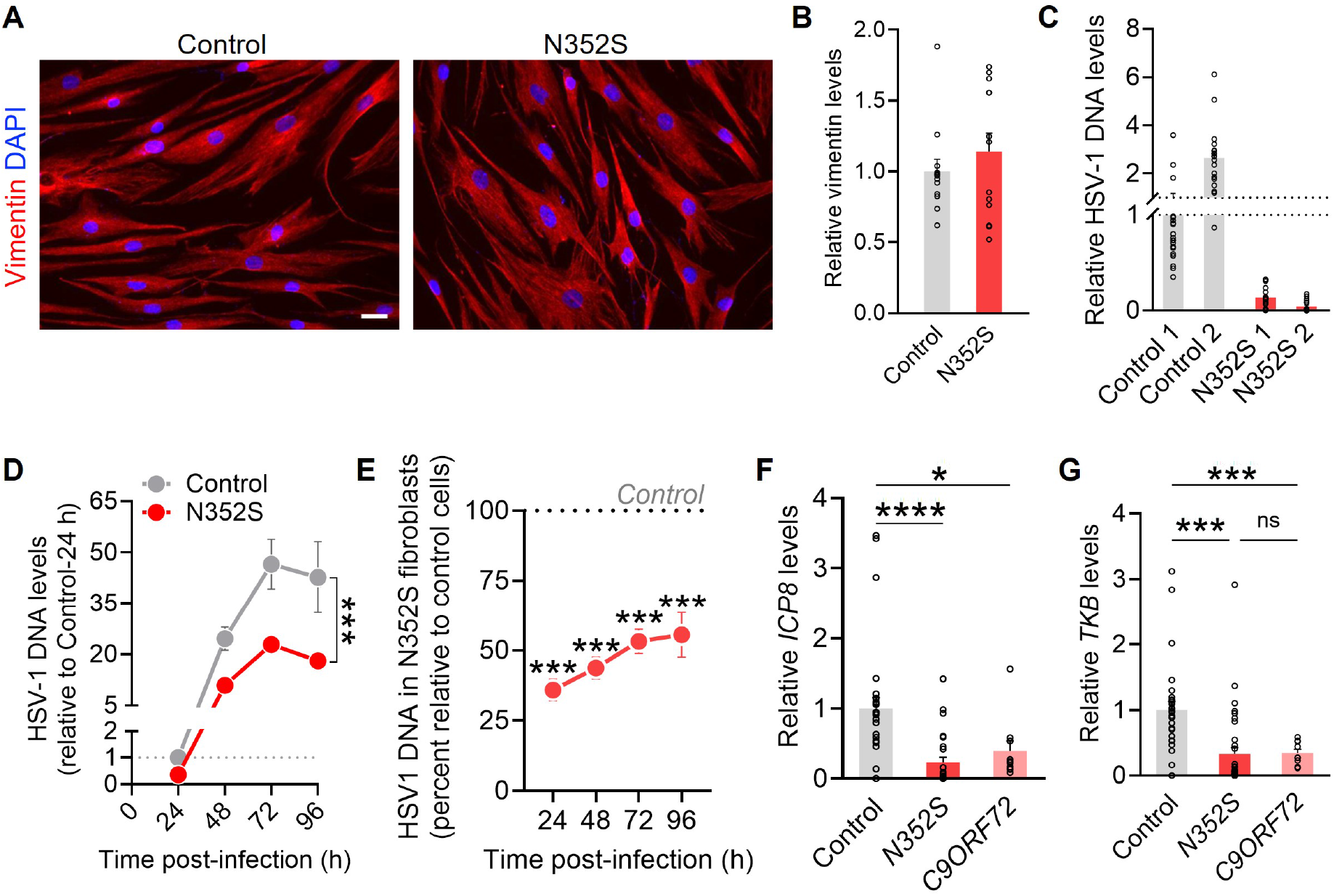
Additional analyses of HSV-1 infection in patient-derived fibroblasts. (**A**, **B**) Immunolabeling (A) and quantification (B) of vimentin (red) in patient-derived fibroblasts. Nuclei were labeled with DAPI (blue). Normalized to control fibroblasts. n = 2–3 wells, 3–4 ROIs per well. Scale bar: 25 µm. (**C**) Data in Fig. 1G separated by patient line. (**D**, **E**) Patient-derived fibroblasts were exposed to HSV-1 for indicated durations and DNA levels of HSV gB were measured by qPCR. Normalized to 18S and either the control genotype at 24 h (D) or the control genotype per time-point (E). Two-way ANOVA: *F*(4, 70) = 2.79, p = 0.033 for interaction effect; *F*(1, 70) = 25.89, p < 0.0001 for genotype effect (D); *F*(1, 56) = 166.9, p < 0.0001 for genotype effect (E). Bonferroni’s test: ***p < 0.0001 vs. Control cells of matching timepoint (E). n = 8 wells per genotype and timepoint from two independent experiments. (**F**, **G**) Patient-derived fibroblasts were exposed to HSV-1-eGFP (0.01 MOI) for 72 h and HSV-1 early transcripts (*ICP8, TKB*) were measured by RT-qPCR. Normalized to *ACTB* and *TBP1* per sample and to controls. n = 10–35 wells from 3–4 patient lines per group. (F) One-way ANOVA: Dunnett’s test: *p = 0.0200, ****p < 0.0001. (G) One-way ANOVA: Dunnett’s test: ***p < 0.0005.

**Fig. S5.**
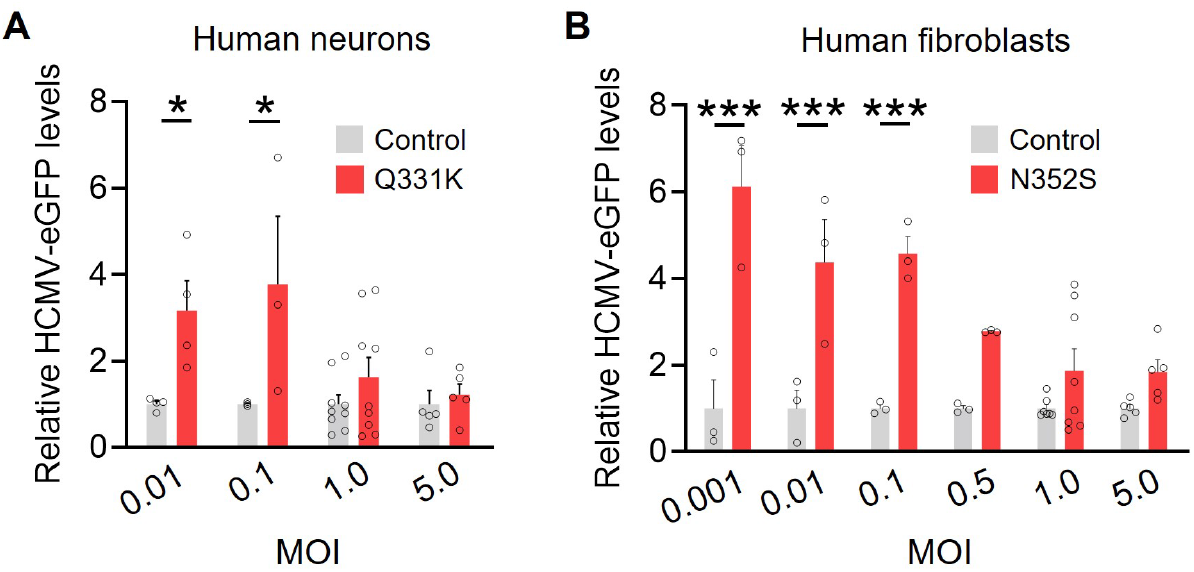
Effects of TDP-43 mutation on HCMV infection. (**A**, **B**) Human iPSC-derived neurons (A) and patient-derived fibroblasts (B) were exposed to HCMV at indicated MOI for 72 h. eGFP fluorescence was measured by immunolabeling, quantifying integrated intensities, and normalizing to DAPI area per well and to control genotype per MOI. Two-way ANOVA: *F*(1, 34) = 15.19, p = 0.0004 for genotype effect (A); two-way ANOVA: *F*(1, 37) = 83.60, p < 0.0001 for genotype effect (B). Bonferroni’s test: *p < 0.05, ***p < 0.001. n = 3–9 wells from 1–2 independent differentiations (A); 3–8 wells from 1–2 independent lines (B).

**Fig. S6.**
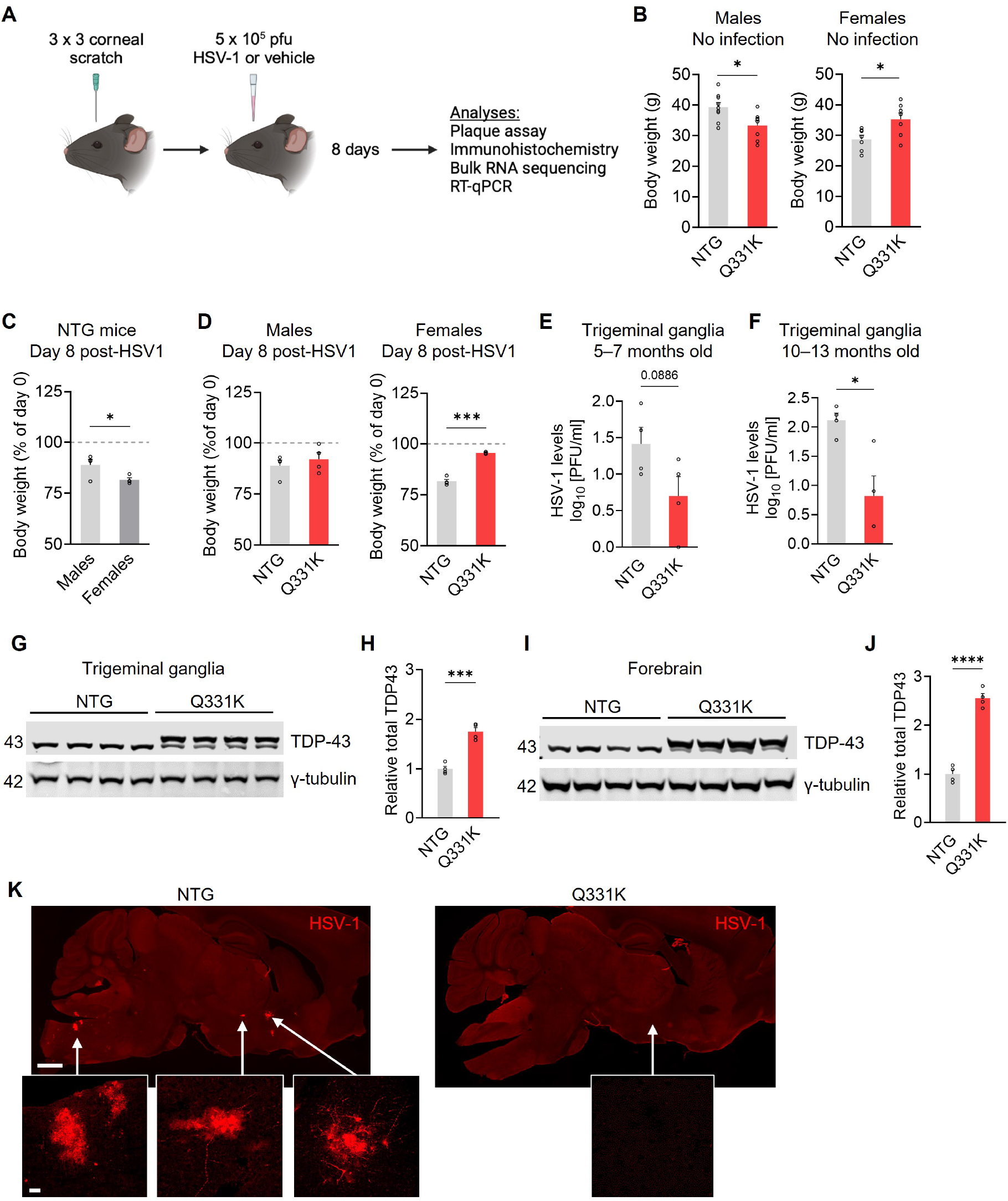
Further analyses of viral infection levels and mutant TDP-43 expression. (**A**) Summary of the viral infection mouse model and experimental design. Cornea of anesthetized mice was scratched using a 30-gauge needle, immediately followed by unilateral inoculation with 5×10^5^ pfu of McKrae strain HSV-1 or vehicle. (**B**) Baseline body weight of male and female NTG and Q331K mice without exposure to virus (mock treatment). 10–13 months old. n = 8–9 males and 7 females per genotype. Student’s *t* test: *p < 0.05. (**C**, **D**) Percent body weight loss in 10–13-month-old male and female NTG (C) and Q331K (D) mice at day 8 after inoculation with McKrae strain HSV-1 through the corneal scarification route. n = 4 NTG males, 4 NTG females, 4 Q331K males, and 3 Q331K females. Student’s *t* test: *p < 0.05, ***p < 0.0001. (**E**, **F**) Plaque-forming units of HSV-1 per ml detected in the trigeminal ganglion at 5–7 months of age (E) and 10–13 months of age (F). n = 4 mice per genotype. Student *t* test: *p = 0.0123. (**G**–**J**) Representative Western blots (G, I) and quantification (H, J) of total (mouse and human) TDP- 43 expression levels in the trigeminal ganglion (G, H) and forebrain (I, J). Normalized to γ-tubulin per sample and to the NTG group. 10–13 months of age. n = 4 mice per genotype. Student’s *t* test: ***p = 0.0003, ****p < 0.0001. (**K**) Additional representative images of HSV-1 immunolabeling. Scale bars: 600 µm, 25 µm (inset).

**Fig. S7.**
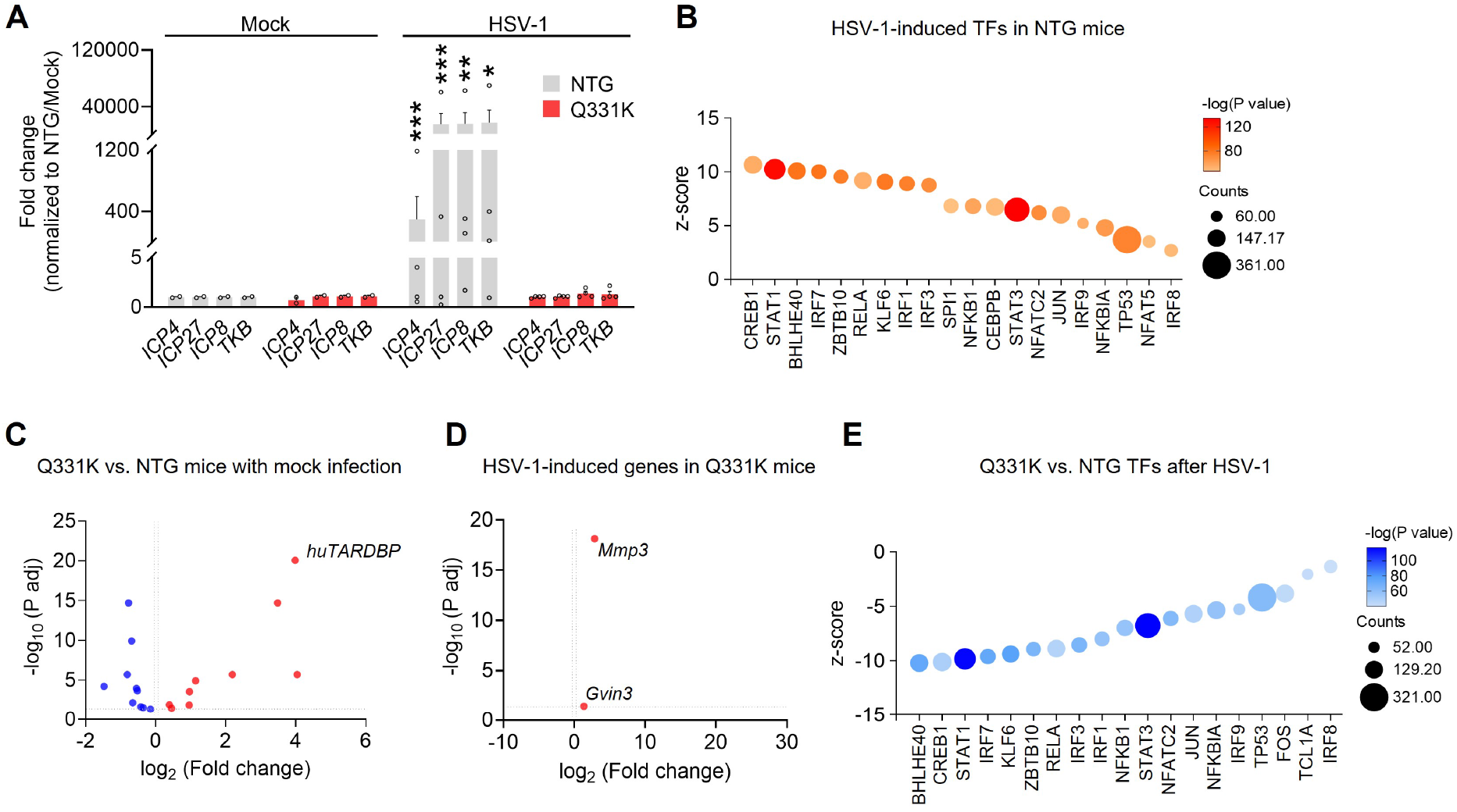
Viral transcripts and transcriptional profiling in mouse hippocampus. **(A)** RT-qPCR analyses of immediate-early (*ICP4, ICP27*) and early (*ICP8, TKB*) HSV-1 transcripts in hippocampal samples from NTG and Q331K mice inoculated with HSV-1 via corneal scarification or receiving mock treatment. Normalized to *Actb* and *Tbp1* per sample and to mock-infected NTG mice. n = 2–4 mice per genotype and condition, 10–13 months old, 8 DPI. One-sample *t* test (vs. theoretical mean 0): ***p < 0.001, **p < 0.01, *p < 0.05. **(B)** Qiagen Ingenuity Pathway Upstream Analysis, transcription factor prediction of differentially expressed genes in main Fig. 2B. Out of the top 25 significant TFs, TFs predicted to be activated are plotted. **(C)** Volcano plot of differentially expressed transcripts in mock-infected Q331K hippocampi versus mock-infected NTG controls. n = 4 mice per genotype, 10–13 months old, 8 DPI. **(D)** Volcano plot of differentially expressed transcripts in Q331K hippocampi from mice inoculated with HSV-1 versus mock-infected Q331K controls. n = 4 mice per genotype, 10–13 months old, 8 DPI. **(E)** Qiagen Ingenuity Pathway Upstream Analysis, transcription factor prediction of differentially expressed genes in main Fig. 2C. Out of the top 25 significant TFs, TFs predicted to be activated are plotted.

**Fig. S8.**
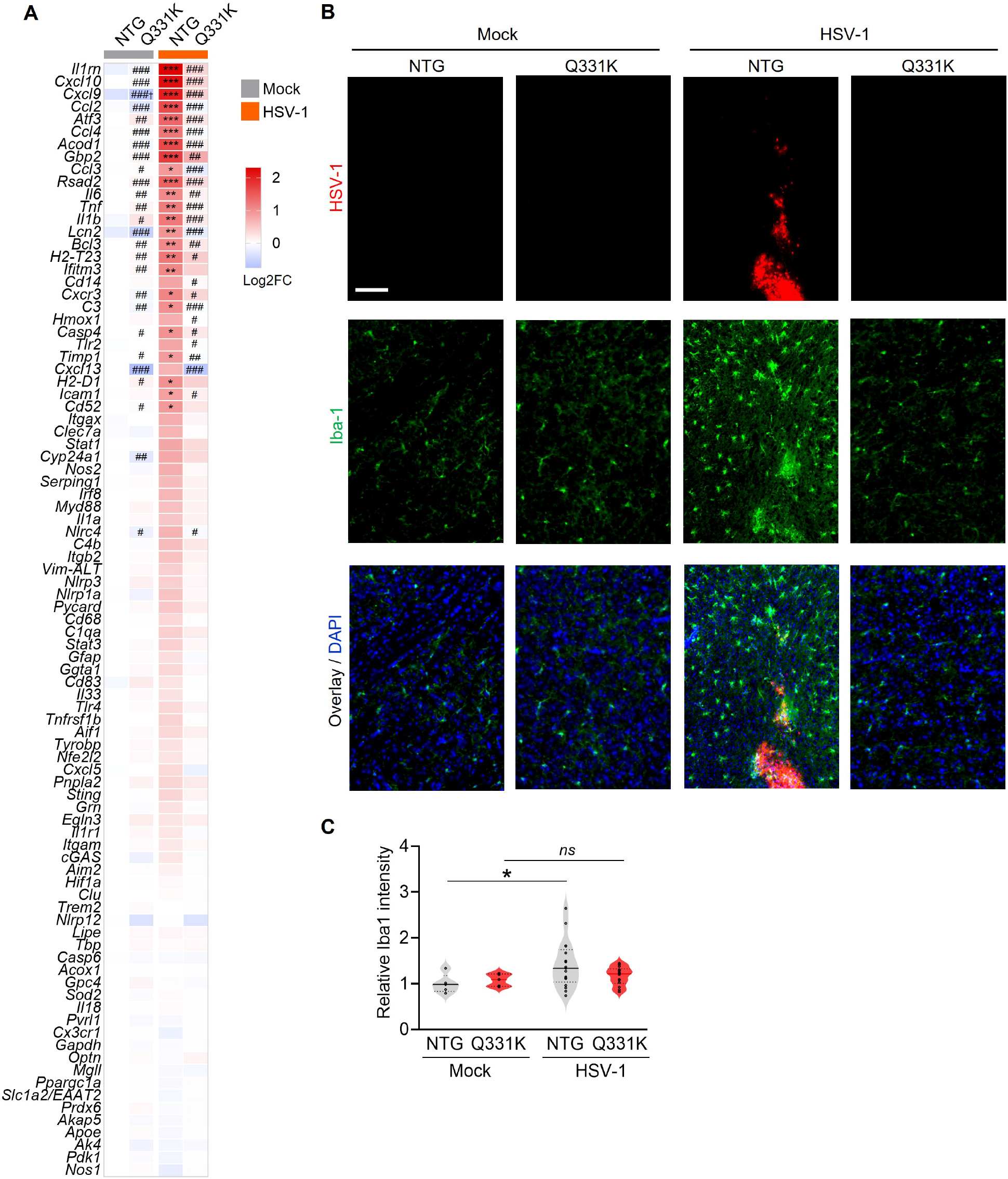
TDP-43 mutation prevents neuroinflammation *in vivo*. **(A)** RNA levels as measured by RT-qPCR. Normalized to *Actb* and *Tbp1* per sample and to RNA levels in mock-infected NTG mice. n = 2–4 mice per condition, 10–13 months old, 8 DPI. Two-way ANOVA, Bonferroni test: *p < 0.05, **p < 0.005, ***p < 0.0005 vs. mock-infected NTGs. Hash tags indicate comparisons to NTG + HSV-1: ^#^p < 0.05, ^##^p < 0.005, ^###^p < 0.0005. Cross indicates comparison to Q331K + HSV-1: p < 0.005. **(B)** Immunolabeling for microglial/macrophage marker (Iba1, green) in focal HSV-1 infection cores (red) in the thalamus. DAPI was used to visualize nuclei (blue). Scale bar: 50 µm. **(C)** Quantification of Iba-1 intensity. n = 3–8, 1–3 sections per mouse; 5–7 months old; 8 DPI. Normalized to NTG / mock. Lognormal Brown-Forsythe ANOVA: p = 0.033; Dunnett’s test: *p = 0.056.

**Fig. S9.**
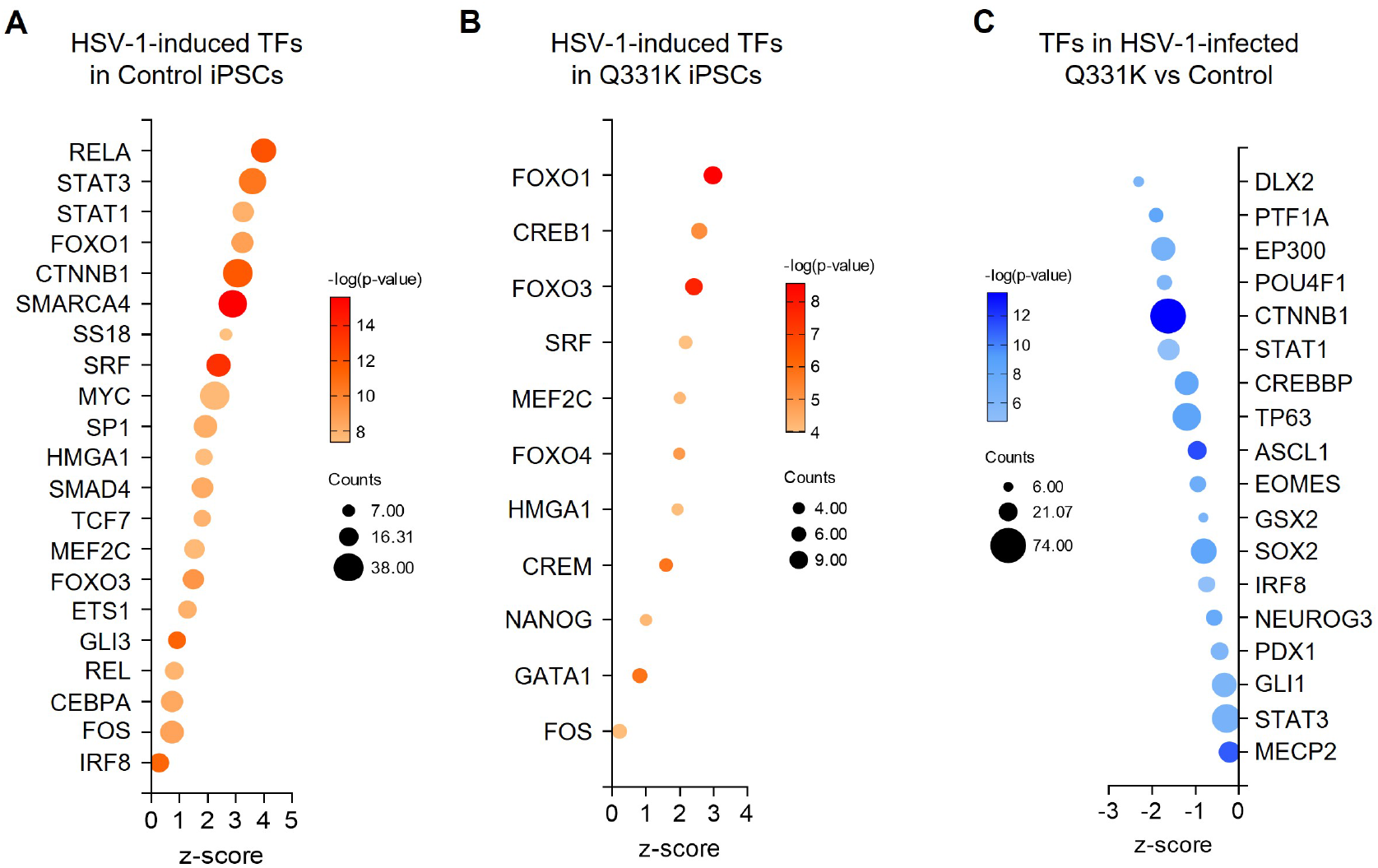
TDP-43 mutation alters predicted transcription factor engagement following HSV-1 infection. (**A**–**C**) Transcription factor predictions of DEGs in main Fig. 3G (A), Fig. 3I (B), and Fig. 3J (C). Out of the top 25 significant TFs, TFs predicted to be activated are plotted.

**Fig. S10.**
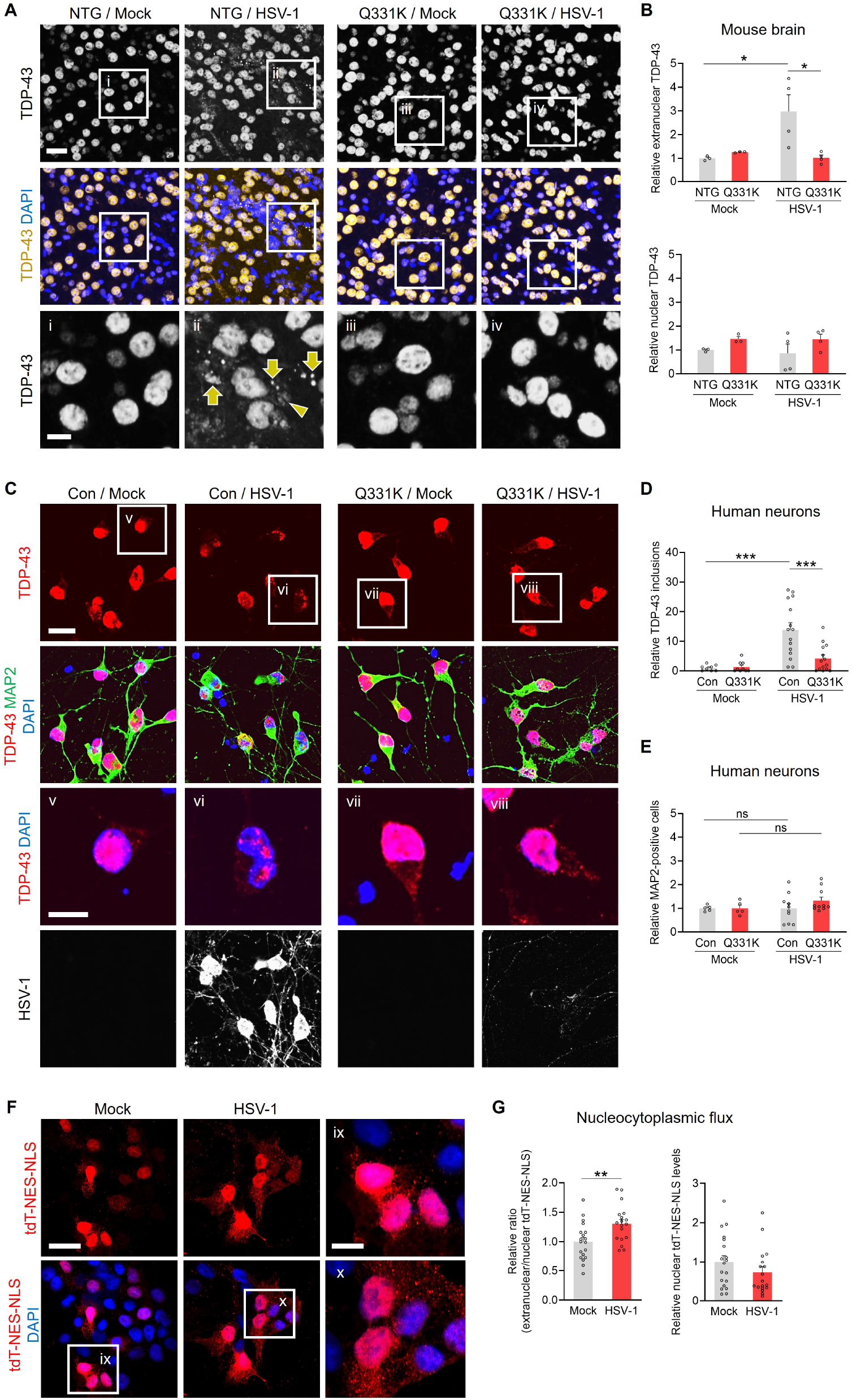
HSV-1 infection induces TDP-43 pathology in mice and human cells. **(A)** Immunolabeling for pan-TDP-43 in the thalamus of 5–7-month-old NTG and Q331K mice that received mock treatment or HSV-1 (8 DPI). Arrows point to TDP-43-positive puncta near focal infection cores in NTG mice; arrowhead highlights an example of extranuclear TDP-43 immunoreactivity. DAPI was used to visualize cell nuclei (blue). **(B)** Quantification of TDP-43 intensities in extranuclear (*top*) and nuclear (*bottom*) regions in NTG and Q331K mouse brain tissue. Normalized to NTG controls with mock infection. n = 2–4 mice per group (averages obtained from 1–3 brain sections per mouse). Two-way ANOVA (extranuclear): *F*(1, 10) = 6.942, p = 0.025 for interaction effect; Bonferroni’s test: *p < 0.05 for indicated comparisons. Scale bars: 25 µm, 10 µm (insets). **(C)** Immunolabeling for pan-TDP-43 (red) in human iPSC-derived control and Q331K neurons that received mock treatment or HSV-1 (MOI 0.01, 72 h). Neuronal marker MAP2 (microtubule-associated protein 2) is shown in green. DAPI was used to visualize cell nuclei (blue). Scale bars: 20 µm, 10 µm (insets). (**D**, **E**) Quantification of nuclear TDP-43 inclusions in MAP2-positive cells (D) and counts of MAP2- positive cells (E) in human iPSC-derived control and Q331K neurons. Averaged values across four non-overlapping areas per well. Normalized to control wells with mock infection. n = 8–15 wells per group from two differentiations (D); 5–10 wells per group from one differentiation (E). Two-way ANOVA (TDP-43 inclusions): *F*(1, 42) = 7.527, p = 0. 0089 for interaction effect; *F*(1, 42) = 18.58, p < 0.0001 for HSV-1 effect; *F*(1, 42) = 6.56, p = 0.0141 for genotype effect. Bonferroni’s test: ***p < 0.0001 for indicated comparisons. (**F**, **G**) Immunolabeling (F) and quantification (G) of tdT-NES-NLS reporter (red) in HEK293T cells that received mock treatment or HSV-1 (MOI 0.1, 48 h). G, *left*: average extranuclear-to-nuclear ratios. G, *right*: total nuclear levels of tdT-NES-NLS normalized to DAPI-positive nuclei per well and to mock-infected controls. Scale bars: 25 µm, 10 µm (insets). n = 18–19 wells per group from two independent cultures. Student’s *t* test: **p = 0.006.

**Fig. S11.**
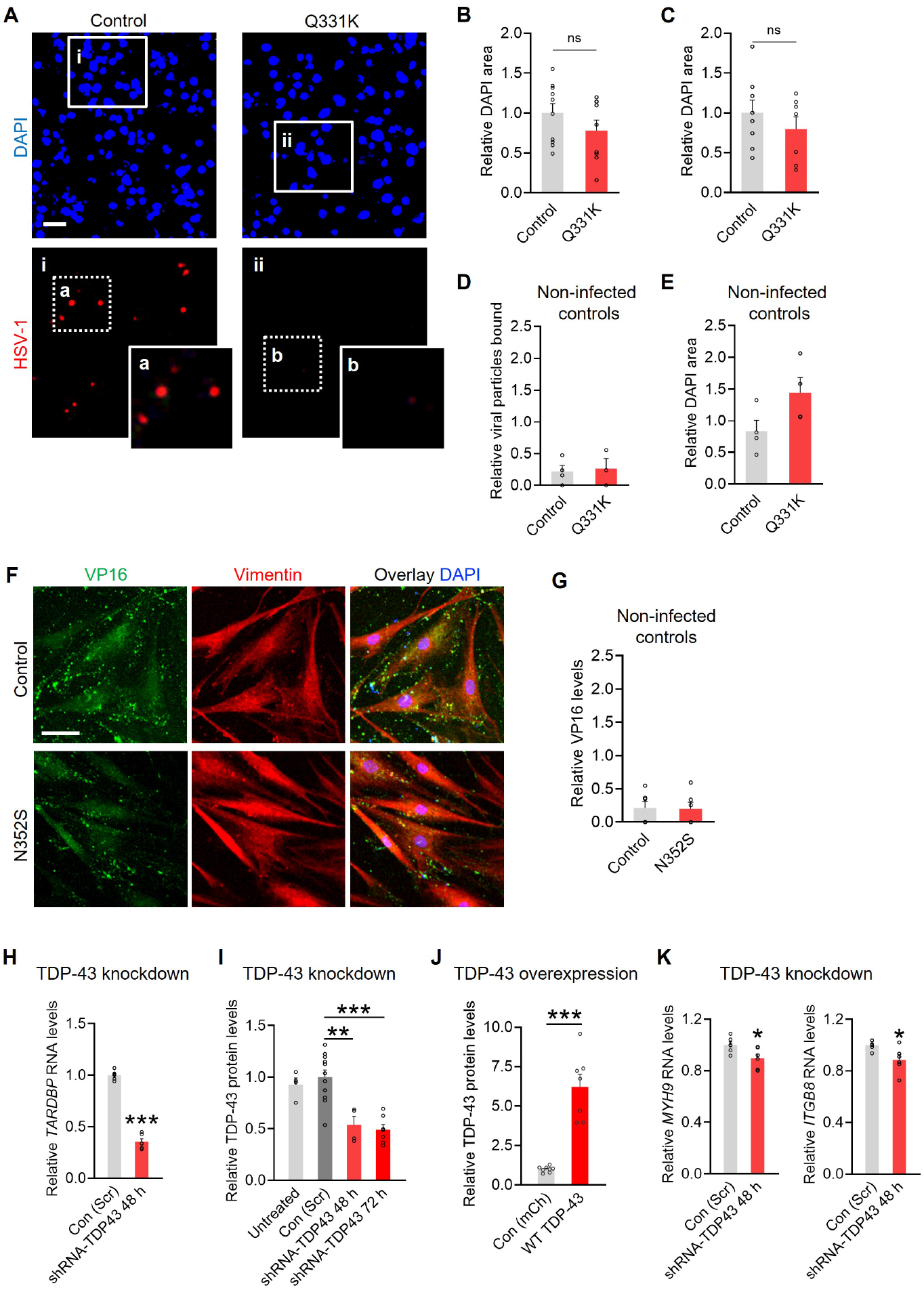
Additional analyses of viral binding and entry and TDP-43 manipulations in isolated cells. (**A**–**E**) Immunocytochemistry for bound HSV-1 particles (red). DAPI-positive nuclei are shown in blue. Relative DAPI area quantified for MOI 5 (B) and MOI 10 (C). n = 7–10 wells from two independent differentiations. Scale bar: 10 µm. (**D**, **E**) Punctate viral particles (D) and DAPI area (E) quantified in noninfected controls. Normalized to MOI 5 Control group. n = 3 wells per group. **(F)** Patient-derived N352S and control fibroblasts infected with cold McKrae strain HSV-1 at MOI 50 for 1 h on ice, followed by 30-min incubation at 37°C. Immunolabeling of VP16 particles (green) and vimentin (red). DAPI-positive nuclei in blue. Particles were normalized to vimentin area and to the control genotype. Scale bar: 30 µm. **(G)** Quantification of VP16 particles in non-infected controls. Normalized to vimentin area and the infected Control group. n = 6 wells from two independent patient lines per group. (**H**, **I**) Quantification of total human TDP-43 RNA (H) and protein (I) levels in HEK293T cells after acute transfection with plasmids encoding TDP-43-targeting shRNA (H) or control nontargeting sequence (Con, Scr). Some cells were not transfected as an additional control (Untreated). Cells were analyzed at 48 or 72 h, as indicated. RNA levels were normalized to ACTB levels per sample, and protein levels were normalized to γ-tubulin per sample, and to the respective Con (Scr) groups. N = 12 wells per condition from two independent experiments (H); n = 4–11 wells per condition from two independent experiments (I). Student’s *t* test: ***p < 0.0001 (H); One-way ANOVA: *F*(3, 22) = 13.49; p < 0.0001; Dunnett’s test: **p = 0.0011, ***p < 0.0001 (I). **(J)** Quantification of TDP-43 levels in HEK293T cells 72 h after acute transfection with plasmid encoding wild-type human TDP-43 or mCherry (mCh) control. Normalized to γ-tubulin protein levels per sample. n = 7–8 wells per condition from four independent experiments; Student’s *t* test: ***p < 0.0001; **(K)** RNA levels of indicated gene targets as measured by qRT-PCR in HEK293T cells after acute transfection with plasmids encoding TDP-43-targeting shRNA or control sequence (Scr, scrambled control) for 48 h. n = 6 wells per condition from one independent experiment; Student’s *t* test: *p < 0.05.

**Fig. S12.**
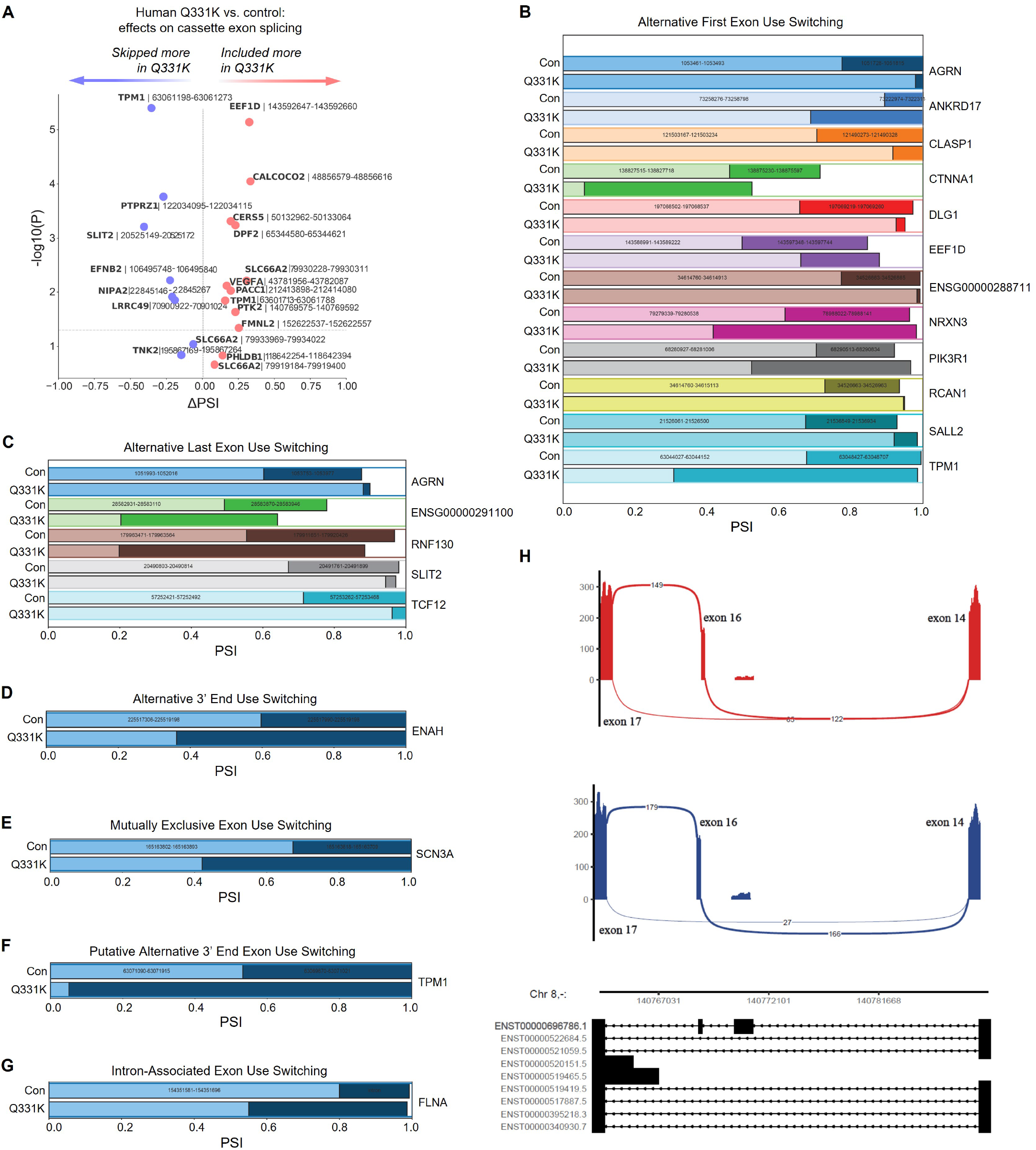
Characterization of changes in exon-level splicing in Q331K neurons. **(A)** Volcano plot showing effects on cassette exon splicing in human iPSC-derived Q331K neurons as compared to isogenic controls. Cassette exon events were extracted from MAJIQ-identified significant splicing modules that passed all three statistical criteria (*t* test, Tnom, and WILCOXON). Each dot corresponds to the junction that indicates a single cassette exon change, with gene names and genomic coordinates of the exons labeled. (**B**–**G**) MAJIQ-identified effects on junction usage associated with alternative first exon (AFE), alternative last exon (ALE), alternative 3′ splice (ALT3), putative alternative 3′ splice site (pALT3), mutually exclusive exons (MXE), and intron retention events were converted into exon-level usage profiles. For each gene, the relative usage (PSI) of the involved exons is shown for Q331K and isogenic controls. Exon usage was inferred from the usage of splice junctions connecting downstream or upstream exons to alternative exons, consistent with the junction-based framework of MAJIQ. Colored blocks represent distinct exons, with genomic coordinates indicated within each block. **(H)** Sashimi plots showing enhanced inclusion of exon 16 in Q331K neurons compared to isogenic controls. The displayed cassette-exon module corresponds to exons 14, 16, and 17 of ENST00000696786.1, with exon numbering based on this Ensembl canonical transcript in the GENCODE v44 primary assembly annotation.

**Fig. S13.**
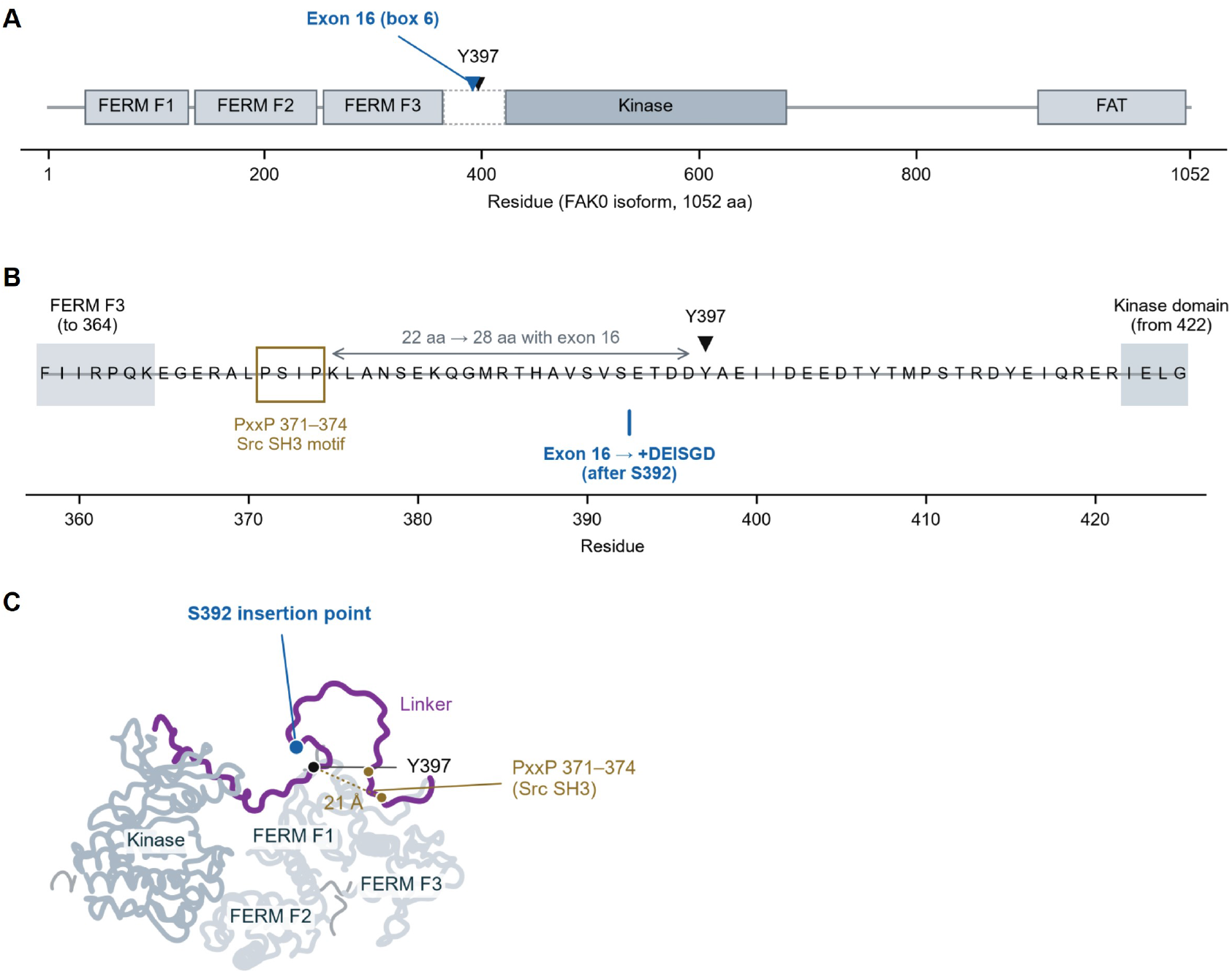
Alternative splicing of exon 16 in PTK2/FAK. Inclusion of PTK2 exon 16 within neurons carrying mutant TDP-43 is predicted to insert a six-residue sequence (box 6) into the linker region between the FERM and kinase domains in FAK protein. (**A**) Protein structure of the full-length human FAK0 isoform (1052 amino acids; UniProt Q05397). Boxes show FERM domains F1 (35–130), F2 (136–248) and F3 (254–364), kinase domain (422–680), and focal adhesion targeting (FAT) domain (912–1048). Dashed outline shows the linker region (365– 421). Blue arrowhead indicates the site of exon 16 insertion located after Ser392. (**B**) The linker sequence of FAK shown at single-residue resolution, spanning amino acids 358–425. Yellow box shows the PxxP motif at 371–374 (PSIP) that binds to Src SH3 domain. Hexapeptide DEISGD insertion is located after Ser392 (blue), lengthening the space between the PxxP motif and Y397 (which interacts with SH2 domain of Src) from 22 to 28 residues (grey double arrow). (**C**) The predicted protein structure of FAK suggests that the hexapeptide insertion is positioned within the disordered linker region. Dotted line connects Cα atoms of residues 371 and 397. Linker region conformation state and relative positions of its various sites are low-confidence predictions. AlphaFold model AF-Q05397-F1.v6, residues 31–686. Schematics were created in Claude-Science.

**Fig. S14.**
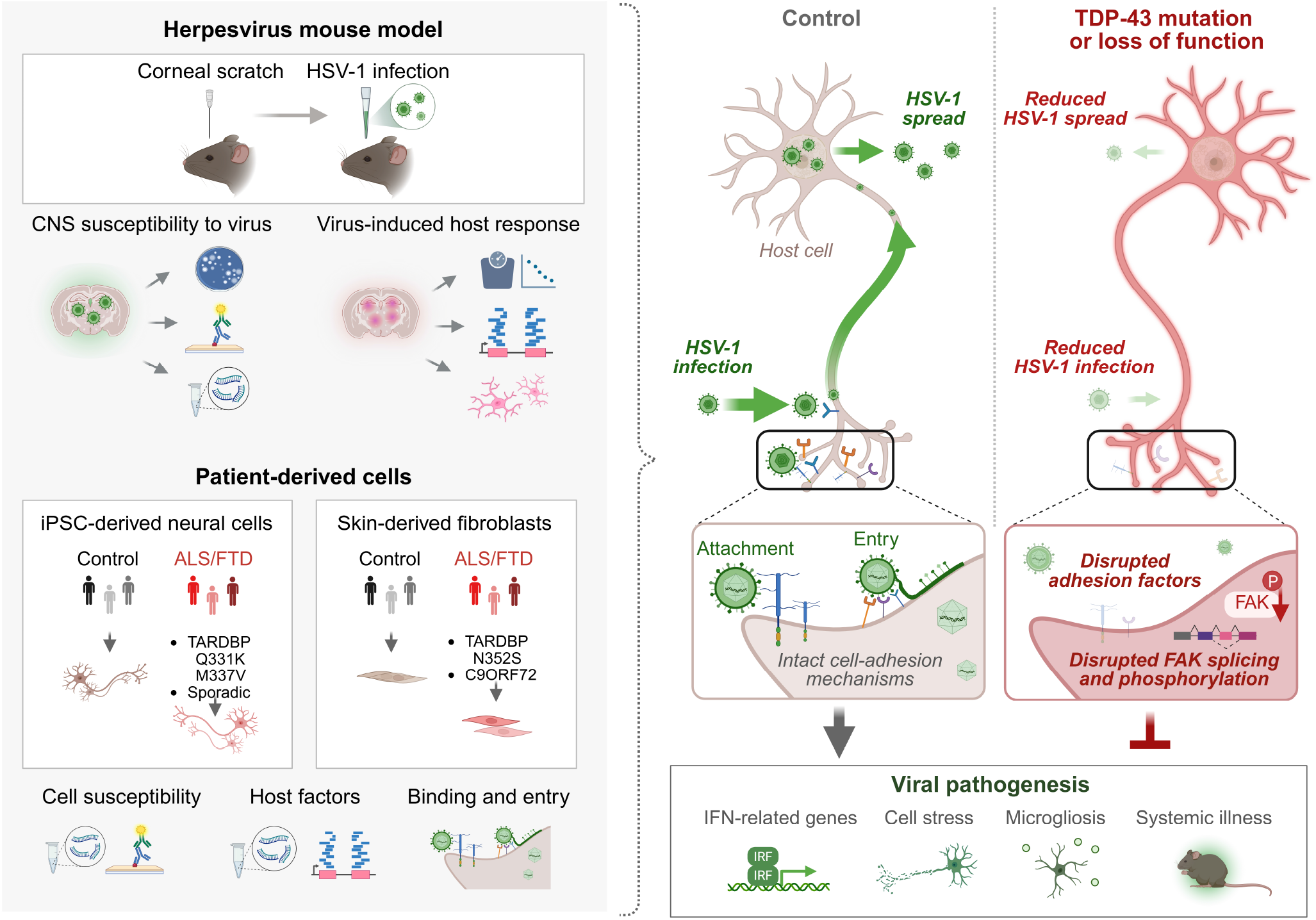
Summary of the main findings. By leveraging diverse experimental models and outcome measures, the results show that ALS/FTD-associated TDP-43 alterations confer resistance against HSV-1 infection. TDP-43 mutation or loss of function was sufficient to induce cell-intrinsic protection from HSV-1, and this protection was associated with disrupted host adhesion and entry factors, including altered RNA splicing and activity of focal adhesion kinase (FAK), which influenced viral infectivity. These findings causally link TDP-43 to viral infections through adhesion pathways and establish new roles of TDP-43 in resistance against viral pathogens.

**Table S1.** Human iPSC lines from sporadic ALS.

| **Sample ID**<br>**(Bank)** | **ALS** | **Age** | **Ethnicity** | **Race** | **Sex** | **Source** |
| --- | --- | --- | --- | --- | --- | --- |
| CS8KDJ | Sporadic | 68 | Not Hispanic or Latino | White | M | Answer ALS |
| CS9GXD | Sporadic | 68 | Not Hispanic or Latino | White | M | Answer ALS |
| CS3FGT | No | 64 | Not Hispanic or Latino | White | M | Answer ALS |

**Table S2.** Patient-derived fibroblasts.

| **Sample ID**<br>**(Bank)** | **ALS** | **Age** | **Race** | **Sex** | **Source** |
| --- | --- | --- | --- | --- | --- |
| ALS17 | N352S | Unknown | Unknown | Unknown | Gift from John Ravits |
| ALS6 | N352S | Unknown | Unknown | Unknown | Gift from John Ravits |
| Kin2ALS17 | N352S | Unknown | Unknown | Unknown | Gift from John Ravits |
| Kin3ALS6 | N352S | Unknown | Unknown | Unknown | Gift from John Ravits |
| 130923019 | C9ORF72 | 66 | White | M | Gift from Chadwick Hales |
| 130923033 | C9ORF72 | 65 | White | M | Gift from Chadwick Hales |
| 130923034 | C9ORF72 | 66 | White | M | Gift from Chadwick Hales |
| 130923006 | No | 65 | White | M | Gift from Chadwick Hales |
| CTRL10 | No | Unknown | Unknown | Unknown | Gift from John Ravits |
| OS-80033 | No | 66 | Unknown | M | Gift from Hiroshi Mitsumoto |
| OS-80036 | No | 63 | Unknown | M | Gift from Hiroshi Mitsumoto |

**Table S3.**
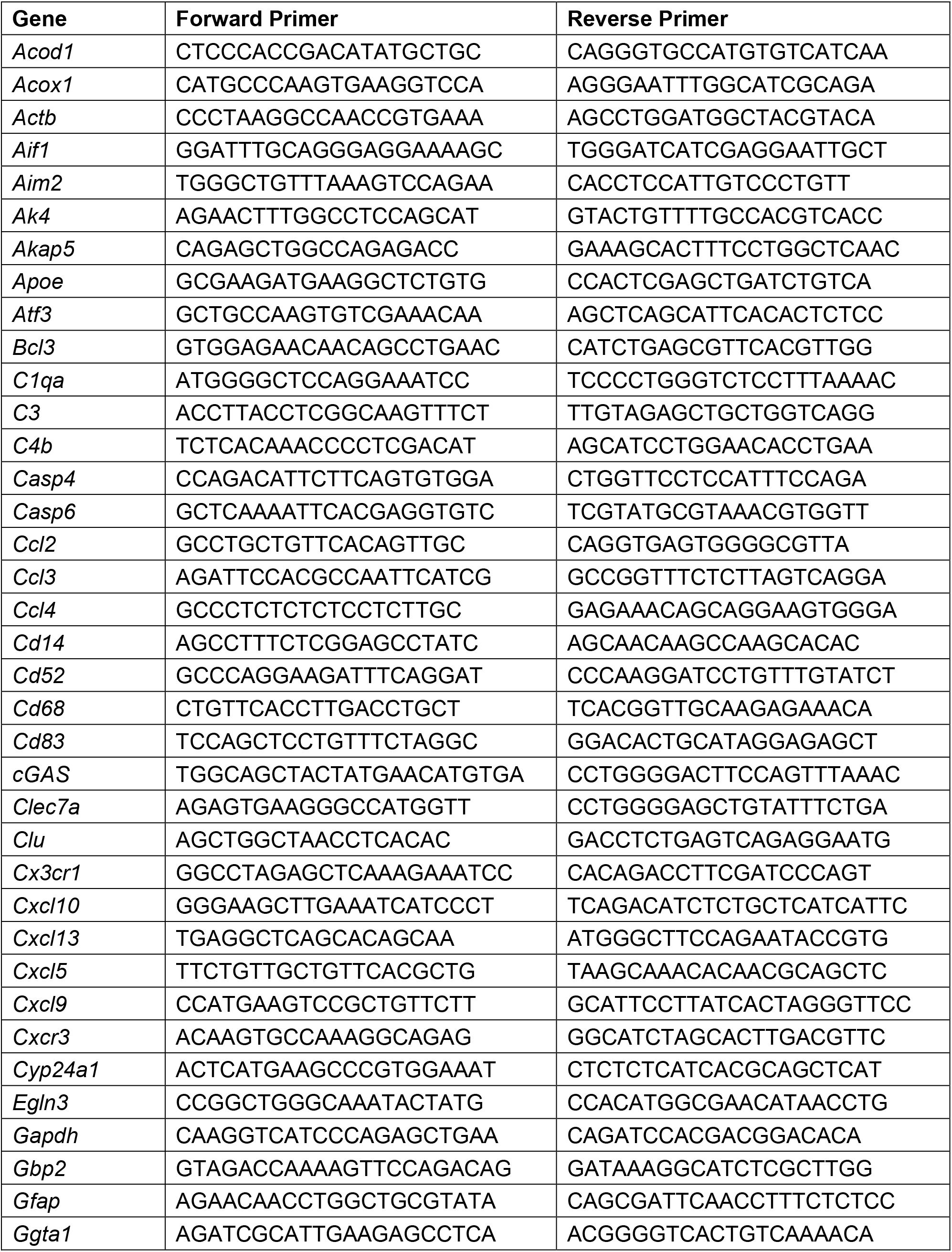
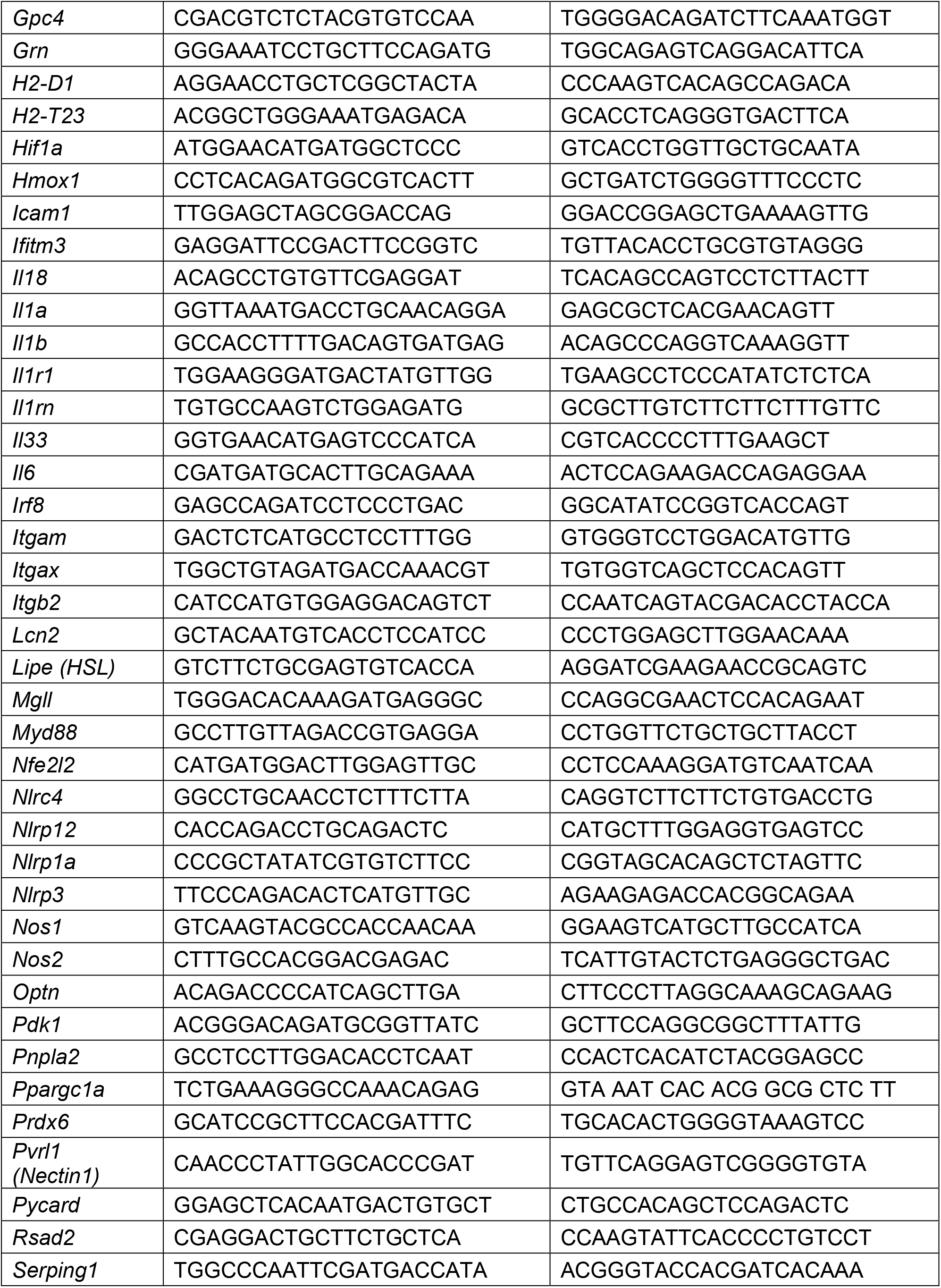
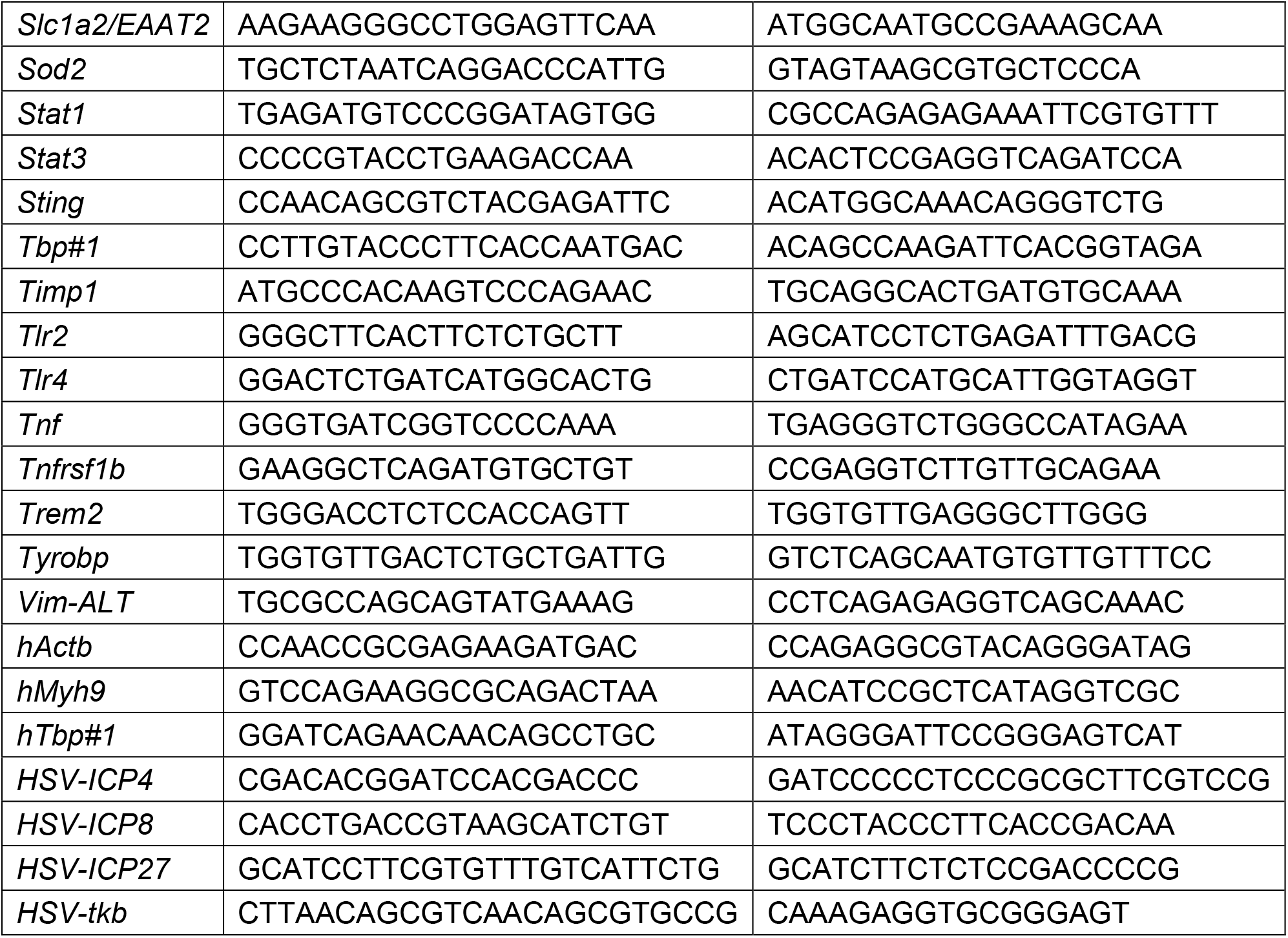
Primers used for RT-qPCR.

**Table S4.**
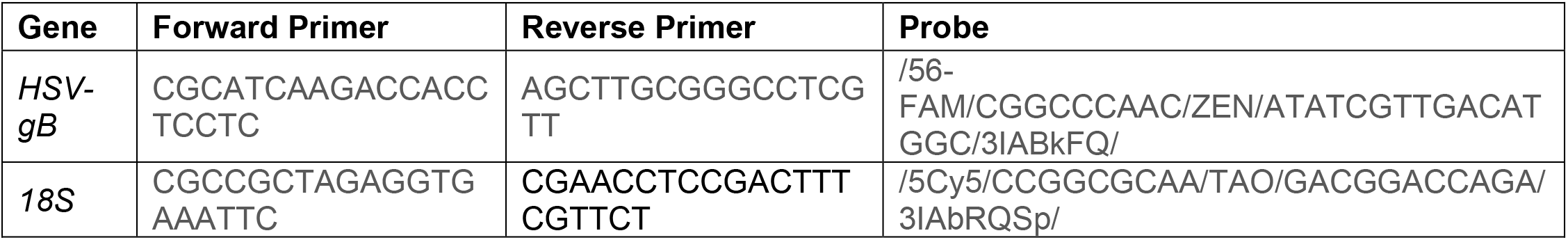
Primers and probes used for qPCR.

## Raw Western blot images

**Figure 3I.**
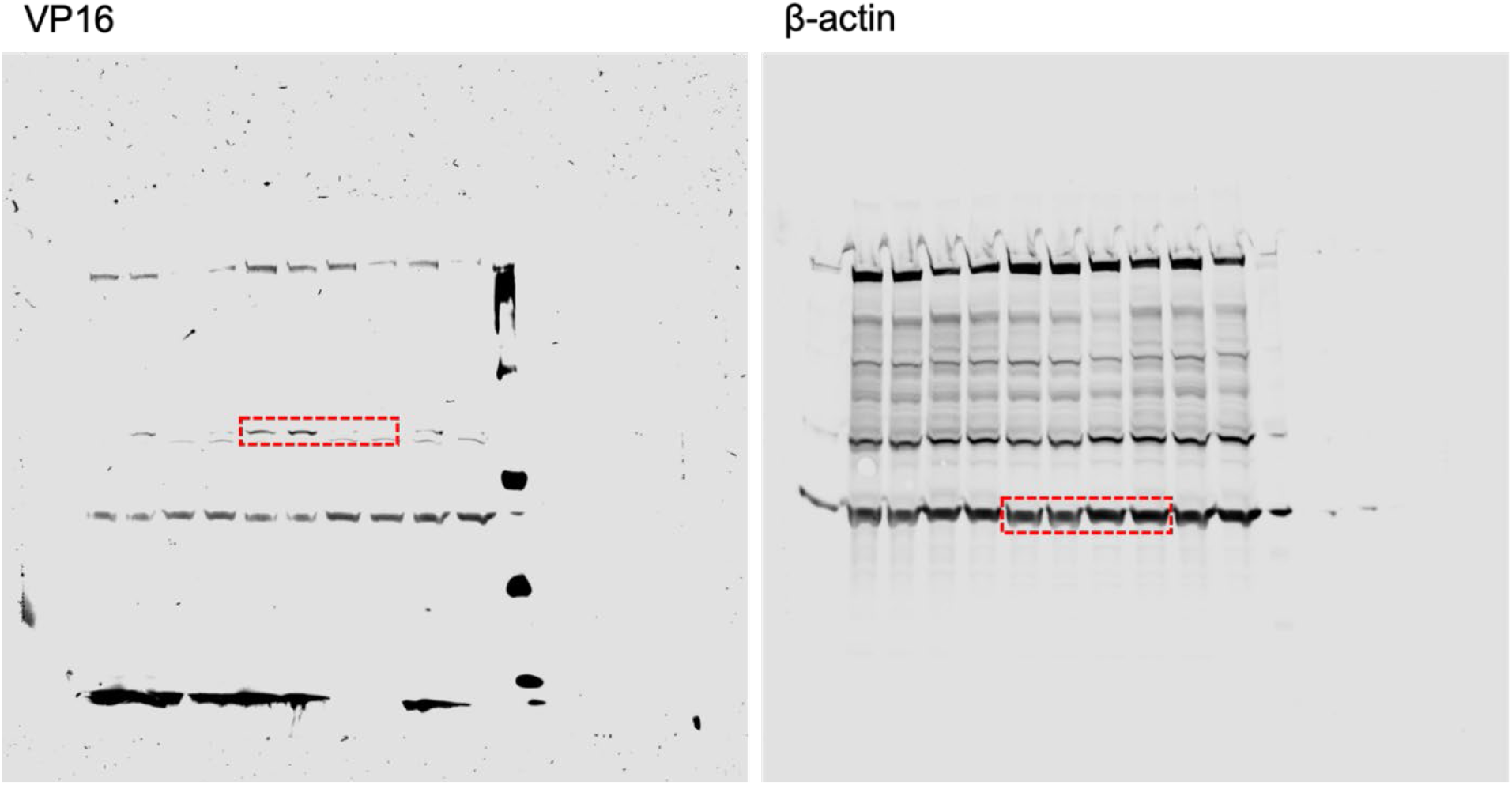

**Figure S6G.**
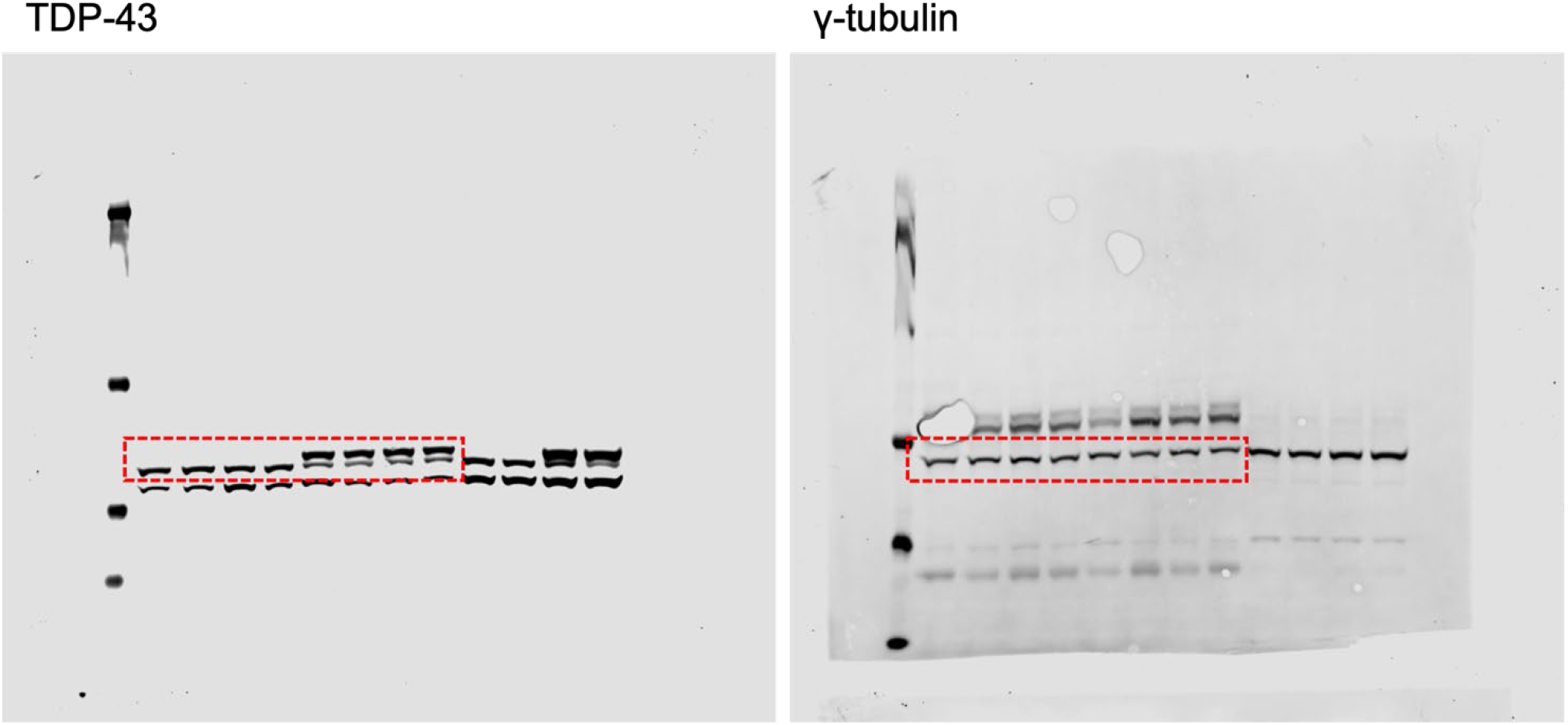

**Figure S6I.**
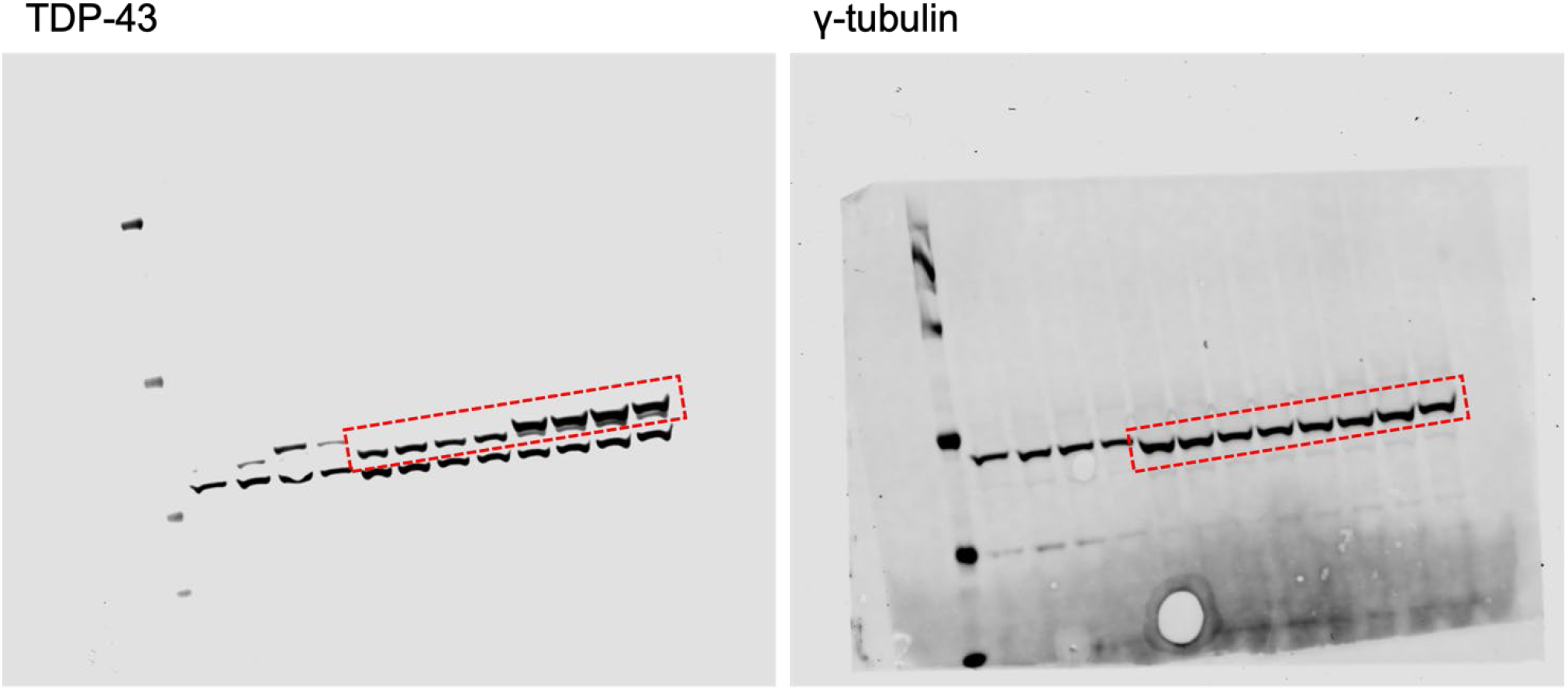

**Figure S11I.**
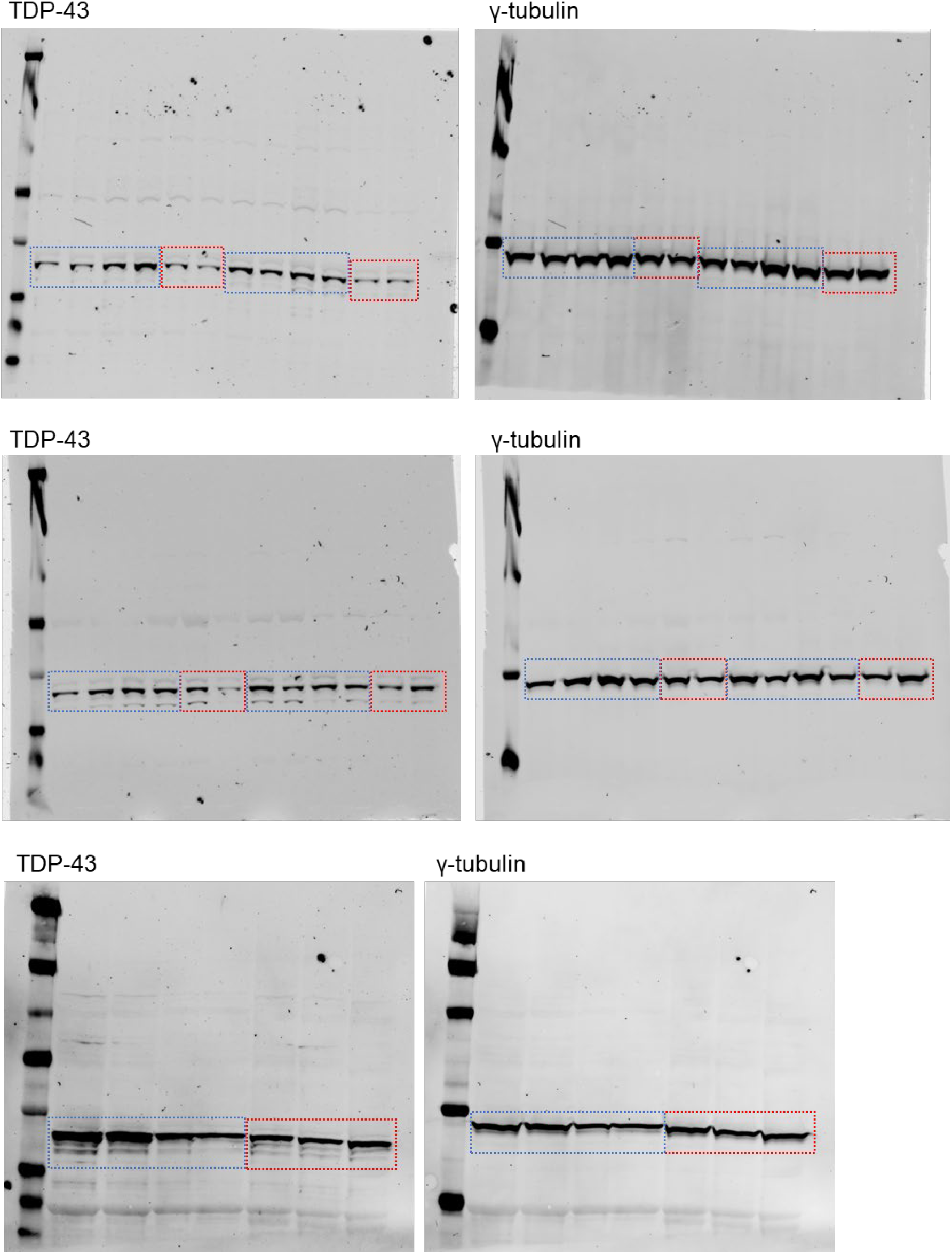

**Figure S11J.**
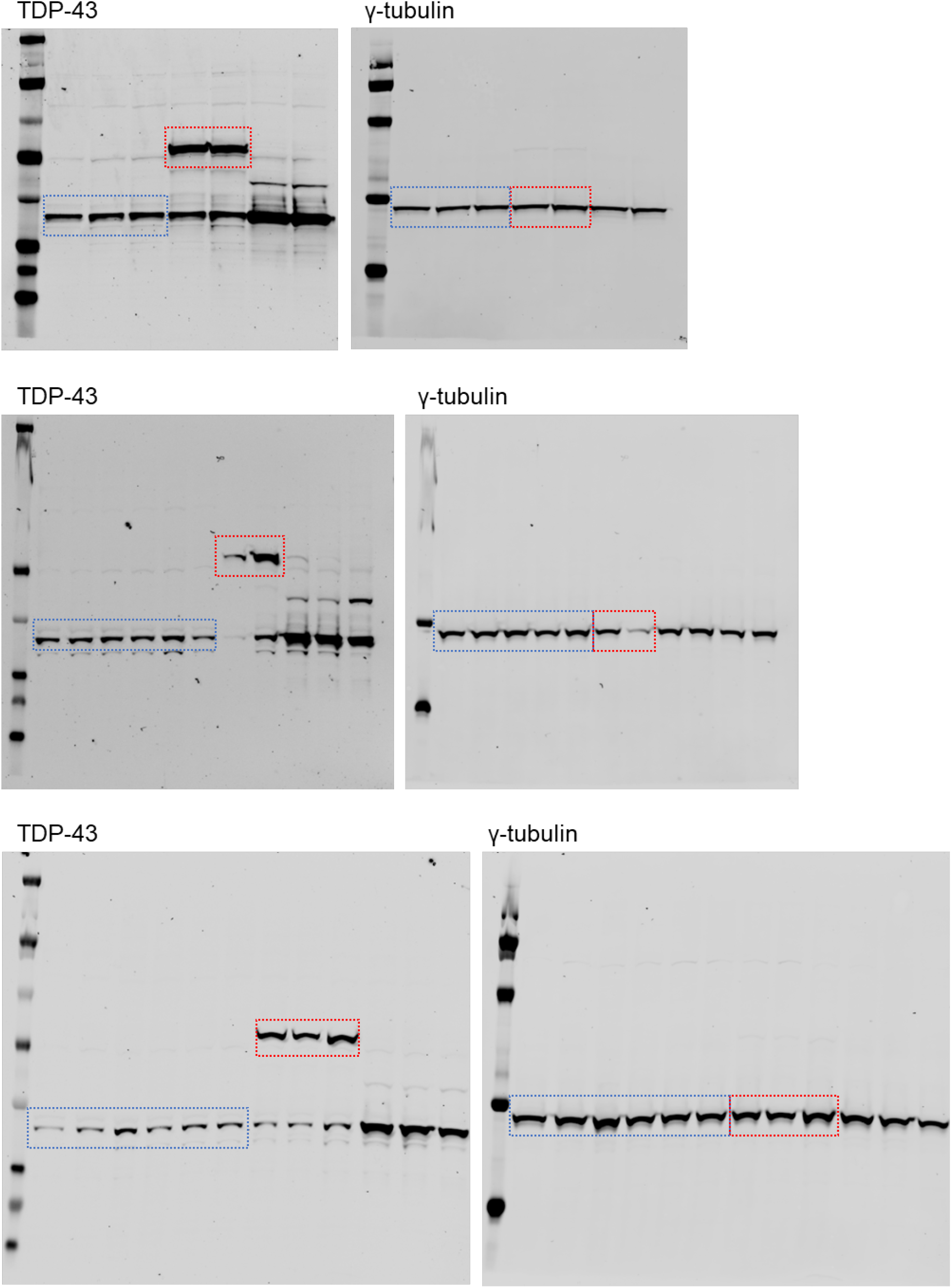

## Additional materials

**Supplemental Dataset 1**: Differential gene expression analysis of hippocampal transcriptomes in corneal scarification cohort.

**Supplemental Dataset 2**: Pathway analysis of hippocampal transcriptomes in infected versus mock- infected NTG mice.

**Supplemental Dataset 3**: Pathway analysis of hippocampal transcriptomes in infected Q331K versus infected NTG mice.

**Supplemental Dataset 4**: HSV-1-induced differential gene expression analysis of iPSC-derived neuronal transcriptomes.

**Supplemental Dataset 5**: Pathway analysis of infected versus noninfected control human iPSC- derived neuronal transcriptomes.

**Supplemental Dataset 6**: Pathway analysis of infected Q331K versus infected isogenic control human iPSC-derived neuronal transcriptomes.

**Supplemental Dataset 7**: Baseline differential gene expression analysis and pathways analysis of iPSC-derived neuronal transcriptomes.

## Notes

### Competing Interest Statement

The authors have declared no competing interest.

